# Hydrazines as versatile chemical biology probes and drug-discovery tools for cofactor-dependent enzymes

**DOI:** 10.1101/2020.06.17.154864

**Authors:** Zongtao Lin, Xie Wang, Katelyn A. Bustin, Lin He, Radu M. Suciu, Nancy Schek, Mina Ahmadi, Kai Hu, Erika J. Olson, William H. Parsons, Eric S. Witze, Paul D. Morton, Ann M. Gregus, Matthew W. Buczynski, Megan L. Matthews

## Abstract

Known chemoproteomic probes generally use warheads that tag a single type of amino acid or modified form thereof to identify cases in which its hyper-reactivity underpins function. Much important biochemistry derives from electron-poor enzyme cofactors, transient intermediates and chemically-labile regulatory modifications, but probes for such species are underdeveloped. Here, we have innovated a versatile class of chemoproteomic probes for this less charted hemisphere of the proteome by using hydrazine as the common chemical warhead. Its electron-rich nature allows it to react by both polar and radicaloid mechanisms and to target multiple, pharmacologically important functional classes of enzymes bearing diverse organic and inorganic cofactors. Probe attachment can be blocked by active-site-directed inhibitors, and elaboration of the warhead supports connection of a target to a lead compound. The capacity of substituted hydrazines to profile, discover and inhibit diverse cofactor-dependent enzymes enables cell and tissue imaging and makes this platform useful for enzyme and drug discovery.

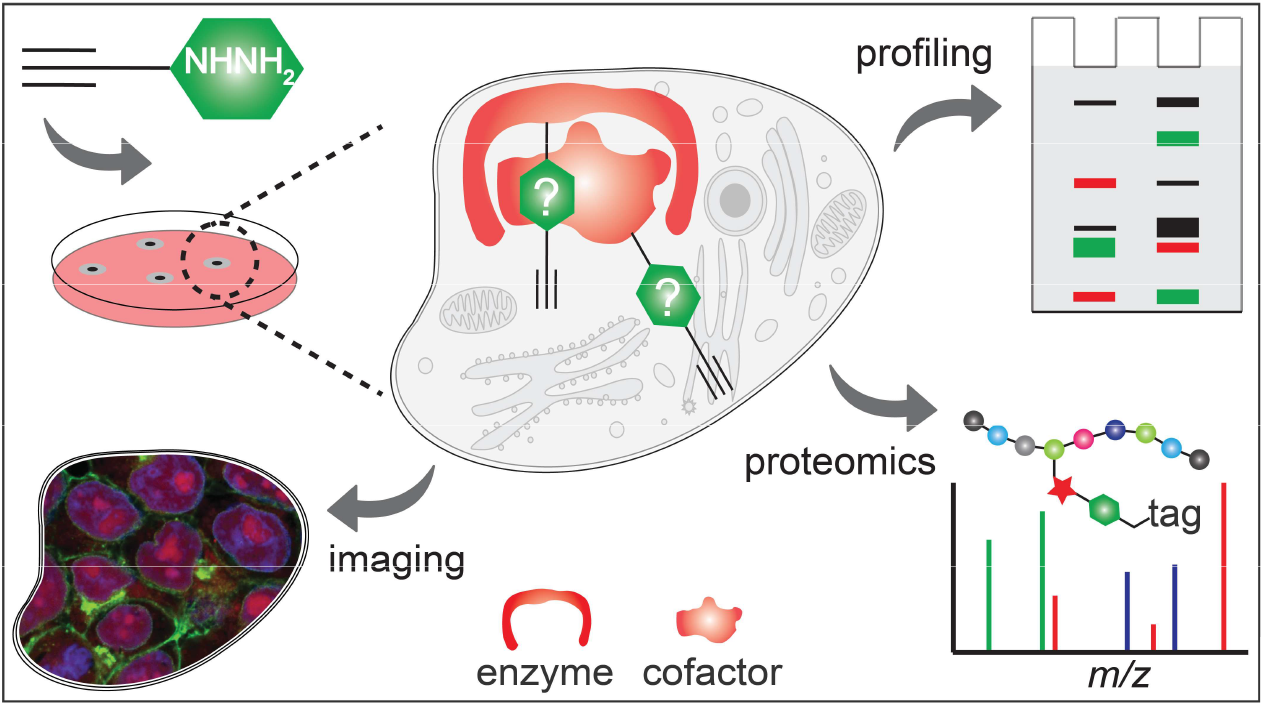

## MAIN

Chemical biology probes for protein profiling have been used to discover enzymes, inhibitors and drugs^1–3^. Many of these probes are, by design, agnostic to molecular recognition by specific active sites but leverage chemically selective warheads that covalently couple to only one of the 20 amino acids^4^ or a modified form thereof^5^. This tagging of functionally hyper-reactive residues often identifies sites of catalysis in enzymes^6^. For such classical activity-based approaches^7^, covalent coupling to the reactive residue that defines an enzyme family or functional class can be used to read out (profile) that type of activity across the proteome, without prior knowledge of the individual substrates or transformations and in the absence of any biased molecular recognition. Because catalytic residues within an enzyme family are often readily predicted by genetics and bioinformatics, probes can be intentionally designed according to the known chemical selectivity of the warhead. In more recent extensions of this approach, ligand-based probes have been used to target hotspots of functionally diverse binding sites across the proteome^8^. Here, amino acid-selective warheads (for cysteine^9^, lysine^10^, aspartate^11^ or tyrosine^12^) or photoreactive handles^13,14^ have been applied to recognize and tag modestly activated amino acids in solvent-accessible binding pockets, including active and allosteric sites of enzymes, receptors and other types of druggable proteins. These approaches enable i) discovery of protein activities or ligandability on the basis of enhanced reactivity with a specific probe and ii) subsequent identification of target-selective small molecules by chemical elaboration of the attacking warhead or by screening for potent inhibitors of attachment of the original, nondescript probe.

Functional reactivity of the proteome extends beyond reactive amino acids, yet chemoproteomic platforms to discover, profile and modulate such functionality are underdeveloped. Cofactor-dependent enzymes, transient intermediates and other chemically labile protein modifications engage in catalytic and regulatory functions that are vital for virtually every cellular process, as they participate in challenging and diverse chemical transformations and sophisticated signal-transduction pathways that are conserved across biology^15,16^. A large fraction of enzyme cofactors are electron-deficient in their resting or intermediate states. A hypothetical chemoproteomic platform to globally target cofactor-harboring enzyme active sites would require an electron-rich warhead that would be agnostic to both active site structure and functional group identity, recognizing only this general electron deficiency. A probe based on such a generalized warhead would have the capacity (i) to target all protein classes that utilize electron-deficient moieties for physiological function and (ii) to globally capture such species while operating in native biological systems across diverse physiological contexts.

Hydrazine could potentially represent such a broad-spectrum covalent warhead for a chemoproteomic platform to capture cofactor-dependent enzymes. Its electron-rich nature makes it both nucleophilic and reducing, allowing for broad reactivity toward electron-deficient groups. In a previous study^17^, hydrazine was used as warhead for reverse-polarity (RP) activity-based protein profiling (ABPP). Initially expecting the warhead to act only as a nucleophile to target polar electrophiles such as carbonyl, ester and imine groups, we identified unexpected targets not known to harbor such electrophiles. Here, we show that the previously unexplained targeting of these proteins resulted from the reactivity of hydrazines also toward radicaloid oxidants. We show that this property confers unprecedented versality for generalized probing of both stable and transient electron deficient moieties in proteins. The hydrazine probes broadly capture enzymes across multiple different functional classes by distinct chemical mechanisms, but retain the active site targeting and dependence on functional state that have made conventional ABPP probes such powerful tools for inhibitor and drug discovery. By targeting reactive groups acquired post-translationally or transiently in the act of catalysis, our probes enable discovery of functionally relevant but not readily genetically predictable reactive moieties in enzymes. As such, they represent unusually versatile tools for identification of inhibitors, drugs and even new targets.

## RESULTS

### Global map of hydrazine-reactive proteome

Alkyl probe **1** and aryl probe **2** (Fig. 1a) were previously used to discover protein electrophiles^17^. However, we envisioned dual profiling platforms of both electrophilic and oxidizing functionality in the proteome, potentially operating as activity-based and mechanism-based inactivators (MBI) of known and unknown oxidizing centers in proteins^18,19^. We synthesized a “clickable” hydrazine probe, **3**, which is chemically similar to probes used in the previous work but more closely resembles the known drug phenelzine (alkylaryl hydrazine **4** in Fig. 1a). Phenelzine is the best studied hydrazine-bearing drug. It functions by irreversibly inhibiting monoamine oxidase (MAO)^18,19^, the neurotransmitter-metabolizing enzyme, by an O_2_-dependent radical mechanism that results in alkylation of the covalently bound flavin adenine dinucleotide (FAD) cofactor in the active site. It was an early therapy for depression and one of the first examples of oxidative activation of a hydrazine by an enzyme. Notably, it penetrates the brain, suggesting that, in general, substituted hydrazines might access even the most isolated tissue space.

**Fig. 1.**
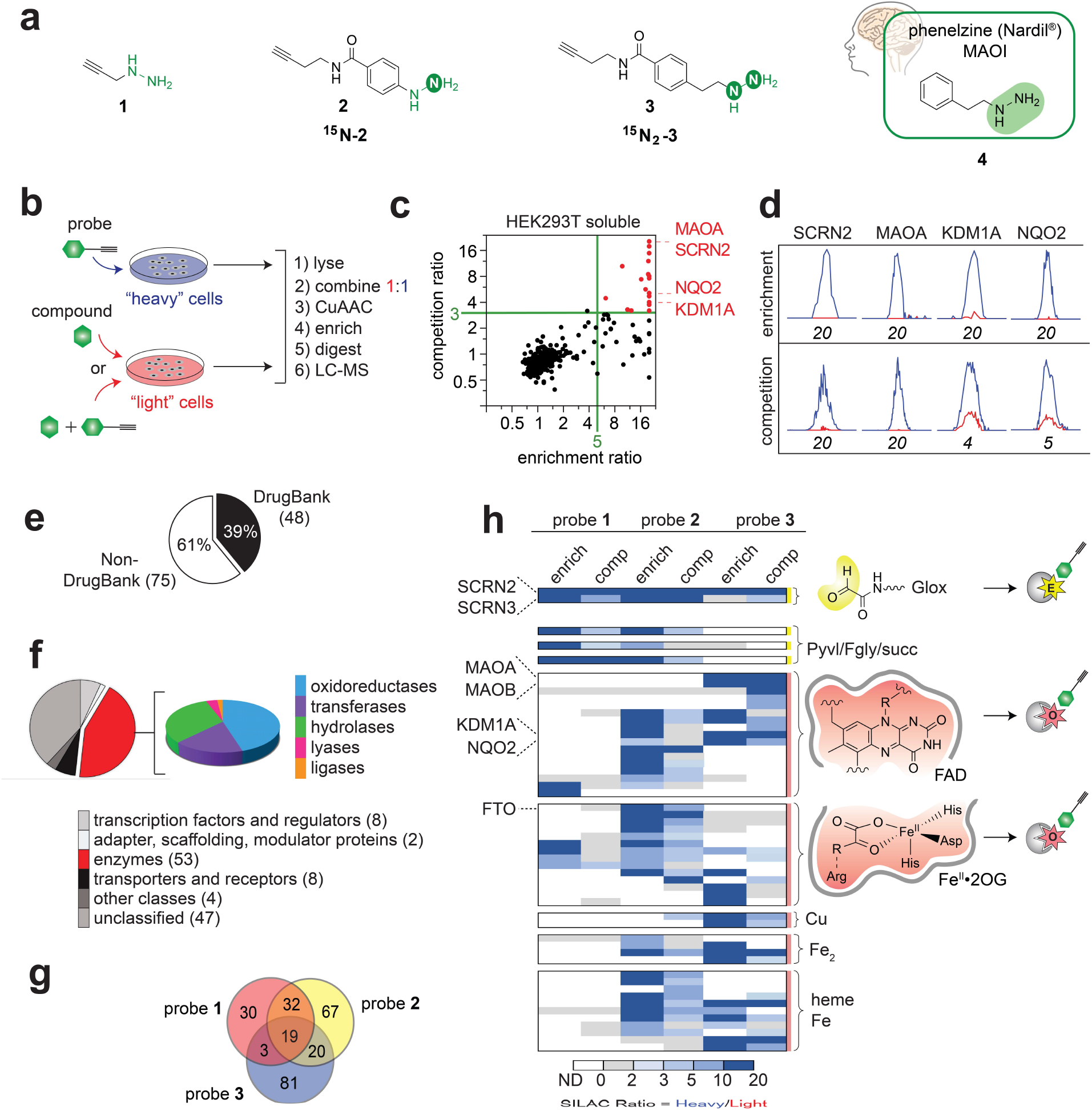
Hydrazine pharmacophores broadly capture enzymes across multiple functional classes in human cells. **a**, Structures of hydrazine probes (alkyl probe **1**, aryl probe **2** and alkylaryl probe **3** with corresponding isotopologues **^15^N**-**2** and ^**15**^**NN_2_**-**3**) and CNS drug phenelzine **4** (non-clickable analogue of **3**). Green designates hydrazine warheads. **b**, Schematic for MS^2^-based quantitative proteomics experiments (enrichment and competition) using stable isotopic labelling by amino acids in cell culture (SILAC)^21^. Isotopically heavy and light cells, proteomes, and peptides are depicted in blue and red, respectively. **c**, Quadrant plot of average competition versus enrichment protein ratios for probe **3** from quantitative proteomic analysis of the soluble proteome of HEK293T cells. Highest-reactivity targets (*upper right* quadrant) found in both experiments and labelled by probe **3** are highlighted in red. **d**, Extracted MS1 chromatograms and corresponding heavy/light ratios (enrichment and competition) for representative tryptic peptides of endogenous forms of highest-reactivity targets. **e**, Fraction of **3**-labelled proteins from HEK293T and MDA-MB-231 cells found in DrugBank and **f**, their known or predicted functional classes. **g**, Venn diagram of shared and unique protein targets of probes **1**-**3** from both cell lines. **h**, Heatmap showing relative protein enrichment and competition indices for targets known or predicted to harbor electron-deficient moieties. Targets are grouped on the basis of type of electron-deficient group, electrophilic (yellow) or oxidizing (red), and electron-deficient cofactor class. Structures for three diverse classes (Glox, FAD and Fe/2OG) are shown.

Gel-based proteomic profiles of **3** were generated by *in situ* treatment (0.5 h, 37 °C) of HEK293T and MDA-MB-231 cells, followed by coupling of probe-captured proteins to an azide-rhodamine (Rh-N_3_) reporter tag using copper(I)-catalyzed azide-alkyne cycloaddition (CuAAC or “click” chemistry)^20^ and visualization by SDS–PAGE with in-gel fluorescence scanning^1,17^. Proteomic reactivity of **3** was both time- and dose-dependent, and labelling was blocked by pretreatment (15 min, 37 °C) with **4** (Supplementary Fig. 1). To identify and quantify protein targets of probe **3***in situ*, proteomic extracts from **3**-treated cells were fractionated, conjugated by CuACC chemistry to biotin-N_3_, enriched by adsorption to streptavidin resin and analyzed by mass spectrometry (MS)-based proteomics (Fig. 1b and Supplementary Fig. 2)^1,17^. To quantify high-reactivity targets, proteomes from isotopically-labelled^21^ **3**-treated cells (1 mM, 0.5 h) were ratiometrically compared against proteomes obtained by cells subjected to two types of control treatments: to assess enrichment, cells were treated with **4** under the same conditions as **3**; to assess competition, cells were pretreated with 10-fold excess (10 mM, 15 min) of **4** relative to **3**. This latter experiment affords estimates from the purely ratiometric quantitation of both the relative abundance of a given target and how quantitively it is captured by the probe. Protein ratios were calculated from median peptide ratios from at least three unique quantified peptides per protein per experiment. Average competition ratios were plotted against average enrichment ratios (*n* ≥ 7 biological replicates; Supplementary Dataset 1) for proteins quantified in both types of experiments (Fig. 1c, Supplementary Dataset 1, and Supplementary Fig. 3). Proteins that were (i) substantially enriched by treatment with **3**, coupling to biotin and absorption on streptavidin resin (ratio ≥ 5) and (ii) depleted by prior treatment with **4** (ratio ≥ 3) in the *upper right* quadrant were taken to be high-reactivity targets (30 total proteins from the soluble and membrane fractions of the HEK293T and MDA-MB-231 cell lines). Representative peptide ratios from enrichment and competition experiments for four targets are shown in Fig. 1d. Meta-analysis of expanded **3**-labelled proteins (enrichment ratio ≥ 5 or competition ratio ≥ 3) showed that, despite the facts that the competitor is an actual clinical drug and the probe is an analog of one, the majority of targets are not found in the DrugBank (61%, Fig. 1e). A large fraction of targets are known or predicted by genetics to have catalytic activity (Fig. 1f, *left*). Many members of this subset are known or predicted oxidoreductases (*right*).

### Capturing enzymes across multiple functional classes

As probe **3** (alkylaryl) is structurally-related to **1** (alkyl) and **2** (aryl)^17^, we aimed to understand how the target profiles compare and whether the hydrazine substituent confers selectivity. Profiles were aligned (Supplementary Dataset 2), and targets with enrichment ratios ≥ 5 or competition ratios ≥ 3 for one or more of the probes in either cell line were compared in a Venn diagram (Fig. 1g). Alignment of targets according to known or predicted cofactor usage (e.g. flavin, heme) or post-translational modification generated an activity-based heatmap (Fig. 1h and Supplementary Table 1). All three probes showed preference for SCRN3 – an uncharacterized enzyme that previously provided the first example of a naturally occurring *N*-terminal glyoxylyl (Glox) electrophile. **1** and **2** both targeted enzymes with functional electrophilic species such as pyruvoyl (Pyvl), formylglycyl (Fgly) or succinimide (Succ)^17^, whereas **2** and **3** targeted only partly overlapping sets of enzymes in other functional classes, including those defined by the use of oxidizing cofactors such as FAD, iron (Fe) or copper (Cu). **3** selectively targets MAOA and MAOB, consistent with its covalent capture of covalently-bound FAD according to the known mechanism of phenelzine (**4**)^18^.

The data illuminate the potential for hydrazine to serve as general mechanism-based warhead for enzyme targets well beyond MAO. Notably, other flavoproteins with noncovalently-bound FAD cofactors (e.g. lysine-specific histone demethylase 1A (KDM1A) and ribosyldihydronicotinamide dehydrogenase [quinone] (NQO2)) are also highest-reactivity targets of **3**, implying that it can directly covalently couple to the enzymes themselves in addition to the FAD cofactor. Iron(II)- and 2-oxoglutarate-dependent (Fe/2OG) enzymes – including fat mass and obesity-associated dioxygenase (FTO), the O_2_-dependent RNA demethylase that is critical for an array of cellular processes and disease states, including CNS function^22^, obesity^23^ and cancer^24^ – were also identified as specific probe targets, with **2** being more selective than **3** toward these enzymes. To the best of our knowledge, this represents the first report of the targeting of an Fe/2OG enzyme with a covalent inactivator.

### Activity-based probes for three unexpected classes

The broad reactivity of the hydrazine probes could represent either an asset or a liability: the question turns on whether their capture of the diverse enzyme targets remains active-site-directed and dependent on functional state, the two crucial traits that have made classical ABPP probes such useful tools in inhibitor and drug discovery. This issue was investigated by gel-based profiling of enzymes from four diverse structural and mechanistic classes: (i) covalently-bound FAD-dependent monoaminergic amine oxidases (MAOA, MAOB) as a control, (ii) noncovalent flavoenzymes (epigenetic eraser KDM1A and detoxifying quinone reductase NQO2), (iii) functionally uncharacterized SCRNs (SCRN1-3) and (iv) FTO. Overexpressed wild-type proteins from probe-treated HEK293T cells retained the hydrazine reactivity and probe preference observed by proteomics analysis of their endogenous forms (Fig. 2a-c). In all cases, labelling was abolished by substitution of active site residues that are essential for activity and/or productive cofactor binding. These observations confirm that capture by the probes requires the enzymes to be active (Fig. 2a-c)^18,25–27^.

**Fig. 2.**
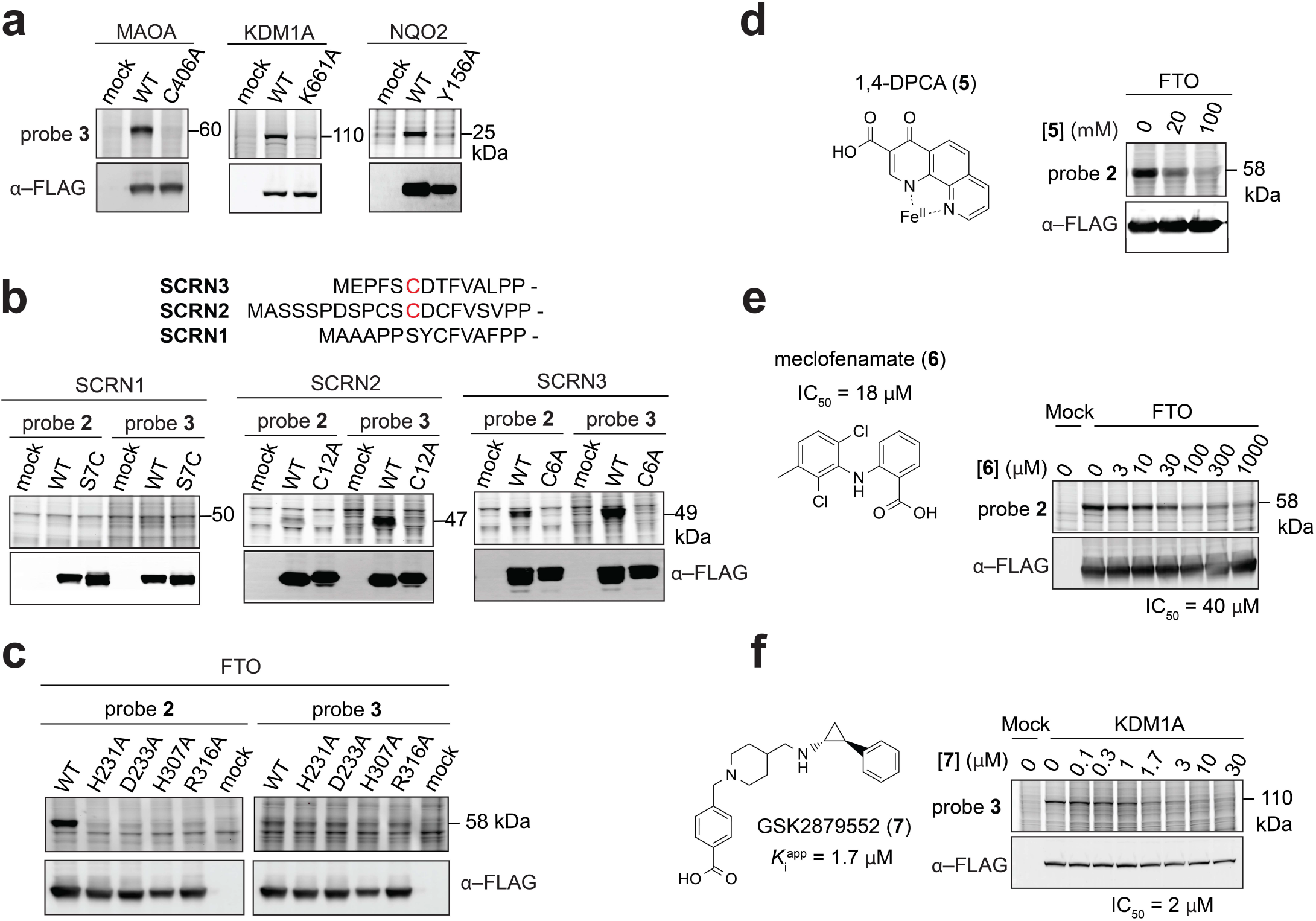
Covalent labelling is dependent on functional state and blocked by active-site-directed inhibitors. Probe labelling (*upper*) and expression profiles (*lower*) for probe-treated cells expressing the indicated protein target or inactive variant thereof. Transfection with the appropriate empty expression vector (‘mock’) is used as a control. Molecular weights are indicated. **a**, **3**-labelling of wild-type MAOA, KDM1A and NQO2 flavoenzymes but not catalytically inactive variants. **b**, Sequence alignment of SCRN proproteins (*upper*), cleavage site, and Cys residue (red) predicted to be catalytic, but shown in SCRN3 to be post-translationally converted to Glox. **2-** vs**. 3**-labelling (*lower*) profiles of SCRN1-3 compared to indicated variants. **c**, **2-** vs**. 3**-labelling of wild-type FTO but not catalytically inactive variants. Dose-dependent inhibition of FTO-labeling by **2** with **d**, iron chelator drug, 1,4-DPCA (**5**) and **e**, meclofenamate (**6**), a cyclooxygenase-2 (COX2) inhibitor also known to inhibit FTO. **f**, Dose-dependent inhibition of KDM1A-labelling by **3** with its covalent inhibitor GSK2879552 (**7**).

Our identification of SCRN2 as a probe-reactive target suggests, on the basis of its analogy to the previously characterized SCRN3, that it also reacts with the hydrazine warhead through a Cys-derived *N*-terminal Glox and/or Pyvl group. Probe labelling was abolished by replacement of cysteine 12, the residue predicted to support its hypothetical hydrolase activity. This residue also aligns with the Cys6 of SCRN3 (Fig. 2b) that is transformed to Glox. Probes deployed in the prior study^17^ could not differentiate between SCRN2 and SCRN3 isoforms, and only for SCRN3 was the distribution of *N*-terminally distinct forms determined. Here, SCRN2 ratios (R_enrich/comp_ ≥ 20) were consistently higher than for SCRN3 (R_enrich/comp_ ≤ 4) in a given cell line, suggesting that probe **3** might be selective for isoform 2 (Fig. 1h and Supplementary Dataset 1). In gel-labelling profiles for overexpressed SCRN2 and SCRN3, probe **3** labelled both isoforms more intensely than probe **2** (Fig. 2b). By contrast, no hydrazine reactivity was observed for SCRN1, which has a serine rather than cysteine at the position that, in isoform 3 (and, as shown below, also isoform 2), becomes Glox. Replacement of the SCRN1 Ser with Cys did not result in acquisition of reactivity toward the probes, suggesting that additional factors are important in determining whether Glox is installed (Fig. 2b). The data suggest that **3** may be a valuable tool for investigating endogenous SCRN2/3 activity and that that even subtle modification to the probe can engender selectivity not only between functional enzyme classes but also between isoforms within the same class.

Similarly, FTO is a selective target of **2**^17^ but not **3** (Fig. 1h and 2c), and its labelling was abolished by substitution of residues that coordinate the cofactor (His231, His307 and Asp233) or make key contacts with the 2OG cosubstrate (Arg361)^28,29^ (Fig. 2c). In addition, pretreating FTO-overexpressing HEK293T cells with the prolyl hydroxylase domain (PHD) inhibitor 1,4-DPCA (**5**), an iron chelator known for its anticancer^30^ and regenerative activities^31^, blocked probe labelling in a dose-dependent manner, suggesting that, in cells, the labelling event is both active-site-directed and dependent on the presence of the cofactor (Fig. 2d). Meclofenamate (**6**), a non-steroidal, anti-inflammatory (NSAID) drug known to inhibit FTO^32^, also blocked probe labelling, and the dependence of signal abrogation on inhibitor concentration was consistent with the IC_50_ for binding of the drug (Fig. 2e). Likewise, GSK2879552 (**7**), a cyclopropylamine-containing KDM1A inhibitor with anticancer activity, also blocked probe labelling with an IC_50_ that is comparable to the published *K*_i_^app^ for its target^33,34^ (Fig. 2f). Probe blocking by other covalent and noncovalent inhibitors is analyzed in Supplementary Fig. 4. The direct blockade of functional enzymes suggests a promising route for inhibitor screening in drug discovery.

### Mapping sites and mechanisms of covalent inactivation by substituted hydrazines in living cells

Phenelzine and related monoamine oxidase inhibitors (MAOIs) covalently target N5 of the FAD cofactor^18^. By the same mechanism, related small molecules inhibit other enzymes with noncovalent FAD cofactors (e.g. NQO2^35^ and KDM1A^36^). Reaction of **3** by an analogous mechanism (Fig.3a, *upper*) would not, in our experiments, lead to enrichment of an enzyme bearing a noncovalently bound flavin, because covalency with the protein is required for retention on the resin and detection by SDS-PAGE. Nevertheless, formation of reactive radicals, first on the nitrogen of the hydrazine warhead and subsequently, by radical elimination of N_2_ on carbon (R• in Fig. 3a), could potentially result in covalent coupling directly to the protein in the vicinity of the active site (Fig.3a, *lower*). To understand the basis of the previously reported inhibition of KDM1A by phenelzine and our observation here that NQO2 is also targeted by the phenelzine-mimicking probe, we identified the small molecule and protein products of the labelling reactions. Following reaction of either **3** or **4** with reconstituted NQO2 overexpressed in and purified from *E. coli*, FAD adducts were readily detected by LC-MS. The detected species had parent masses implying loss of the N_2_H_4_ fragment from the probe during the coupling reactions (Supplementary Fig. 5). To confirm that radical coupling with the protein can also occur and that this event explains targeting of the enzymes by the probes (Fig. 2a), we determined sites of labelling using a workflow adapted from the published study^17^ (isoTOP-ABPP^6^) to enrich and identify the probe-labelled protein fragment(s) as isotopically differentiated pairs of peptides (Fig. 3b and Supplementary Fig. 2). The predominant site that we identified for NQO2 labelling (Fig. 3c and Supplementary Fig. 6) is involved in cofactor binding^37^, and the parent mass determined for this peptide implies a net loss of N_2_H_4_ during coupling by the probe. Similarly, we resolved the predominant site in KDM1A (site 1) to two residues in the FAD-binding pocket that define the interface accommodating the dimethyl-isoalloxazine ring of the cofactor^38^ (Fig. 3d and Supplementary Fig. 7). A secondary site on the periphery of the active site was detected at much lower levels (Fig. 3d and Supplementary Fig. 8). For both NQO2 and KDM1A, the analytical data are consistent with MBI of the flavoenzymes by the hydrazine-based probes. Mass errors and statistics for all probe-labelled peptides of proteins discussed are summarized in Supplementary Tables 2 and 3, respectively.

**Fig. 3.**
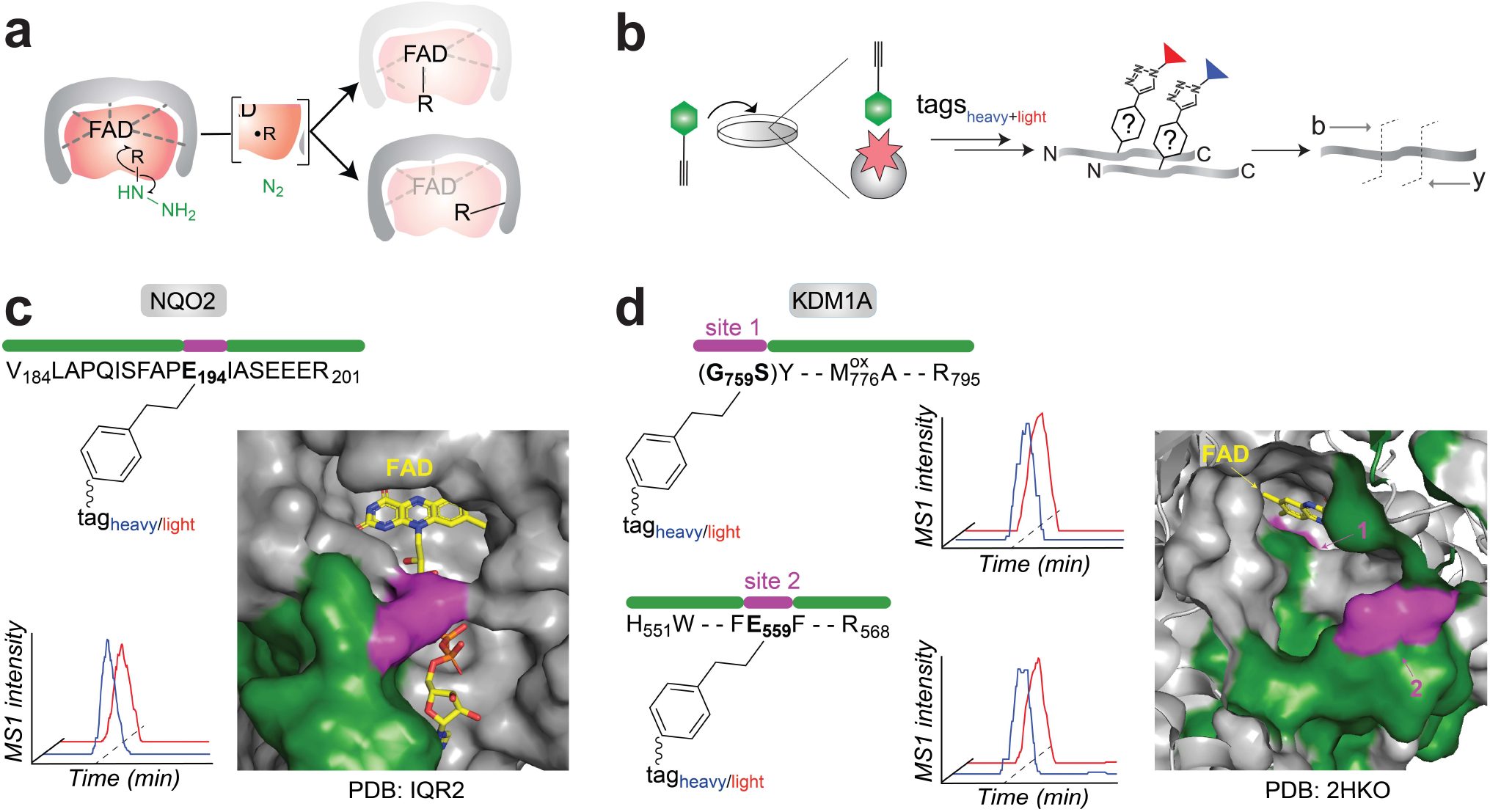
Covalent probes for noncovalent flavin enzymes. Identification of sites of probe labelling in NQO2 and KDM1A. **a**, Schematic for oxidative activation of hydrazine probes and subsequent coupling of resultant carbon-centered radical to FAD (red) and active site (grey). **b**, Condensed isoTOP-ABPP workflow to identify **3**-labelled peptides from treated cells overexpressing each flavoprotein target. **c**, Probe labelling site mapped onto the crystal structure of NQO2 bound to FAD (yellow). The identity of the probe **3**-labelled peptide (aa 184-201) and site are shown in green and purple, respectively (*left*). Extracted MS1 ion chromatograms of co-eluting heavy- and light-tagged peptides are shown in blue and red, respectively. The peptide modified in human NQO2 (PDB: IQR2) is highlighted. **d**, Probe labelling sites mapped onto the crystal structure of KDM1A bound to FAD (yellow). The identities of **3**-labelled peptides (aa 759-795 and 551-568) and sites are shown in green and purple, respectively (*left*). Extracted ion chromatograms of co-eluting heavy- and light-tagged peptides (blue and red, respectively) are shown in the *middle*. Peptides modified in human KDM1A (PDB: 2HKO) are highlighted.

To test the hypothesis that SCRN2 harbors a Cys-derived Glox that is captured by **3** to form a hydrazone (Fig. 4a), we applied the same method used for the flavoproteins above. Processed proteomes from **3**-treated SCRN2-transfected HEK293T cells were analyzed by LC-MS/MS using an unbiased search strategy as described in Supplemental Methods to reveal a **3**-Glox adduct at the position of Cys12 in the primary translation product (Fig. 4b and Supplementary Fig. 9). Consistent with the assignment of this peptide, its parent mass shifted the appropriate 2 Da when **3** was replaced with its 15*N*-isotopologue (**^15^N_2_-3**) (inverted spectra in Fig. 4b and Supplementary Fig. 10). Importantly, daughter b ions but not y ions of the MS2 fragmentation spectra shifted also by 2 Da, confirming that both *N*-atoms of hydrazine were retained in the protein-probe adduct. As positive controls, we also characterized **3**- and **^15^N_2_-3**-captured Glox (Supplementary Fig. 11 and 12) and Pyvl (Supplementary Fig. 13 and 14) peptides from SCRN3; they also demonstrated retention of the N_2_ unit. It is noteworthy that, unlike for the case of SCRN3, no pyruvoyl-appended SCRN2 (Pyvl-SCRN2) could be detected, suggesting that the *N*-terminus is more completely processed in isoform 2 and lending weight to the prior suggestion that Glox is the functional form of SCRN2/3. More broadly, the use of ^15^*N*-isotopologues to demonstrate whether the nitrogen atoms of the hydrazine are or are not retained during probe capture allows for immediate distinction of electrophilic versus radicaloid mechanisms of hydrazine inactivation, as applied below (Supplementary Fig. 15).

**Fig. 4.**
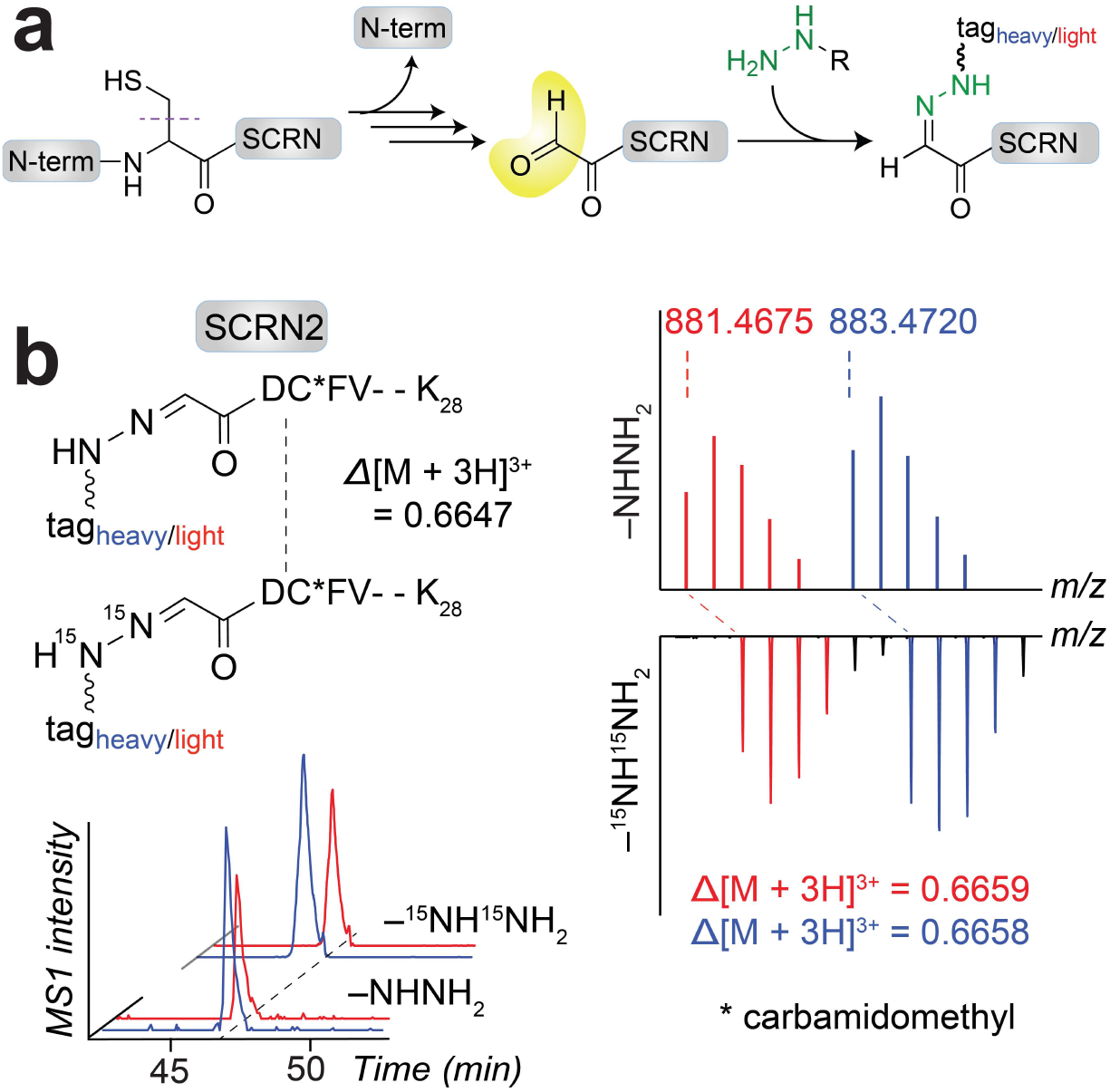
*De novo* discovery of Glox in SCRN2. **a**, Schematic for formation of Glox in SCRNs and subsequent capture by hydrazine probes. **b**, Monoisotopic *m/z* values of co-eluting heavy- and light-tagged peptides (blue and red, respectively) for the **3**-labelled Glox12-Lys28 SCRN2 peptide (*upper left*), extracted triply-charged MS1 ion chromatograms (*lower left*) and corresponding isotopic envelopes (*right*) shift the appropriate 2 Da (Δ*m/z* 0.6647) when **3** is substituted with ^**15**^**N**_2_-**3** (inverted y-axis).

We observed that only probe **2** (not **3**) targeted FTO (Fig. 1h and 2c). This probe selectivity contrasts with the behavior of the Glox- and FAD-harboring enzymes and suggests a distinct mechanism of probe capture of the Fe/2OG enzymes. It also establishes that relatively subtle details of structure can tune probes to be either more promiscuous or more selective with respect to different functional classes. Here, iron-hydrazine chelation may be an initial step, as hydrazide- and hydroxylamine-containing compounds have been shown to inhibit FTO^39^ by binding iron. However, to our knowledge, hydrazine reactivity leading to covalent enzyme modification has not yet been reported for a member of the Fe/2OG enzyme class. To define the mode of probe labelling, we used *N*-isotopologues of **2** to capture FTO from overexpressing HEK293T cells. SCRN3 was targeted as a control (Supplementary Fig. 16-19). Peptide pairs from both datasets were filtered according to retention time, parent mass, MS1 intensity and sample specificity (Supplementary Dataset 3). This analysis yielded only ^15^*N*-insensitive pairs (three) from the FTO dataset, suggesting a loss of at least the terminal nitrogen atom from the probe during coupling to the Fe/2OG enzyme (Supplementary Fig. 15 and Supplementary Fig. 20). The experimental MS2 spectra of the most abundant pair were searched against theoretical MS2 spectra for all possible peptide assignments for FTO. Using this approach, the predominant site of probe capture could be definitively assigned to His231 (Fig. 5a-b and Supplementary Fig. 21-22), one of the three cofactor ligands^28,29^. The mass of the peptide and corresponding ^15^*N* insensitivity are consistent with enzyme activation of the probe in a radical manifold that results in arylation of the His-coordinated side chain with loss of N_2_ (Fig. 5c). As can be seen from the crystal structure of FTO with bound nucleic acid substrate, His arylation should almost certainly disrupt binding of the substrate, 2OG cosubstrate, and/or iron cofactor (Fig. 5d). The data show that hydrazines define a new class of mechanism-based inactivator for this Fe/2OG enzyme. In addition to FTO, we identified 14 other Fe/2OG enzymes belonging to separate sequence clades and functional classes as high-reactivity probe targets (Fig. 1h and Supplementary Fig. 23), suggesting that the coupling mechanism is likely to be general.

**Fig. 5.**
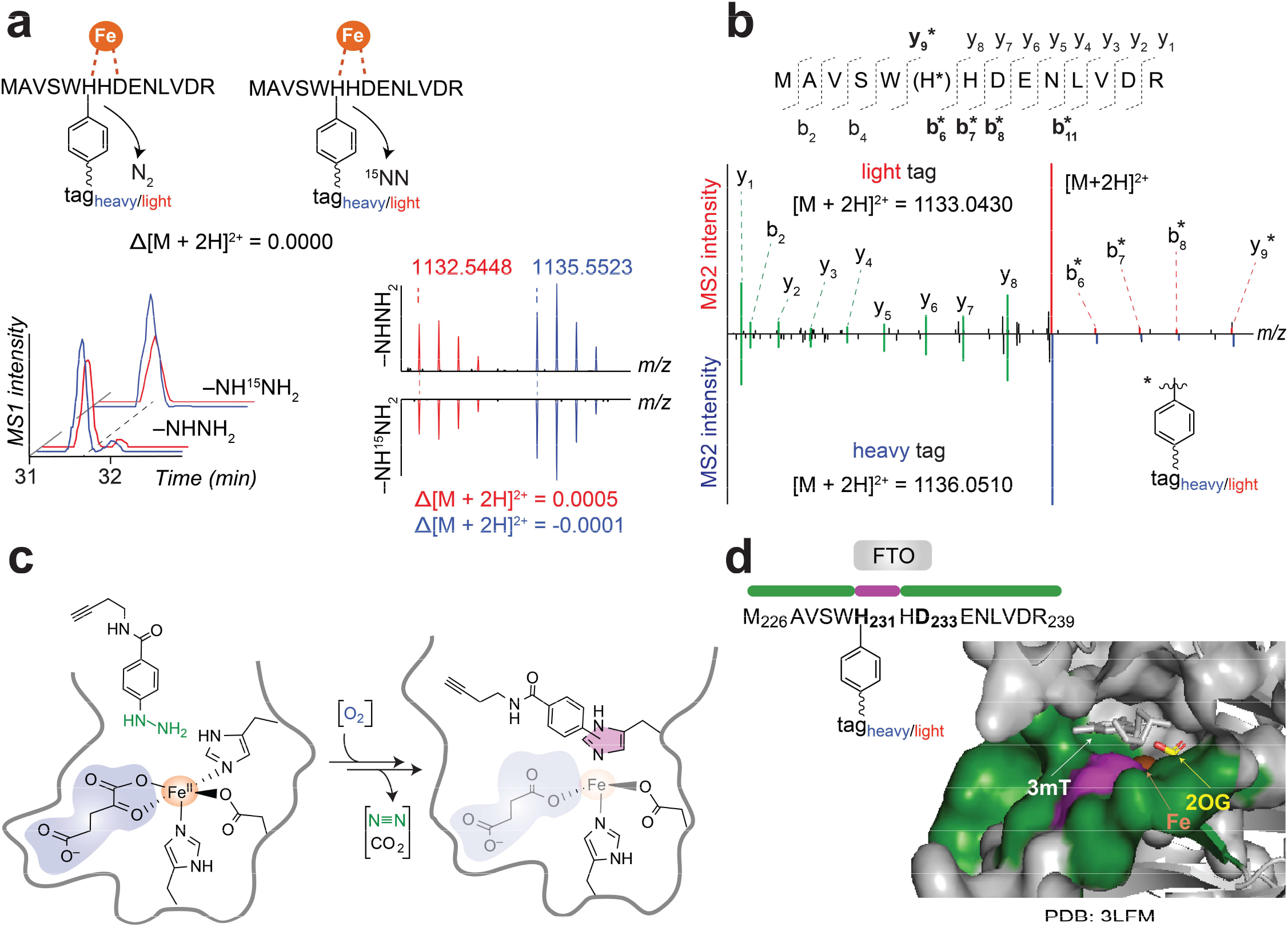
Discovery of a mechanism-based inactivator of FTO and other Fe/2OG enzymes. **a**, MS1 peptide (aa 226-239) pairs and corresponding isotopic envelopes from FTO-transfected cells treated with **2** versus ^**15**^**N-2**. Extracted ion chromatograms using monoisotopic *m/z* values of co-eluting heavy- and light-tagged peptides (blue and red, respectively) are shown on the *lower left*. **b,** Comparison of high-resolution MS2 spectra generated from light-versus heavy-tagged parent ions. **c,** Schematic for enzyme activation of probe **2** in a radical manifold, resulting in arylation of the active site His side chain that coordinates to the iron center. The hydrazine warhead, 2OG and iron are shown in green, blue and brown, respectively. **d,** Probe labelling site mapped onto FTO crystal structure. The identity of the probe **2**-labelled peptide (aa 226-239) and site (H231) are shown in green and purple, respectively (*left*). Modified peptide is highlighted in human FTO (PDB: 3LFM) with iron (brown), 2OG (yellow), and 3-methylthymine (gray) in the active site.

### Spatial resolution across diverse physiological contexts

We sought to assess whether hydrazine probes could also serve as versatile tools for activity-based imaging. For live cell imaging, **3**-treated HEK293T cells were visualized with and without in-cell fluorophore conjugation (CuAAC of Rh-N_3_), and with and without pretreatment with drug **4** under the same conditions described for the gel- and MS-based proteome profiling experiments (Fig. 6a). Labelling by **3** (*lower left*) was dependent on CuAAC (*upper right*), and **4** (*lower right*) diminished the mean fluorescence intensity by 48%, consistent with the average extent of competition seen in probe labelling profiles of soluble and membrane proteomes (45%) (Fig. 6a *upper left* and Supplementary Fig. 1). Fixed tissues from clinically relevant samples have altered metabolic activities for many enzyme classes. To evaluate the fraction of enzyme activities that can be captured in this context by our platform, the hydrazine-sensitive proteome preserved in formalin-fixed paraffin-embedded (FFPE) mouse lung tissue was investigated. Labelling by **3** (333 μM, 1 h) was strongly suppressed by a 10-fold excess of **4** (3.3 mM, 1 h), which diminished the composite fluorescence intensity by 84% (Fig. 6b). These results demonstrate that probe labelling can be effectively performed in both live cells and tissues. It should be possible to further elaborate probe molecules to develop more selective compounds that enable spatially resolved activity maps for existing histological samples and chemical druggability maps for existing genetics and epigenetics. With these complementary capabilities, the platform can be expected to enable discovery of new targets and biomarkers and how (and when) to drug them.

**Fig. 6.**
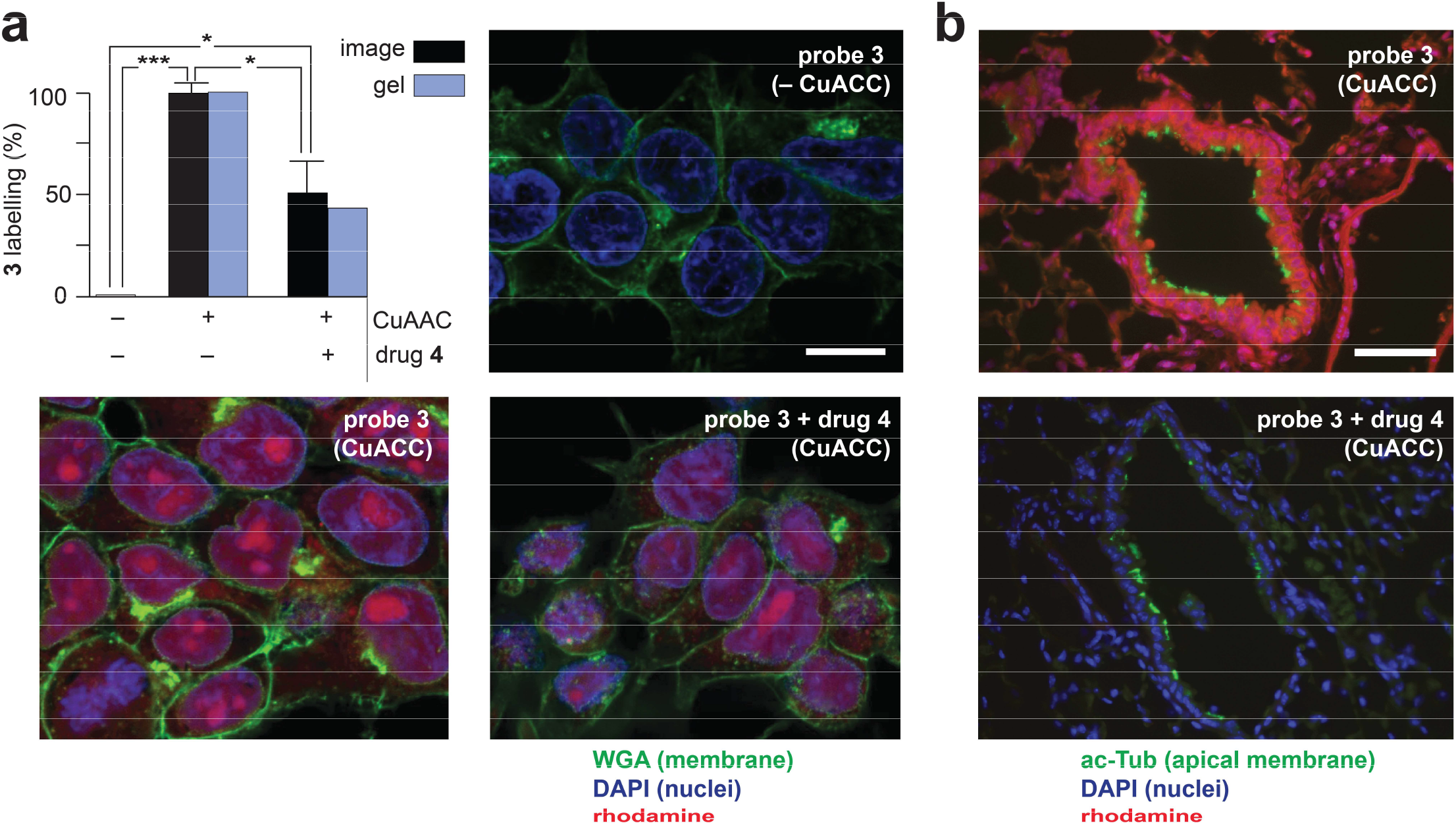
Imaging probe-sensitive activities in cells and tissues. **a**, Image-based analysis compared to gel-based analysis of (*upper left*) subcellular localization of probe reactivity in HEK293T cells. **3**-labelling in the absence (*upper right*) and presence (*bottom left*) of CuAAC and in competition with drug **4** (*bottom right*). Rhodamine is conjugated to the probe by CuAAC (magenta). Nuclei and membranes were stained with DAPI (blue) and WGA (green), respectively. Mean rhodamine labelling intensity was quantified using ImageJ (*n* = 3 per group), and group differences were evaluated by ANOVA followed by Bonferroni Post-hoc analysis (scale bar = 10 μm; * *p* < 0.05, *** *p* < 0.001). **b**, Image-based analysis of formalin-fixed paraffin-embedded (FFPE) mouse lung tissue labelled with probe **3** (magenta) and counterstained for acetylated tubulin (ac-Tub, green) and the nuclei with DAPI (blue) (*upper*). A ten-fold excess of competing drug **4** reduced probe labelling by 84% (scale bar = 50 μm; *p*-value = 3.375 × 10^−8^, Student’s *t*-test) (*lower*).

## DISCUSSION

Classical ABPP represented a creative fusion of established and emerging technologies. Amino acid-selective chemical modifiers, developed decades before, were repurposed to identify proteins with a specific type of residue serving a functional role. Whereas labelling every instance of a specific amino acid (e.g. all serines, cysteines etc.) in a raw, complex proteome would not obviously, *a priori,* be a profitable endeavor, the emerging power of MS^n^ proteomics to resolve nmol levels of thousands, if not tens of thousands, of proteins in a complex mixture enabled individual targets to be identified^7^. Moreover, it rested on the tenet that proteins having a specific amino acid to engage covalently as part of its function often tune that residue to be hyperreactive with respect to purely structural amino acids of the same type. Thus, limited treatments with sub-saturating probe were used to selectively identify the functional residues of the type being targeted. Once a particular modifier of an active site residue is identified, other molecules that bind selectively to that site can be identified by virtue of their ability to prevent probe attachment^1^. Importantly, because the chemical functionalities being targeted invariably arose from the common amino acids, selectivity for a single type was crucial to avoid excessive complexity and enable sorting of the many targets in a complex mixture.

Recognizing the historical bias for genetically encoded nucleophilic groups, we previously reversed the polarity of ABPP by using a nucleophilic probe, with the intent of discovering electrophiles^17^. Such species are present in proteins primarily in the forms of regulatory post-translational modifications and (in)organic enzyme cofactors. In this other less explored hemisphere of the proteome, reactive functionality is comparatively rare and almost uniformly associated with chemical function. In this case, use of a single probe warhead to capture many or all electrophilic or electron-deficient chemical functionality in the proteome without compromising the ability to resolve the individual targets by MS^n^ approaches seemed feasible. Our data demonstrate that, by virtue of their nucleophilicity and capacity to form carbon radicals that can attach promiscuously to side chains and backbones of proteins, organohydrazines provide such a versatile, if not completely general, warhead to identify electron deficient functional groups in proteins without prior knowledge of their identity or even their presence. The hydrazine probes deployed herein are mechanism-based, active-site-directed and blocked by inhibitors that occupy active sites. As such, they can be used in the same way that amino acid-selective modifiers have been applied so successfully. Further, their capacity to access even the most isolated tissue space of the brain (e.g., in the antidepressant, phenelzine)^18,40^ will make them useful for *in vivo* imaging and assessment of target engagement or tissue distribution.

A central tenant of the ABPP approach is the ability to measure biomolecular activity, independent of transcription and translation levels, to uncover post-transcriptional and metabolic modes of regulation^7^. Methods to globally and directly measure active states of enzymes are invaluable for understanding how their activities are (dys)regulated, how they can be pharmacologically modulated, and in which physiological context doing so might be therapeutically relevant. For Fe/2OG enzymes, for example, activity is impacted not only by gene and transcript levels but also by the availability of oxygen, iron and energetic metabolites, as these enzymes are cellular sensors that control the fundamental response pathways in which their cofactors/cosubstrates participate (e.g. hypoxia, metal trafficking and 2OG metabolism)^41^. Similarly, flavoenzyme activation is determined not only by the availability of vitamin precursors and integrity of biosynthetic enzymes required for cofactor generation but also by its trafficking and compartmentalization, which must also be precisely titrated to mirror dynamic levels of myriad client apoenzymes with diverse and essential functions^42^. For these reasons, hydrazine-based probes represent a novel approach for evaluating cofactor-dependent enzyme biochemistry and facilitating new areas of therapeutic development.

## Methods

Methods section is provided in the Supplementary Information with details available.

## Acknowledgements

The authors are grateful for the computational support from Simone Sidoli from Department of Biochemistry at Albert Einstein College of Medicine, Hee Jong Kim and Benjamin A. Garcia from Biochemistry & Molecular Biophysics at University of Pennsylvania and Benjamin F. Cravatt from Department of Chemistry at The Scripps Research Institute. We thank Xian Han and Junmin Peng from Department of Structural Biology at St. Jude Children’s Research Hospital for developing postCIMAGE script (R), Sara M. Martin (Matthews group) for mutagenesis of MAOA plasmid, Ellen Heber-Katz from Lankenau Institute for Medical Research for providing 1,4-DPCA, Bin Yu from School of Pharmaceutical Sciences at Zhengzhou University for providing KDM1A inhibitors, Philip A. Cole from Department of Biological Chemistry and Molecular Pharmacology at Harvard Medical School for providing KDM1A, and Philip E. Dawson from The Scripps Research Institute for assistance with preparation of the TEV tags. This work was supported by the University of Pennsylvania (M.L.M), Oberlin College (W.H.P.), National Institutes of Health [NIAMS R01AR075241 (A.M.G.), NIDA R00DA35865 (M.W.B.), NINDS R15NS108183 (P.D.M.) and NCI R01-CA-181633-01A1 (E.S.W)] and American Cancer Society [RSG-15-027-01 (E.S.W)].

## Author contributions

Z.L. and M.L.M. conceived the research, wrote the manuscript and analyzed data, Z.L. developed methods and performed experiments, X.W., Z.L. and W.H.P. designed and synthesized probes, K.A.B. expressed and purified NQO2 and performed associated experiments, L.H. developed computational workflow for MS data analysis, R.M.S. managed CIMAGE software for proteomics data, M.A. tested inhibitors, K.H. generated dendrograms for enzyme families, E.J.O. synthesized TEV tags, N.S. and E.S.W. developed imaging methods for FFPE tissues, P.D.M., A.M.G. and M.W.B. developed imaging methods for cells, all authors revised the manuscript.

## Data availability

The data supporting the findings in this study are available within the paper, any associated data are available from the corresponding author upon request.

