## Supplementary Information for "Hydrazines as versatile chemical biology probes and drug-discovery tools for cofactor-dependent enzymes"

### 1. Supplementary Tables

#### Supplementary Table 1. Average SILAC ratios from two human cell lines for generation of heatmap for probe 1-, 2- and 3-targeted proteins.

| *functional group*[‡] | *UniProt ID* | *description* | *probe 1* | | *probe 2* | | *probe 3* | |
|---|---|---|---|---|---|---|---|---|
| | | | *enrichment** | *competition** | *enrichment** | *competition** | *enrichment**[†] | *competition**[†] |
| succinimide | Q99538 | LGMN Legumain | 20 | 12 | 20 | 4 | | |
| gloxylyl | Q96FV2 | SCRN2 Secernin2 | 20 | 15 | 20 | 12 | 20 | 20 |
| gloxylyl | Q0VDG4 | SCRN3 Secernin3 | 20 | 6 | 18 | 15 | 2 | 3 |
| pyruvovl | P17707 | AMD1 Sadenosylmethionine decarboxylase proenzyme | 17 | 5 | 18 | 5 | | |
| sulfatase | P15586 | GNS cetylglucosamine6sulfatase | 10 | 3 | 7 | 2 | 1 | |
| flavin | Q9UHG3 | PCYOX1 Prenylcysteine oxidase 1 | | | | | | 8 |
| flavin | P27338 | MAOB Amine oxidase [flavincontaining] B | | | | | 20 | 20 |
| flavin | P21397 | MAOA Amine oxidase [flavincontaining] A | | | | | 20 | 20 |
| flavin | P16083 | NQO2 Ribosyldihydronicotimide dehydrogese | | | 4 | 1 | 20 | 5 |
| flavin | O60341 | KDM1A Lysinespecific histone demethylase 1A | | | 20 | 2 | 19 | 11 |
| flavin | Q8N2H3 | PYROXD2 Pyridine nucleotidedisulfide oxidoreductase domai | | | 20 | 20 | | |
| flavin | P30043 | BLVRB Flavin reductase (DPH) | 18 | 2 | 3 | 2 | | |
| flavin | O75629 | CREG1 Protein CREG1 | 20 | | | | | |
| flavin | O60427 | FADS1 Fatty acid desaturase 1 | | | 20 | 4 | 14 | 5 |
| flavin | P41229 | KDM5C Lysinespecific demethylase 5C | | | 19 | 3 | | |
| flavin | Q96DG6 | CMBL Carboxymethylenebutenolidase homolog | | 2 | 19 | 3 | 15 | 1 |
| flavin | Q8NB78 | KDM1B Lysinespecific histone demethylase 1B | | | 17 | 2 | | 6 |
| flavin | O15254 | ACOX3 Peroxisomal acylcoenzyme A oxidase 3 | | | 20 | 1 | | |
| flavin | Q15067 | ACOX1 Peroxisomal acylcoenzyme A oxidase 1 | | | 20 | 3 | | |
| flavin | P49748 | ACADVL Very longchain specific acylCoA dehydrogese, m | 2 | 1 | 5 | 3 | 1 | |
| flavin | Q9H9P8 | L2HGDH L2hydroxyglutarate dehydrogese, mitochondrial | | | | | | 8 |
| flavin | P31040 | SDHA Succite dehydrogese | 2 | 1 | 1 | 1 | 1 | 10 |
| Fe/2OG | Q9C0B1 | FTO Alphaketoglutaratedependent dioxygese FTO | | 1 | 14 | 5 | | |
| Fe/2OG | P13674 | P4HA1 Prolyl 4hydroxylase subunit alpha1 | 15 | 2 | 8 | 3 | 7 | 2 |
| Fe/2OG | O60568 | PLOD3 Procollagenlysine,2oxoglutarate 5dioxygese 3 | 14 | 2 | 18 | 5 | | |
| Fe/2OG | Q32P28 | LEPRE1 Prolyl 3hydroxylase 1 | 8 | 3 | 8 | 2 | | |
| Fe/2OG | Q02809 | PLOD1 Procollagenlysine,2oxoglutarate 5dioxygese 1 | 6 | 2 | 8 | 3 | | 2 |

| | | | | | | | | |
|---|---|---|---|---|---|---|---|---|
| Fe/2OG | Q13686 | ALKBH1 Alkylated D repair protein alkB homolog 1 | | | 20 | 8 | 1 | 1 |
| Fe/2OG | Q6NS38 | ALKBH2 Alphaketoglutaratedependent dioxygese alkB hom | | | | 20 | | 4 |
| Fe/2OG | Q9BV57 | ADI1 1,2dihydroxy3keto5methylthiopentene dioxyge | | 1 | | | 20 | |
| Fe/2OG | Q96SZ5 | ADO 2aminoethanethiol dioxygese | | | | 2 | 20 | 1 |
| Fe/2OG | O15460 | P4HA2 Prolyl 4hydroxylase subunit alpha2 | | | 20 | 3 | 1 | |
| Fe/2OG | O00469 | PLOD2 Procollagenlysine,2oxoglutarate 5dioxygese 2 | | 2 | 6 | 3 | | |
| Fe/2OG | Q9NVH6 | TMLHE Trimethyllysine dioxygese, mitochondrial | | | | | 15 | 1 |
| Fe/2OG | Q9UPP1 | PHF8 Histone lysine demethylase PHF8 | | | 20 | 15 | 2 | 2 |
| Fe/2OG | Q5SRE7 | PHYHD1 PhytanoylCoA dioxygese domaincontaining protei | | | 14 | 4 | 20 | |
| Cu | Q9H7Z7 | PTGES2 Prostaglandin E synthase 2 | | | 2 | 2 | 20 | 14 |
| Cu | Q6UVY6 | MOXD1 DBHlike monooxygese protein 1 | | | | 3 | 20 | 5 |
| Cu | Q14061 | COX17 Cytochrome c oxidase copper chaperone | | | | | 12 | 3 |
| iron iron | P31350 | RRM2 Ribonucleosidediphosphate reductase subunit M2 | 2 | 1 | 9 | 1 | | |
| iron iron | Q7LG56 | RRM2B Ribonucleosidediphosphate reductase subunit M2 B | | | 7 | 2 | 20 | |
| iron iron | O00767 | SCD AcylCoA desaturase | | | 20 | 8 | 20 | 20 |
| iron iron | Q99807 | COQ7 5demethoxyubiquinone hydroxylase, mitochondrial | | | | | 20 | 2 |
| heme | Q02318 | CYP27A1 Sterol 26hydroxylase, mitochondrial | | | 20 | 3 | | |
| heme | O95864 | FADS2 Fatty acid desaturase 2 | | | 20 | 4 | 20 | 20 |
| heme | Q16850 | CYP51A1 Lanosterol 14alpha demethylase | | | 14 | 6 | | 3 |
| heme | O15121 | DEGS1 Sphingolipid delta(4)desaturase DES1 | | | | 6 | | |
| heme | Q92626 | PXDN Peroxidasin homolog | | | 20 | 5 | | |
| heme | P30519 | HMOX2 Heme oxygese 2 | | | 18 | 4 | 16 | 3 |
| heme | P99999 | CYCS Cytochrome c | | 1 | 7 | 0 | 3 | 1 |
| heme | P53701 | HCCS Cytochrome ctype heme lyase | | | | 2 | 14 | 7 |
| iron | P02792 | FTL Ferritin light chain | | 2 | 7 | 3 | | 3 |
| iron | P02794 | FTH1 Ferritin heavy chain | 2 | 2 | 20 | 5 | 5 | |

‡protein functional groups based on UniProt or literature reports.

*average ratios from HEK293T and MDA-MB-231 cell lines.

†average ratios from soluble and membrane proteomes in each cell line.

**Supplementary Table 2. Mass errors for chemically modified peptides in various proteins.**

| protein | probe | captured peptide‡ | modification | parent ion intensity | charge | $m/z_{theoretical}$ | $m/z_{measured}$ | error (ppm) |
|---|---|---|---|---|---|---|---|---|
| SCRN3 | **2** | Glox*DTFVALPPATVDNR | $TEV_{heavy}$ | 2.9E+06 | 2 | 1074.0549 | 1074.0543 | -0.6 |
| | | Glox*DTFVALPPATVDNR | $TEV_{light}$ | 2.3E+06 | 2 | 1071.0480 | 1071.0483 | 0.3 |
| | | Pyvl*DTFVALPPATVDNR | $TEV_{heavy}$ | 1.4E+06 | 2 | 1081.0627 | 1081.0609 | -1.7 |
| | | Pyvl*DTFVALPPATVDNR | $TEV_{light}$ | 9.9E+05 | 2 | 1078.0558 | 1078.0553 | -0.5 |
| | **$^{15}$N-2** | Glox*DTFVALPPATVDNR | $TEV_{heavy}$ | 4.1E+07 | 2 | 1074.5534 | 1074.5531 | -0.3 |
| | | Glox*DTFVALPPATVDNR | $TEV_{light}$ | 3.3E+07 | 2 | 1071.5465 | 1071.5464 | -0.1 |
| | | Pyvl*DTFVALPPATVDNR | $TEV_{heavy}$ | 2.5E+07 | 2 | 1081.5612 | 1081.5610 | -0.2 |
| | | Pyvl*DTFVALPPATVDNR | $TEV_{light}$ | 1.9E+07 | 2 | 1078.5543 | 1078.5548 | 0.5 |
| | **3** | Glox*DTFVALPPATVDNR | $TEV_{heavy}$ | 7.5E+06 | 2 | 1088.0742 | 1088.0721 | -1.9 |
| | | Glox*DTFVALPPATVDNR | $TEV_{light}$ | 5.5E+06 | 2 | 1085.0673 | 1085.0638 | -3.2 |
| | | Pyvl*DTFVALPPATVDNR | $TEV_{heavy}$ | 1.2E+06 | 2 | 1095.0784 | 1095.0789 | 0.5 |
| | | Pyvl*DTFVALPPATVDNR | $TEV_{light}$ | 7.2E+05 | 2 | 1092.0714 | 1092.0717 | 0.3 |
| | **$^{15}N_2$-3** | Glox*DTFVALPPATVDNR | $TEV_{heavy}$ | 3.0E+08 | 2 | 1089.0676 | 1089.0676 | 0 |
| | | Glox*DTFVALPPATVDNR | $TEV_{light}$ | 3.0E+08 | 2 | 1086.0607 | 1086.0610 | 0.3 |
| | | Pyvl*DTFVALPPATVDNR | $TEV_{heavy}$ | 1.5E+07 | 2 | 1096.0754 | 1096.0758 | 0.4 |
| | | Pyvl*DTFVALPPATVDNR | $TEV_{light}$ | 1.5E+07 | 2 | 1093.0685 | 1093.0681 | -0.4 |
| SCRN2 | **3** | Glox*DCFVSVPPASAIPAVIFAK | $TEV_{heavy}$ | 6.8E+06 | 3 | 883.4732 | 883.4720 | -1.4 |
| | | Glox*DCFVSVPPASAIPAVIFAK | $TEV_{light}$ | 5.2E+06 | 3 | 881.4686 | 881.4675 | -1.2 |
| | **$^{15}N_2$-3** | Glox*DCFVSVPPASAIPAVIFAK | $TEV_{heavy}$ | 1.5E+08 | 3 | 884.1379 | 884.1378 | -0.1 |
| | | Glox*DCFVSVPPASAIPAVIFAK | $TEV_{light}$ | 1.5E+08 | 3 | 882.1333 | 882.1334 | 0.1 |
| KDM1A | **3** | (GS)*YSYVAAGSSGNDYDLM$^{ox}$AQPITPGPSIPGAPQPIPR | $TEV_{heavy}$ | 9.1E+05 | 4 | 1085.0387 | 1085.0359 | -2.6 |
| | | (GS)*YSYVAAGSSGNDYDLM$^{ox}$AQPITPGPSIPGAPQPIPR | $TEV_{light}$ | 9.8E+05 | 4 | 1083.5352 | 1083.5416 | 5.9 |
| | | HWDQDDDFE*FTGSHLTVR | $TEV_{heavy}$ | 3.6E+05 | 4 | 699.3305 | 699.3301 | -0.6 |
| | | HWDQDDDFE*FTGSHLTVR | $TEV_{light}$ | 6.3E+05 | 4 | 697.8270 | 697.8264 | -0.9 |
| NQO2 | **3** | VLAPQISFAPE*IASEEER | $TEV_{heavy}$ | 4.5E+06 | 3 | 859.1231 | 859.1224 | -0.8 |
| | | VLAPQISFAPE*IASEEER | $TEV_{light}$ | 3.1E+06 | 3 | 857.1185 | 857.1180 | -0.6 |
| FTO | **2** | MAVSWH*HDENLVDR | $TEV_{heavy}$ | 1.8E+06 | 2 | 1135.5494 | 1135.5523 | 2.6 |
| | | MAVSWH*HDENLVDR | $TEV_{light}$ | 1.3E+06 | 2 | 1132.5426 | 1132.5448 | 1.9 |
| | **$^{15}$N-2** | MAVSWH*HDENLVDR | $TEV_{heavy}$ | 1.7E+06 | 2 | 1135.5494 | 1135.5522 | 2.5 |
| | | MAVSWH*HDENLVDR | $TEV_{light}$ | 9.7E+05 | 2 | 1132.5426 | 1132.5453 | 2.4 |
| | **2** | EE(PY)*FGMGK | $TEV_{heavy}$ | 5.1E+06 | 3 | 540.2622 | 540.2627 | 0.9 |

| | | | | | | | |
|---|---|---|---|---|---|---|---|
| | EE(PY)*FGMGK | $TEV_{light}$ | 4.0E+06 | 3 | 538.2574 | 538.2582 | 1.5 |
| $^{15}$N-2 | EE(PY)*FGMGK | $TEV_{heavy}$ | 4.5E+06 | 3 | 540.2622 | 540.2626 | 0.7 |
| | EE(PY)*FGMGK | $TEV_{light}$ | 3.5E+06 | 3 | 538.2574 | 538.2584 | 1.9 |

‡ each peptide was detected in at least two technical or biological replicates ($n \geq 2$).

* site of modification by probe clicked with TEV tags.

$^{OX}$ methionine oxidation.

### Supplementary Table 3. Statistics of chemically modified peptides in KDM1A, NQO2 and FTO.

| protein | probe | captured peptide | modification | $m/z_{theoretical}$ | average $m/z_{measured}$ | error (ppm) | SD | $n$ |
|---|---|---|---|---|---|---|---|---|
| KDM1A | **3** | (GS)*YSYVAAGSSGNDYDLM$^{ox}$AQPITPGPSIPGAPQPIPR | $TEV_{heavy}$ | 1085.0387 | 1085.0359 | -2.6 | 0.0007 | 58§ |
| | | (GS)*YSYVAAGSSGNDYDLM$^{ox}$AQPITPGPSIPGAPQPIPR | $TEV_{light}$ | 1083.5352 | 1083.5416 | 5.9 | 0.0008 | 53§ |
| | | HWDQDDDFE*FTGSHLTVR | $TEV_{heavy}$ | 699.3305 | 699.3301 | -0.6 | 0.0034 | 54§ |
| | | HWDQDDDFE*FTGSHLTVR | $TEV_{light}$ | 697.8270 | 697.8264 | -0.9 | 0.0005 | 44§ |
| NQO2 | **3** | VLAPQISFAPE*IASEEER | $TEV_{heavy}$ | 859.1231 | 859.1224 | -0.8 | 0.0006 | 40§ |
| | | VLAPQISFAPE*IASEEER | $TEV_{light}$ | 857.1185 | 857.1180 | -0.6 | 0.0015 | 36§ |
| FTO | **2** | MAVSWH*HDENLVDR | $TEV_{heavy}$ | 1135.5494 | 1135.5523 | 2.6 | 0.0015 | 88† |
| | | MAVSWH*HDENLVDR | $TEV_{light}$ | 1132.5426 | 1132.5448 | 1.9 | 0.0017 | 87† |
| | **$^{15}$N-2** | MAVSWH*HDENLVDR | $TEV_{heavy}$ | 1135.5494 | 1135.5522 | 2.5 | 0.0019 | 21§ |
| | | MAVSWH*HDENLVDR | $TEV_{light}$ | 1132.5426 | 1132.5453 | 2.4 | 0.0018 | 20§ |
| | **2** | EE(PY)*FGMGK | $TEV_{heavy}$ | 540.2622 | 540.2627 | 0.9 | 0.0008 | 129† |
| | | EE(PY)*FGMGK | $TEV_{light}$ | 538.2574 | 538.2582 | 1.5 | 0.0012 | 138† |
| | **$^{15}$N-2** | EE(PY)*FGMGK | $TEV_{heavy}$ | 540.2622 | 540.2626 | 0.7 | 0.0009 | 50§ |
| | | EE(PY)*FGMGK | $TEV_{light}$ | 538.2574 | 538.2584 | 1.9 | 0.0011 | 54§ |

* site of modification by probe clicked with TEV tags.

§ the sum of detections ($n \geq 20$) in MS1 spectra from two technical replicates ($n = 2$).

† the sum of detections ($n \geq 87$) in MS1 spectra from two biological replicates (four technical replicates, $n = 4$).

### 2. Supplementary Figures

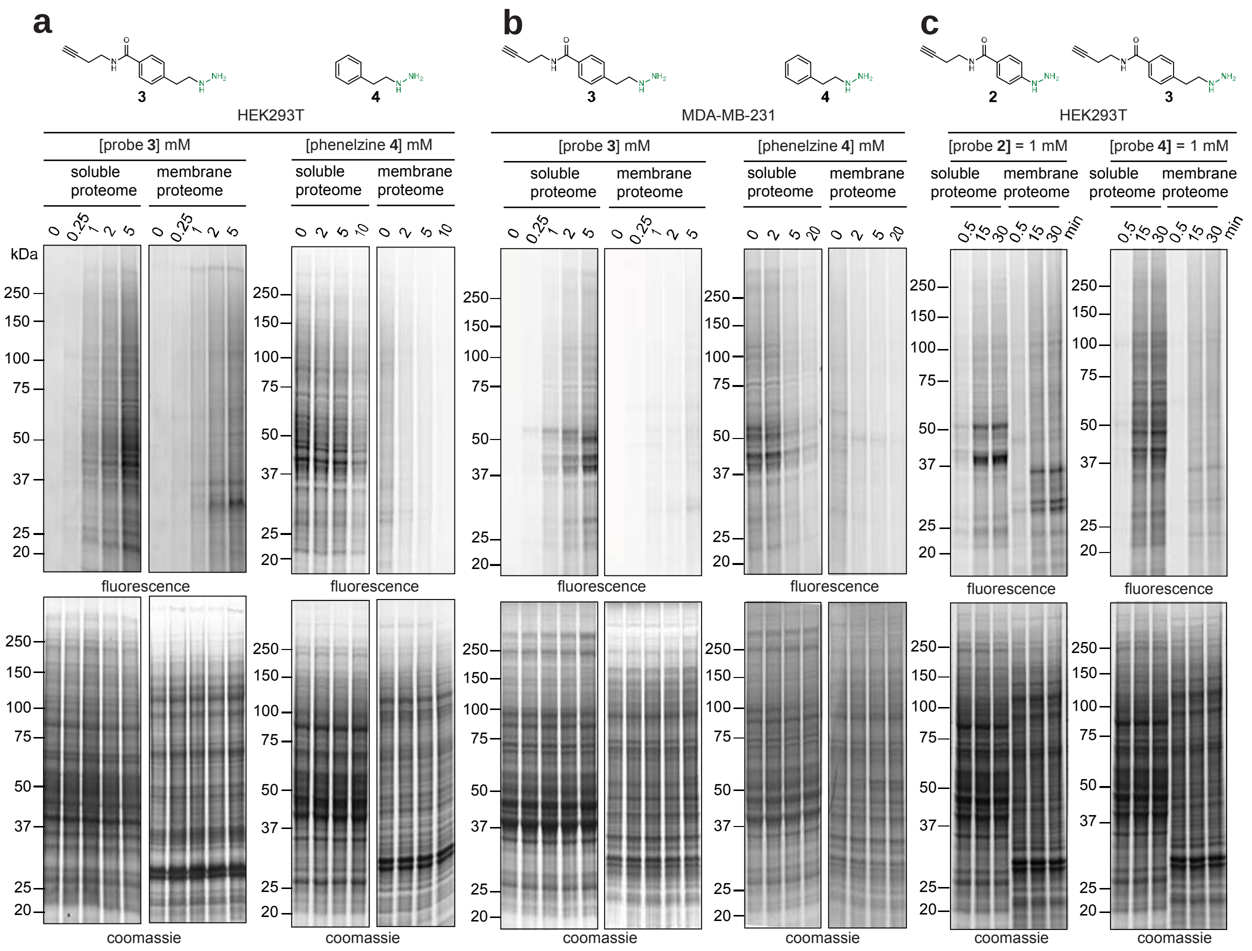

**Supplementary Fig. 1**. **SDS-PAGE profiles for hydrazine probe-treated human cells.**

**a**, dose-dependent labelling profiles by probe **3** in the soluble and membrane proteomes of HEK293T cells (*upper left*), competition of probe **3** labelling by non-clickable analog **4** in the soluble and membrane proteomes of HEK293T cells pretreated with varying concentrations of **4** (*upper right*). Corresponding expression profiles are shown after Coomassie staining (*lower*). **b**, dose-dependent labelling profiles by probe **3** in the soluble and membrane proteomes of MDA-MB-231 cells (*upper left*), competition of probe **3** labelling by non-clickable analog **4** in the soluble and membrane proteomes of MDA-MB-231 cells pretreated with varying concentrations of **4** (*upper right*). Corresponding expression profiles are shown after Coomassie staining (*lower*). **c**, time-dependent labelling profiles by probes **2** and **3** in the soluble and membrane proteomes of HEK293T cells incubated for varying times. Concentrations and molecular weight markers are indicated.

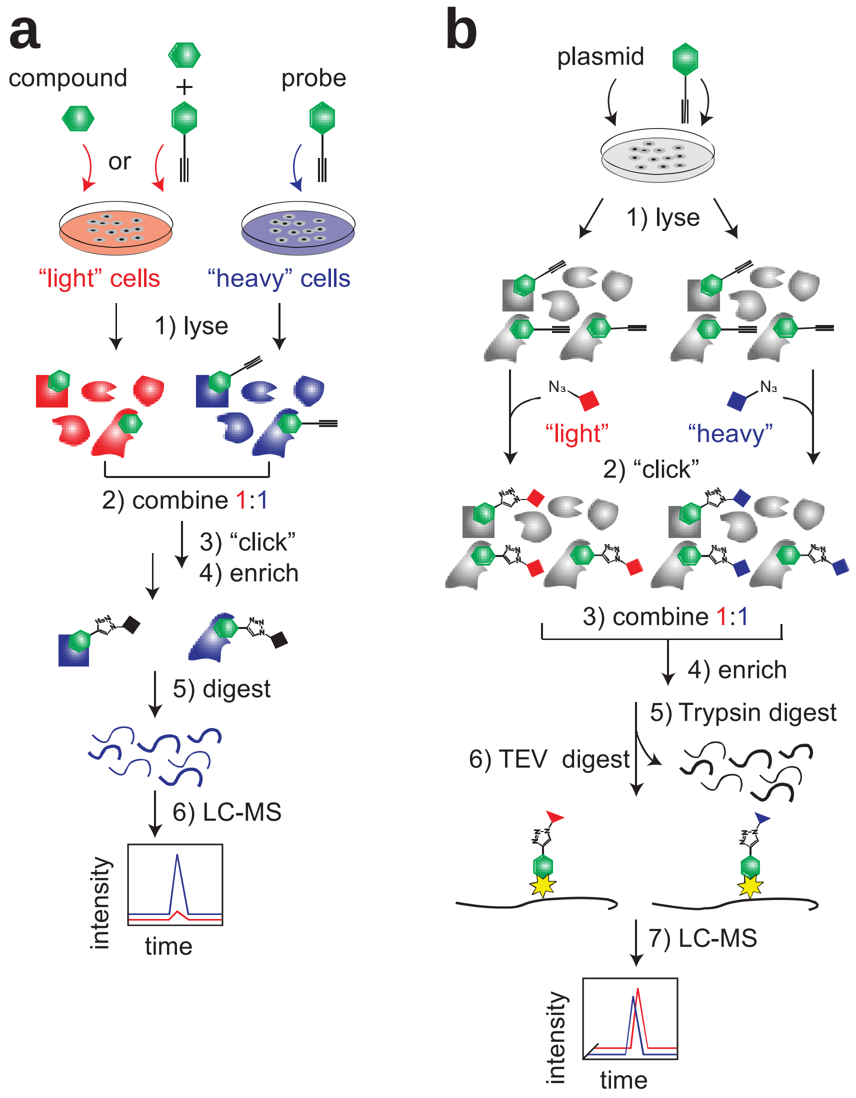

**Supplementary Fig. 2**. **Workflow schematics for proteomics experiments.**

**a**, Workflow for enrichment and competition experiments in Fig. 1. SILAC cells are treated with probe and/or compound, lysed, and combined (heavy: light = 1:1) for conjugation to biotin-azide tags by CuAAC. Probe-labelled proteins are then enriched by streptavidin resin, proteolytically digested on-bead by trypsin, and desalted for LC-MS/MS analysis. **b**, Workflow for isoTOP-ABPP experiments. Lysates of probe-treated cells are conjugated to isotopically differentiated biotin-azide tags (heavy and light, respectively) containing an internal TEV protease cleavage site and combined in a 1:1 ratio. Probe-labelled proteins are enriched by streptavidin resin, digested on-bead first by trypsin to remove unlabelled peptides, then by TEV protease to release isotopic probe-labelled pairs for LC-MS/MS analysis.

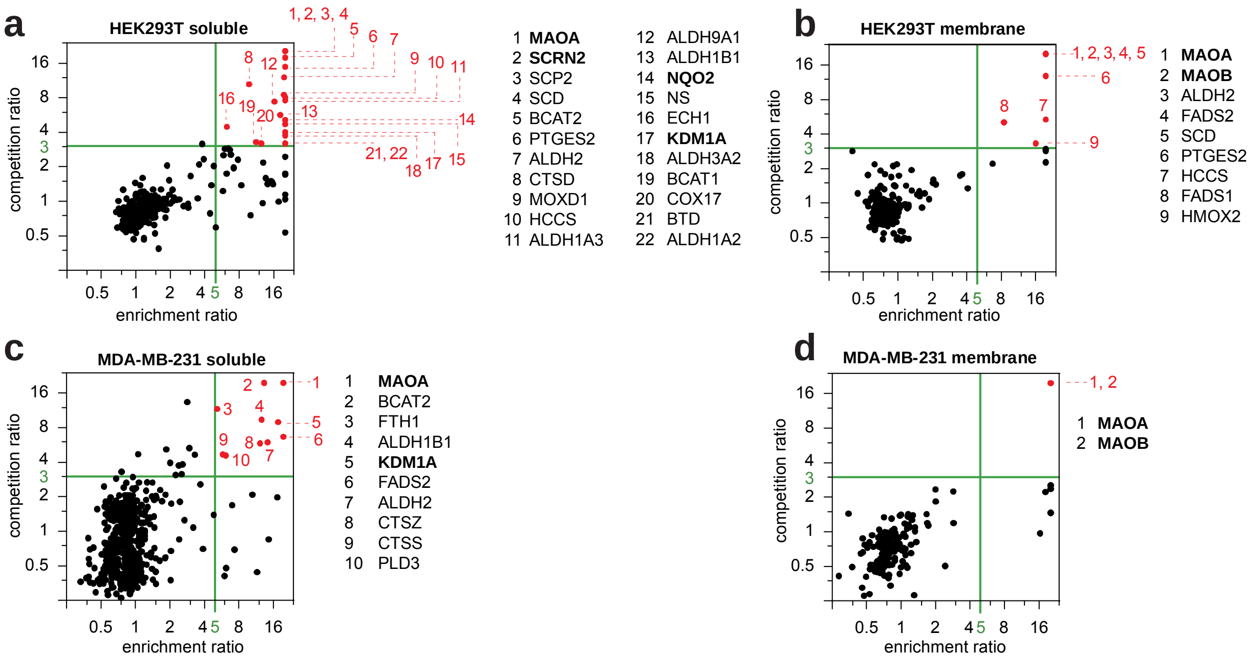

**Supplementary Fig. 3. Identification of protein targets of probe 3 in two human cell lines.**

**a**, Quadrant plot of average competition versus enrichment SILAC ratios from quantitative proteomics experiments for the soluble proteomes of HEK293T cells. Highly promising targets are found in the *upper right* quadrant (22 in total), numbered and highlighted in red in the plot and listed by protein symbol to the *right* of the plot. **b**, quadrant plot of average competition versus enrichment SILAC ratios from quantitative proteomics experiments in the membrane proteomes of HEK293T cells. Highly promising targets are found in the *upper right* quadrant (9 in total), numbered and highlighted in red in the plot and listed by protein symbol to the *right* of the plot. **c**, quadrant plot of average competition versus enrichment SILAC ratios from quantitative proteomics experiments in the soluble proteomes of MDA-MB-231 cells. Highly promising targets are found in the *upper right* quadrant (10 in total), numbered and highlighted in red in the plot and listed by protein symbol to the *right* of the plot. **d**, quadrant plot of average competition versus enrichment SILAC ratios from quantitative proteomics experiments in the membrane proteomes of MDA-MB-231 cells. Highly promising targets are found in the *upper right* quadrant (9 in total), numbered and highlighted in red in the plot and listed by protein symbol to the *right* of the plot. Bolded targets are proteins further investigated and discussed in this work.

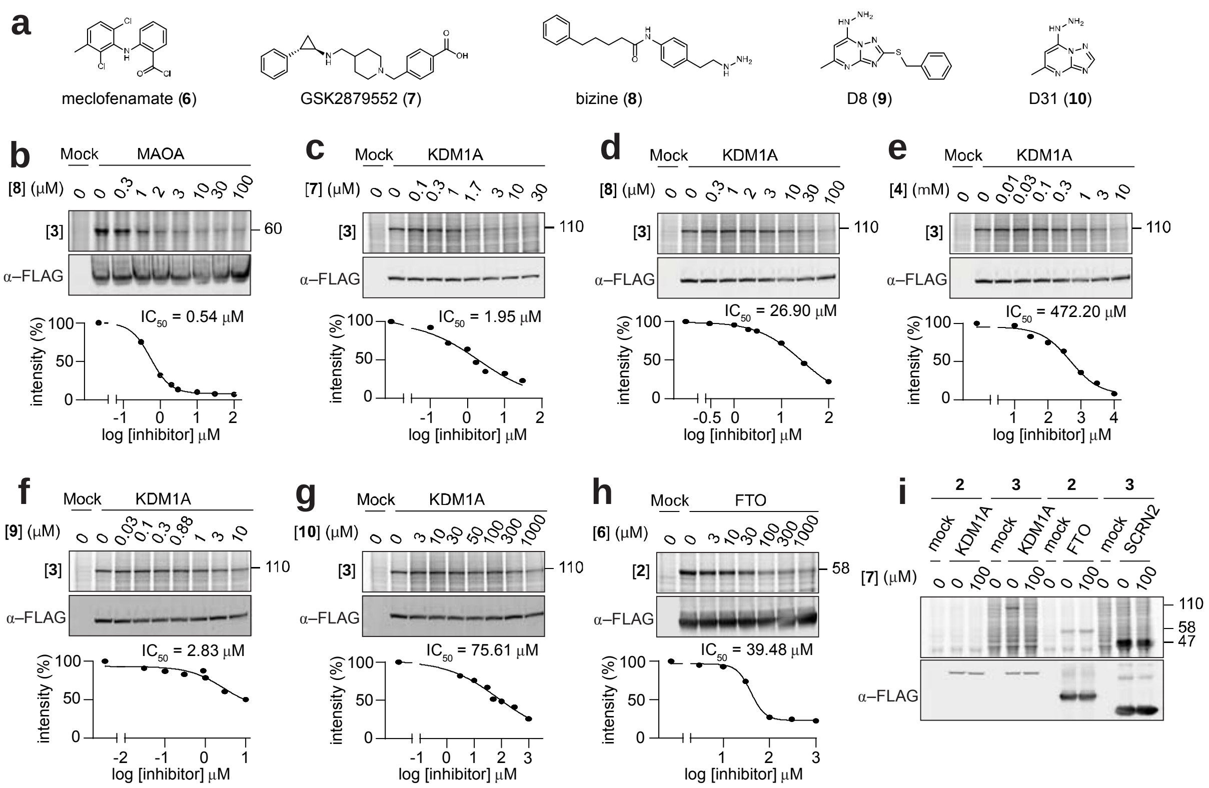

**Supplementary Fig. 4. Inhibition of probe-labelling by various inhibitors.**

**a**, structures of reported KDM1A inhibitors (**7**, **8**, **9** and **10**), and FTO inhibitor **6**. **b**, gel (*upper*) and gel-derived (*lower*) curve plot showing dose-dependent competitor **8** blockade of probe **3** labelling of recombinant MAOA in HEK29T cells. **c**, gel (*upper*) and gel-derived curve plot (*lower*) showing dose-dependent competitor **7** blockade of probe **3** labelling of recombinant KDM1A in HEK29T cells. **d**, gel (*upper*) and gel-derived curve plot (*lower*) showing dose-dependent competitor **8** blockade of probe **3** labelling of recombinant KDM1A in HEK29T cells. **e**, gel (*upper*) and gel-derived curve plot (*lower*) showing dose-dependent competitor **4** blockade of probe **3** labelling of recombinant KDM1A in HEK29T cells. **f**, gel (*upper*) and gel-derived curve plot (*lower*) showing dose-dependent competitor **9** blockade of probe **3** labelling of recombinant KDM1A in HEK29T cells. **g**, gel (*upper*) and gel-derived curve plot (*lower*) showing dose-dependent competitor **10** blockade of probe **3** labelling of recombinant KDM1A in HEK29T cells. **h**, gel (*upper*) and gel-derived curve plot (*lower*) showing dose-dependent competitor **6** blockade of probe **2** labelling of recombinant FTO in HEK29T cells. **i**, gel showing selective competitor **7** blockade of probe **3** labelling of recombinant KDM1A in HEK29T cells.

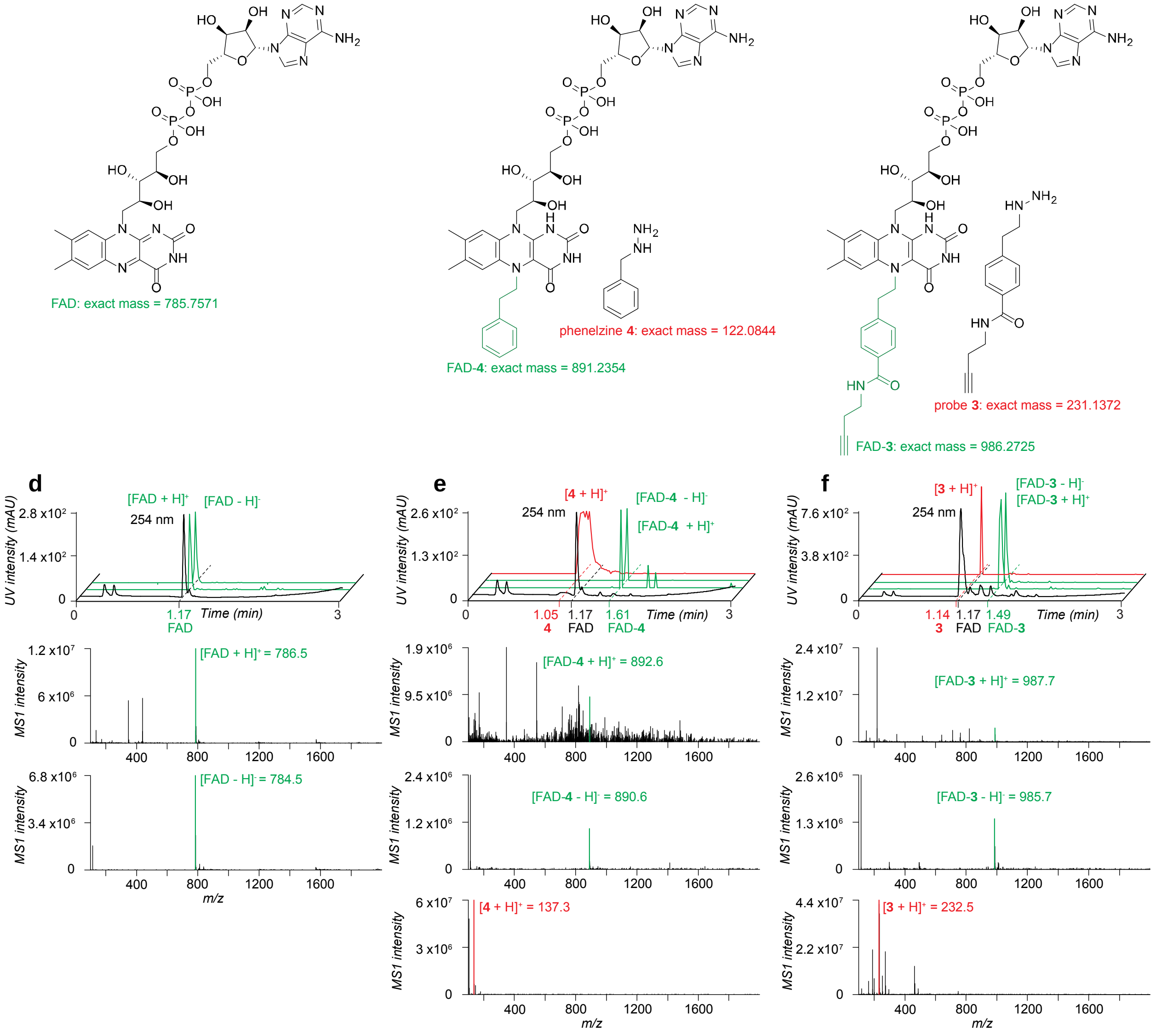

**Supplementary Fig. 5. LC-MS characterization of the FAD-hydrazine adducts in NQO2**.

**a**, structure and theoretical mass of FAD. **b**, structures and theoretical masses of phenelzine **4** and FAD-phenelzine adduct. **c**, structures and theoretical masses of probe **3** and FAD-probe **3** adduct. **d**, LC-MS characterization of FAD including LC chromatogram in **black** (*upper*), EICs generated from positive and negative ionization modes in green (*upper*), and pseudo molecular ions of FAD (green) in positive and negative modes (*middle* and *lower*). **e**, LC-MS characterization of FAD-phenelzine adduct including LC chromatogram in **black** (*upper*), EICs of adduct generated from positive and negative ionization modes in green (*upper*), EIC of phenelzine (red) in positive mode, pseudo molecular ions (green) of adduct in positive and negative modes (*middle*), and pseudo molecular ion of phenelzine (red) in positive mode (*lower*). **f**, LC-MS characterization of FAD-probe **3** adduct including LC chromatogram in **black**

(*upper*), EICs of adduct generated from positive and negative ionization modes in green (*upper*), EIC of probe **3** (red) in positive mode, pseudo molecular ions (green) of adduct in positive and negative modes (*middle*), and pseudo molecular ion of probe **3** (red) in positive mode (*lower*).

**a**

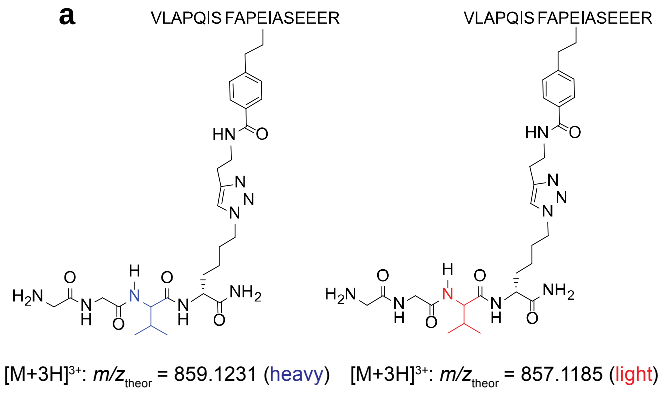

**c**

| | Theoretical *m/z* Heavy | Theoretical *m/z* Light | Measured *m/z* (error in ppm) Heavy | Measured *m/z* (error in ppm) Light |
|---|---|---|---|---|
| $y_{12}$ | 1953.9697 | 1947.9559 | 1953.9684 (-0.7) | 1947.9539 (-1.0) |
| $y_{11}$ | 1866.9377 | 1860.9239 | 1866.9354 (-1.2) | - |
| $y_{10}$ | 1719.8693 | 1713.8555 | 1719.87 (0.4) | 1713.8469 (-5.0) |
| $y_9$ | 1648.8322 | 1642.8184 | 1648.8336 (0.9) | 1642.8124 (-3.7) |
| $y_8$ | 1551.7794 | 1545.7656 | - | 1545.7708 (3.3) |
| $b_2$ | 213.1598 | | 213.1597 (-0.6) | 213.1595 (-1.3) |
| $b_3$ | 284.1969 | | 284.1971 (0.7) | 284.1966 (-1.1) |
| $b_5$ | 509.3082 | | 509.3077 (-0.9) | 509.3092 (2.0) |
| $b_6$ | 622.3923 | | 622.3932 (1.4) | 622.3907 (-2.6) |
| $b_9$ | 927.5298 | | 927.5208 (-9.7) | - |
| $y_7$ | 833.3999 | | 833.4014 (1.8) | 833.3995 (-0.4) |
| $y_6$ | 720.3159 | | 720.3173 (1.9) | 720.3157 (-0.3) |
| $y_5$ | 649.2788 | | 649.2794 (0.9) | 649.2782 (-0.9) |
| $y_4$ | 562.2467 | | 562.2485 (3.2) | 562.2462 (-0.9) |
| $y_3$ | 433.2041 | | 433.2046 (1.1) | 433.2036 (-1.3) |
| $y_2$ | 304.1615 | | 304.1614 (-0.3) | 304.1614 (-0.3) |
| $y_1$ | 175.1190 | | 175.1188 (-1.3) | 175.1187 (-1.5) |

**b**

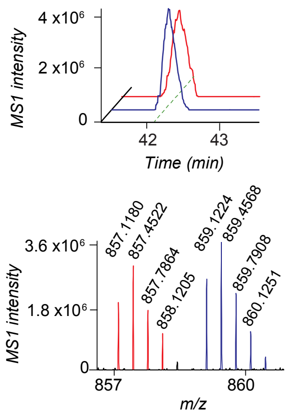

**d**

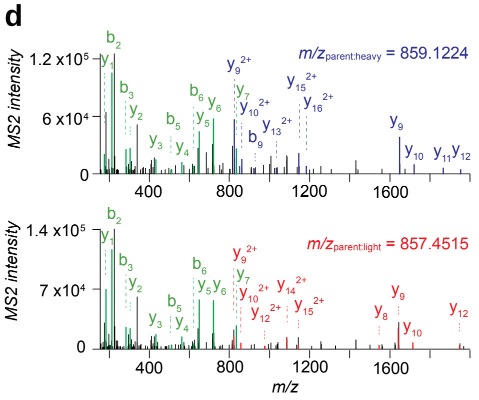

**Supplementary Fig. 6. MS characterization of the probe 3-labelled peptide of NQO2**.

**a**, structures and theoretical parent masses of heavy- and light-tagged NQO2 peptides labelled by probe **3** and processed by the isoTOP-ABPP method. **b**, parent EICs (*upper*) and corresponding isotopic envelopes (*lower*) for heavy- (blue) and light- (red) tagged peptides detected from NQO2-transfected HEK293T cells. **c**, summary table of theoretical versus observed spectra assignments generated under high-resolution MS2 conditions. **d**, MS2 spectra generated from parent ions of **b**. unshifted ions are shown in green and the shifted ion series in blue (heavy) and red (light).

**a**

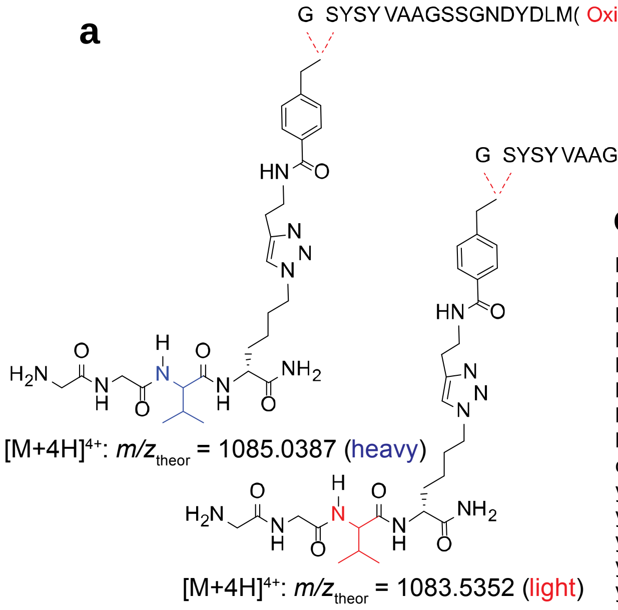

**c**

| | Theoretical $m/z$ Heavy | Theoretical $m/z$ Light | Measured $m/z$ (error in ppm) Heavy | Measured $m/z$ (error in ppm) Light |
|---|---|---|---|---|
| $b_3$ | 897.4610 | 891.4472 | 897.4612 (0.3) | - |
| $b_4$ | 984.4930 | 978.4792 | 984.4922 (-0.8) | 978.4839 (4.8) |
| $b_5$ | 1147.5563 | 1141.5425 | 1147.5529 (-3.0) | 1141.5469 (3.8) |
| $b_6$ | 1246.6247 | 1240.6109 | 1246.6255 (0.6) | 1240.6146 (3.0) |
| $b_7$ | 1317.6619 | 1311.6481 | 1317.6615 (-0.3) | - |
| $b_8$ | 1388.6990 | 1382.6852 | 1388.6963 (-2.0) | 1382.688 (2.0) |
| $b_9$ | 1445.7204 | 1439.7066 | 1445.7186 (-1.2) | 1439.7053 (-0.9) |
| $b_{10}$ | 1532.7525 | 1526.7387 | - | 1526.7521 (8.8) |
| $c_2$ | 751.4243 | 745.4103 | - | 745.4243 (18.7) |
| $y_{17}$ | 1694.9588 | | 1694.9553 (-2.1) | 1694.9612 (1.4) |
| $y_{16}$ | 1597.9061 | | 1597.9083 (1.4) | 1597.901 (-3.2) |
| $y_{15}$ | 1484.8220 | | 1484.8199 (-1.4) | 1484.8269 (3.3) |
| $y_{14}$ | 1383.7743 | | 1383.7734 (-0.6) | 1383.775 (0.5) |
| $y_{13}$ | 1286.7215 | | 1286.7206 (-0.7) | 1286.7234 (1.5) |
| $y_{12}$ | 1229.7001 | | 1229.7002 (0.1) | 1229.7029 (2.3) |
| $y_{11}$ | 1132.6473 | | 1132.6459 (-1.3) | 1132.652 (4.1) |
| $y_{10}$ | 1045.6153 | | 1045.6140 (-1.2) | 1045.6121 (-3.1) |
| $y_9$ | 932.5312 | | 932.5316 (0.4) | 932.5326 (1.5) |
| $y_8$ | 835.4785 | | 835.4794 (1.0) | 835.4791 (0.7) |
| $y_7$ | 778.4570 | | 778.4556 (-1.8) | 778.4557 (-1.7) |
| $y_6$ | 707.4199 | | 707.4202 (0.5) | 707.4206 (1.0) |
| $y_5$ | 610.3671 | | 610.3653 (-3.0) | 610.3622 (-8.0) |
| $y_4$ | 482.3085 | | 482.3087 (0.4) | 482.3085 (0.0) |
| $y_2$ | 272.1717 | | 272.1716 (-0.4) | 272.1715 (-0.8) |
| $y_1$ | 175.1190 | | 175.1188 (-1.4) | 175.1189 (-0.7) |

**b**

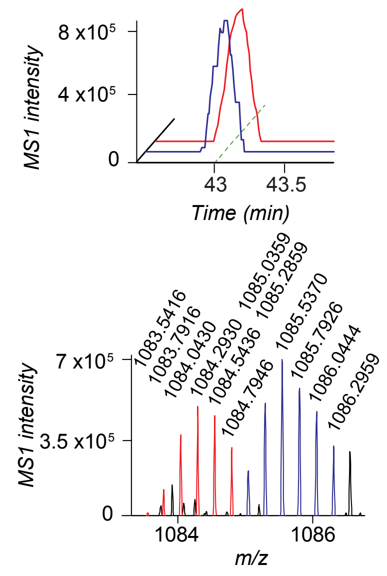

**d**

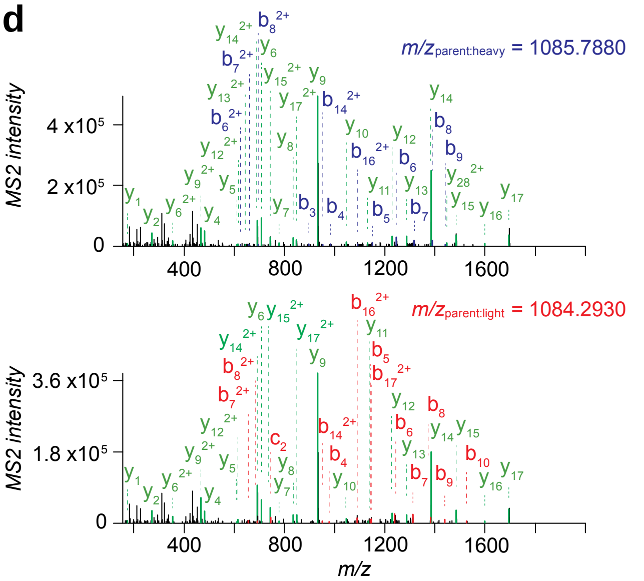

**Supplementary Fig. 7. MS characterization of the probe 3-labelled peptide of KDM1A.**

**a**, structures and theoretical parent masses of heavy- and light-tagged KDM1A peptides labelled by probe **3** and processed by the isoTOP-ABPP method. **b**, parent EICs (*upper*) and corresponding isotopic envelopes (*lower*) for heavy- (blue) and light- (red) tagged peptides detected from KDM1A-transfected HEK293T cells. **c**, summary table of theoretical versus observed spectra assignments generated under high-resolution MS2 conditions. **d**, MS2 spectra generated from parent ions of **b**. unshifted ions are shown in green and the shifted ion series in blue (heavy) and red (light).

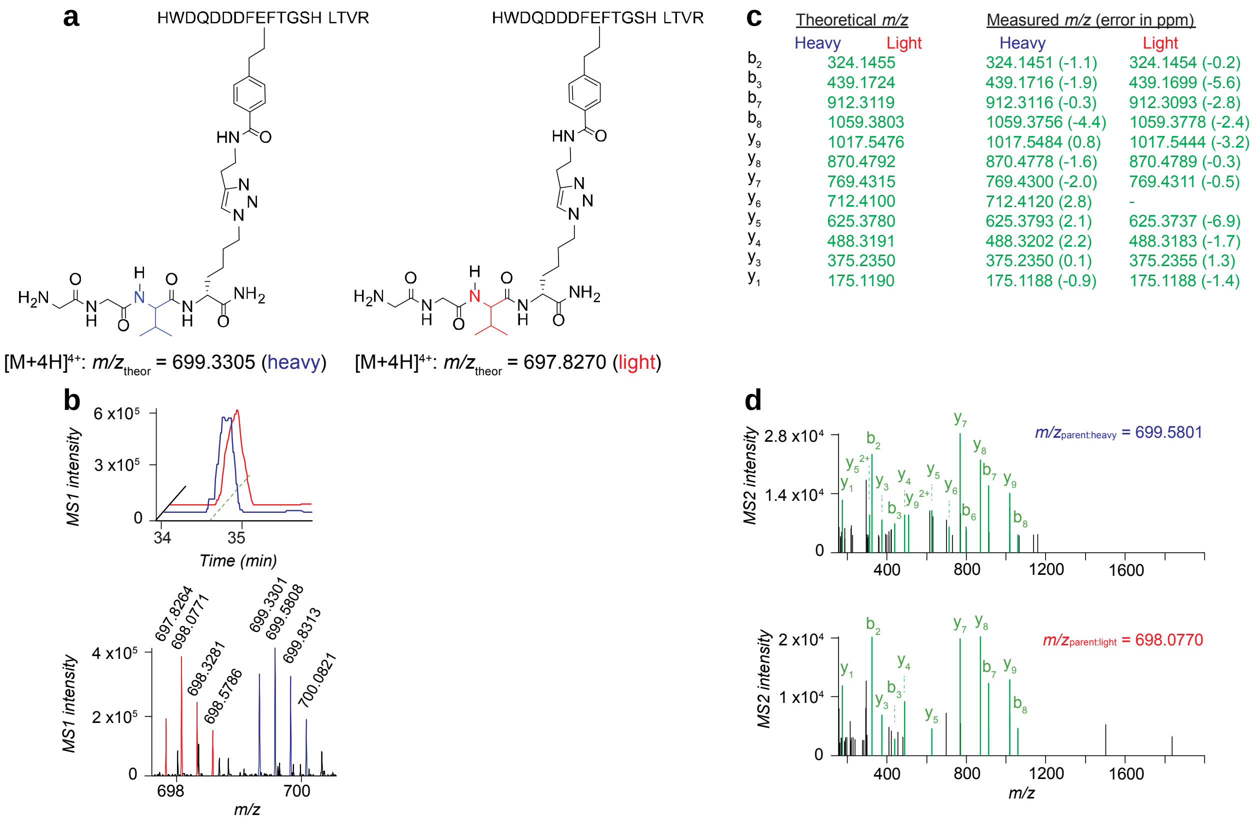

**Supplementary Fig. 8. MS characterization of the probe 3-labelled peptide of KDM1A.**

**a**, structures and theoretical parent masses of heavy- and light-tagged KDM1A peptides labelled by probe **3** and processed by the isoTOP-ABPP method. **b**, parent EICs (*upper*) and corresponding isotopic envelopes (*lower*) for heavy- (blue) and light- (red) tagged peptides detected from KDM1A-transfected HEK293T cells. **c**, summary table of theoretical versus observed spectra assignments generated under high-resolution MS2 conditions. **d**, MS2 spectra generated from parent ions of **b**. unshifted ions are shown in green and the shifted ion series in blue (heavy) and red (light).

a

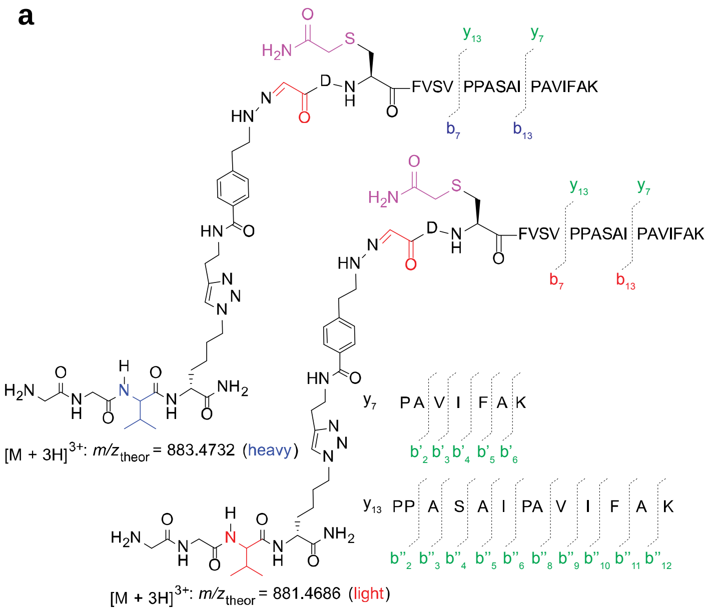

c

| | Theoretical $m/z$ | | Measured $m/z$ (error in ppm) | |
|---|---|---|---|---|
| | Heavy | Light | Heavy | Light |
| $b_2$ | 775.3878 | 769.3740 | 775.3890 (1.5) | 769.3726 (-1.8) |
| $b_3$ | 935.4185 | 929.4047 | 935.4176 (-1.0) | 929.4070 (2.5) |
| $b_4$ | 1082.4869 | 1076.4731 | 1082.4938 (6.4) | 1076.4667 (-5.9) |
| $b_7$ | 1367.6557 | 1361.6419 | - | 1361.6490 (5.2) |
| $b_2$-$NH_3$ | 758.3613 | 752.3475 | 758.3605 (-1.0) | - |
| $b_3$-$NH_3$ | 918.3920 | 912.3782 | 918.3928 (0.9) | 912.3767 (-1.6) |
| $b_4$-$NH_3$ | 1065.4604 | 1059.4466 | 1065.4556 (-4.5) | 1059.4413 (-5.0) |
| $b_6$-$NH_3$ | 1251.5608 | 1245.5470 | 1251.5817 (16.7) | - |
| $b_7$-$NH_3$ | 1350.6292 | 1344.6154 | - | 1344.6321 (12.5) |
| $b_2$-$H_2OCO$ | 713.3636 | 707.3498 | - | 707.4104 (85.7) |
| $b_2$-$H_2O$ | 757.3773 | 751.3634 | 757.3760 (-1.7) | 751.3570 (-8.6) |
| $b_3$-$H_2O$ | 917.4080 | 911.3941 | 917.4141 (6.7) | - |
| $b_6$-$H_2O$ | 1250.5768 | 1244.5629 | 1250.5715 (-4.2) | 1244.5608 (-1.7) |
| $y_{13}$ | 1281.7565 | | 1281.7578 (1.0) | 1281.7574 (0.7) |
| $y_{12}$ | 1184.7038 | | 1184.7096 (4.9) | 1184.7064 (2.2) |
| $y_{11}$ | 1087.6510 | | 1087.6584 (6.8) | 1087.6637 (11.7) |
| $y_{10}$ | 1016.6139 | | 1016.6184 (4.4) | 1016.6169 (3.0) |
| $y_9$ | 929.5819 | | 929.5822 (0.3) | 929.5847 (3.0) |
| $y_8$ | 858.5448 | | 858.5467 (2.2) | 858.5413 (-4.1) |
| $y_7$ | 745.4607 | | 745.4613 (0.8) | 745.4618 (1.5) |
| $y_5$ | 577.3708 | | 577.3694 (-2.4) | 577.3719 (1.9) |
| $y_4$ | 478.3024 | | 478.3042 (3.8) | 478.3048 (5.0) |
| $y_3$ | 365.2183 | | 365.2183 (0.0) | 365.2182 (-0.3) |
| $y_2$ | 218.1499 | | 218.1498 (-0.5) | 218.1498 (-0.5) |
| $y_1$ | 147.1128 | | 147.1128 (1.4) | 147.1128 (0.0) |
| $b''_{12}$ | 1135.6510 | | 1135.6467 (-3.8) | 1135.6643 (11.7) |
| $b''_{11}$ | 1064.6139 | | 1064.6190 (4.8) | 1064.6208 (6.5) |
| $b''_{10}$ | 917.5455 | | 917.5464 (1.0) | 917.5472 (1.9) |
| $b''_9$ | 804.4612 | | 804.4628 (2.0) | 804.4630 (2.2) |
| $b''_8$ | 705.3930 | | 705.3925 (-0.7) | 705.3927 (-0.4) |
| $b''_6$ | 537.3031 | | 537.3035 (0.7) | 537.3038 (1.3) |
| $b''_5$ | 424.2191 | | 424.2196 (1.2) | 424.2194 (0.7) |
| $b''_4$ | 353.1819 | | 353.1820 (0.3) | 353.1815 (-1.1) |
| $b''_3$ | 266.1499 | | 266.1484 (-5.6) | 266.1483 (-6.0) |
| $b''_2$ | 195.1128 | | 195.0877 (-128.6) | 195.0878 (-128.1) |
| $b'_6$ | 599.3552 | | 599.3555 (0.5) | 599.3580 (4.7) |
| $b'_5$ | 528.3180 | | 528.3174 (-1.1) | 528.3183 (0.6) |
| $b'_4$ | 381.2496 | | 381.2494 (-0.5) | 381.2490 (-1.6) |
| $b'_3$ | 268.1656 | | 268.1656 (0.0) | 268.1653 (-1.1) |
| $b'_2$ | 169.0972 | | 169.0974 (1.2) | 169.0974 (1.2) |

b

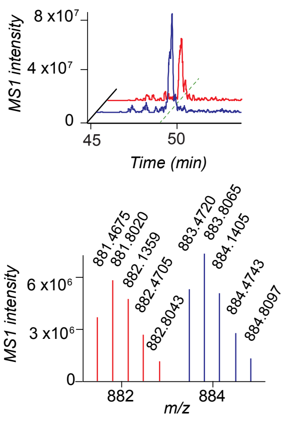

d

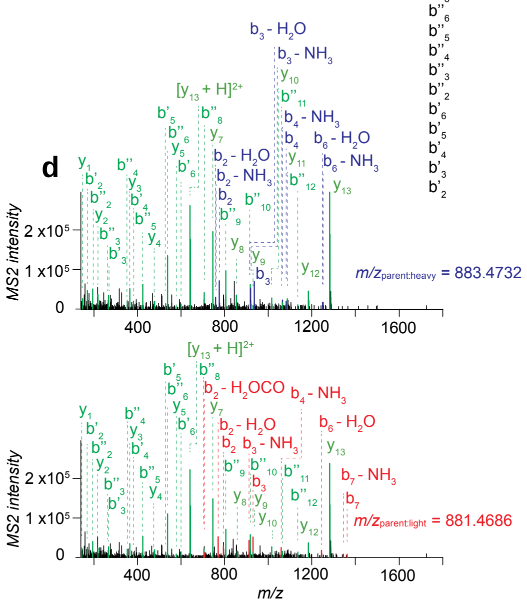

**Supplementary Fig. 9. MS characterization of the probe 3-labelled glyoxylyl group of SCRN2.**

**a**, structures and theoretical parent masses of heavy- and light-tagged SCRN2 glyoxylyl peptides labelled by probe **3** and processed by the isoTOP-ABPP method. **b**, parent EICs (*upper*) and corresponding isotopic envelopes (*lower*) for heavy- (blue) and light- (red) tagged peptides detected from SCRN2-transfected HEK293T cells. **c**, summary table of theoretical versus observed spectra assignments generated under high-resolution MS2 conditions. **d**, MS2 spectra generated from parent ions of **b**. unshifted ions are shown in green and the shifted ion series in blue (heavy) and red (light).

**a**

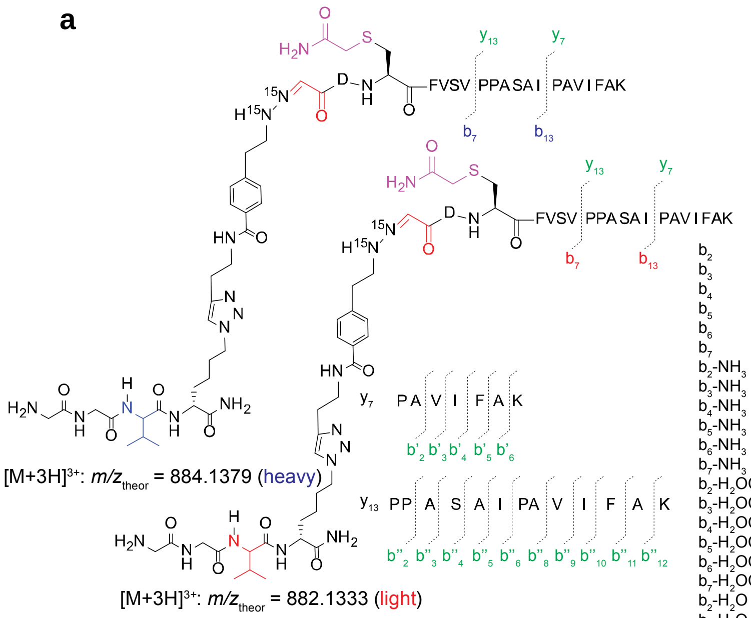

**b**

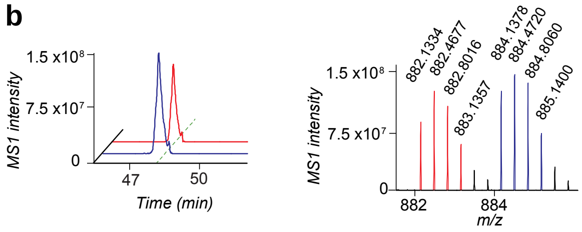

**c**

| | Theoretical *m/z* Heavy | Theoretical *m/z* Light | Measured *m/z* (error in ppm) Heavy | Measured *m/z* (error in ppm) Light |
|---|---|---|---|---|
| $b_2$ | 777.3819 | 771.3681 | 777.3815 (-0.5) | 771.3689 (1.0) |
| $b_3$ | 937.4125 | 931.3987 | 937.4125 (0.0) | 931.3994 (0.8) |
| $b_4$ | 1084.4809 | 1078.4671 | 1084.4773 (-3.3) | 1078.4674 (0.3) |
| $b_5$ | 1183.5493 | 1177.5355 | - | 1177.5337 (-1.5) |
| $b_6$ | 1270.5814 | 1264.5676 | - | 1264.5659 (-1.3) |
| $b_7$ | 1369.6498 | 1363.6360 | 1369.6494 (-0.3) | 1363.6371 (0.8) |
| $b_2$-$NH_3$ | 760.3554 | 754.3416 | 760.3586 (4.3) | 754.3444 (3.8) |
| $b_3$-$NH_3$ | 920.3860 | 914.3722 | 920.3892 (3.5) | 914.3735 (1.5) |
| $b_4$-$NH_3$ | 1067.4544 | 1061.4406 | 1067.4548 (0.4) | 1061.4429 (2.2) |
| $b_5$-$NH_3$ | 1166.5228 | 1160.5090 | 1166.5192 (-3.1) | 1160.5114 (2.1) |
| $b_6$-$NH_3$ | 1253.5549 | 1247.5411 | 1253.5673 (9.9) | 1247.5562 (12.1) |
| $b_7$-$NH_3$ | 1352.6233 | 1346.6095 | 1352.6260 (2.0) | 1346.6150 (4.1) |
| $b_2$-$H_2OCO$ | 715.3577 | 709.3439 | - | 709.3367 (-10.2) |
| $b_3$-$H_2OCO$ | 875.3883 | 869.3745 | - | 869.3571 (-20.0) |
| $b_4$-$H_2OCO$ | 1022.4567 | 1016.4429 | - | 1016.4225 (-20.1) |
| $b_5$-$H_2OCO$ | 1121.5251 | 1115.5113 | - | 1115.4958 (-13.9) |
| $b_6$-$H_2OCO$ | 1208.5572 | 1202.5434 | - | 1202.5409 (-2.1) |
| $b_7$-$H_2OCO$ | 1307.6256 | 1301.6118 | - | 1301.6062 (-4.3) |
| $b_2$-$H_2O$ | 759.3714 | 753.3575 | 759.3712 (-0.2) | 753.3585 (1.3) |
| $b_3$-$H_2O$ | 919.4020 | 913.3881 | 919.4012 (-0.8) | 913.3865 (-1.8) |
| $b_4$-$H_2O$ | 1066.4704 | 1060.4565 | 1066.4725 (2.0) | 1060.4584 (1.8) |
| $b_5$-$H_2O$ | 1165.5388 | 1159.5249 | - | 1159.5288 (3.3) |
| $b_6$-$H_2O$ | 1252.5709 | 1246.5570 | 1252.5709 (0.0) | 1246.5576 (0.4) |
| $b_7$-$H_2O$ | 1351.6393 | 1345.6254 | 1351.6442 (3.7) | 1345.6256 (0.1) |
| $y_{15}$ | 1467.8570 | | 1467.8489 (-5.5) | 1467.8508 (-4.2) |
| $y_{14}$ | 1380.8249 | | - | 1380.8328 (5.7) |
| $y_{13}$ | 1281.7565 | | 1281.7565 (0.0) | 1281.7572 (0.5) |
| $y_{12}$ | 1184.7038 | | 1184.7025 (-1.1) | 1184.7045 (0.6) |
| $y_{11}$ | 1087.6510 | | 1087.6538 (2.6) | 1087.6511 (0.1) |
| $y_{10}$ | 1016.6139 | | 1016.6140 (0.1) | 1016.6147 (0.8) |
| $y_9$ | 929.5819 | | 929.5803 (-1.7) | 929.5821 (0.2) |
| y8 | 858.5448 | | 858.5447 (-0.1) | 858.5456 (0.9) |
| y7 | 745.4607 | | 745.4611 (0.5) | 745.4613 (0.8) |
| y5 | 577.3708 | | 577.3740 (5.5) | 577.3720 (2.1) |
| y4 | 478.3024 | | 478.3024 (0.0) | 478.3023 (-0.2) |
| y3 | 365.2183 | | 365.2184 (0.3) | 365.2183 (0.0) |
| y2 | 218.1499 | | 218.1495 (-1.8) | 218.1497 (-0.9) |
| $b''_{12}$ | 1135.6510 | | 1135.6473 (-3.3) | 1135.6515 (0.4) |
| $b''_{11}$ | 1064.6139 | | 1064.6149 (0.9) | 1064.6149 (0.9) |
| $b''_{10}$ | 917.5455 | | 917.5458 (0.3) | 917.5462 (0.8) |
| $b''_9$ | 804.4612 | | 804.4614 (0.2) | 804.4619 (0.9) |
| $b''_8$ | 705.3930 | | 705.3926 (-0.6) | 705.3939 (1.3) |
| $b''_7$ | 634.3559 | | - | 634.3561 (0.3) |
| $b''_6$ | 537.3031 | | 537.3032 (0.2) | 537.3033 (0.4) |
| $b''_5$ | 424.2191 | | 424.2186 (-1.2) | 424.2192 (0.2) |
| $b''_4$ | 353.1819 | | 353.1818 (-0.3) | 353.1820 (0.3) |
| $b''_3$ | 266.1499 | | 266.1498 (-0.4) | 266.1498 (-0.4) |
| $b''_2$ | 195.1128 | | 195.1129 (0.5) | 195.1125 (-1.5) |
| $b'_6$ | 599.3552 | | 599.3562 (1.7) | 599.3558 (1.0) |
| $b'_5$ | 528.3180 | | 528.3177 (-0.6) | 528.3197 (3.2) |
| $b'_4$ | 381.2496 | | - | 381.2496 (0.0) |
| $b'_3$ | 268.1656 | | 268.1656 (0.0) | 268.1656 (0.0) |
| $b'_2$ | 169.0972 | | 169.0970 (-1.2) | 169.0970 (-1.2) |

**d**

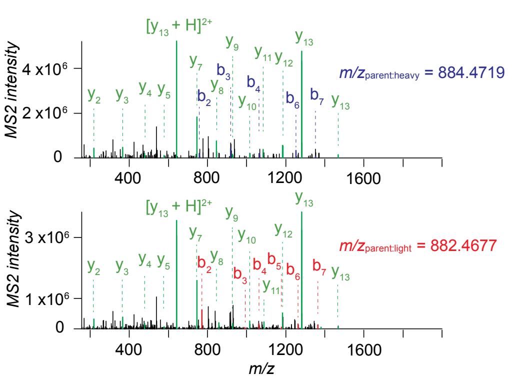

**Supplementary Fig. 10. MS characterization of the probe $^{15}N_2$-3 labelled glyoxylyl group of SCRN2.**

**a**, structures and theoretical parent masses of heavy- and light-tagged SCRN2 glyoxylyl peptides labelled by probe **$^{15}N_2$-3** and processed by the isoTOP-ABPP method. **b**, parent EICs (*left*) and corresponding isotopic envelopes (*right*) for heavy- (blue) and light- (red) tagged peptides detected from SCRN2-transfected HEK293T cells. **c**, summary table of theoretical versus observed spectra assignments generated under high-resolution MS2 conditions. **d**, MS2 spectra generated from parent ions of **b**. unshifted ions are shown in green and the shifted ion series in blue (heavy) and red (light).

**a**

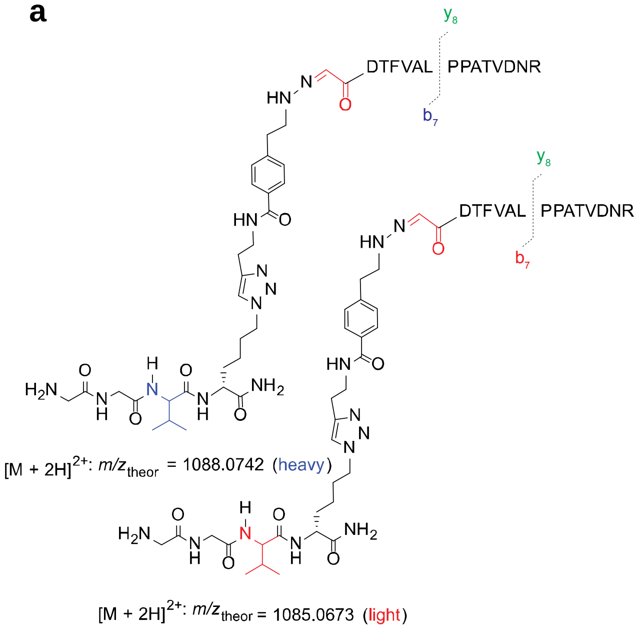

**c**

| | Theoretical *m/z* Heavy | Theoretical *m/z* Light | Measured *m/z* (error in Da) Heavy | Measured *m/z* (error in Da) Light |
|---|---|---|---|---|
| $b_2$ | 775.39 | 769.37 | 775.62 (0.23) | 769.45 (0.08) |
| $b_3$ | 876.44 | 870.42 | 876.43 (0.00) | 870.54 (0.12) |
| $b_4$ | 1023.50 | 1017.49 | 1023.87 (0.37) | - |
| $b_5$ | 1122.57 | 1116.56 | 1122.63 (0.05) | 1116.76 (0.20) |
| $b_6$ | 1193.61 | 1187.60 | 1193.66 (0.05) | 1187.76 (0.16) |
| $b_7$ | 1306.69 | 1300.68 | 1306.82 (0.12) | 1300.88 (0.20) |
| $b_8$ | 1403.75 | 1397.73 | 1403.94 (0.19) | 1398.09 (0.36) |
| $b_9$ | 1500.80 | 1494.79 | 1500.90 (0.10) | 1494.98 (0.19) |
| $b_{10}$ | 1571.84 | 1565.82 | 1571.86 (0.02) | 1565.78 (-0.04) |
| $b_{11}$ | 1672.88 | 1666.87 | 1672.60 (-0.28) | 1666.56 (-0.31) |
| $b_{12}$ | 1771.95 | 1765.94 | - | - |
| $b_{13}$ | 1886.98 | 1880.97 | 1887.23 (0.25) | 1880.98 (0.02) |
| $b_{14}$ | 2001.02 | 1995.01 | - | 1995.07 (0.06) |
| $b_7$ - $NH_3$ | 1289.67 | 1283.65 | 1289.75 (0.09) | 1283.80 (0.14) |
| $b_7$ - $NH_3CO$ | 1261.67 | 1255.66 | 1261.67 (0.00) | 1255.81 (0.15) |
| $b_7$ - $H_2O$ | 1288.68 | 1282.67 | 1288.95 (0.27) | 1282.77 (0.10) |
| $y_{14}$ | 1515.78 | | 1515.82 (0.04) | 1515.70 (-0.09) |
| $y_{13}$ | 1400.75 | | 1400.80 (0.05) | 1400.81 (0.06) |
| $y_{12}$ | 1299.71 | | 1299.84 (0.13) | 1299.60 (-0.11) |
| $y_{11}$ | 1152.64 | | 1152.65 (0.01) | 1152.67 (0.03) |
| $y_{10}$ | 1053.57 | | 1053.57 (0.00) | 1053.61 (0.04) |
| $y_9$ | 982.53 | | 982.57 (0.04) | 982.57 (0.04) |
| $y_8$ | 869.45 | | 869.45 (0.00) | 869.46 (0.01) |
| $y_7$ | 772.39 | | 772.42 (0.02) | 772.42 (0.02) |
| $y_6$ | 675.34 | | 675.46 (0.12) | 675.41 (0.07) |
| $y_5$ | 604.30 | | 604.37 (0.07) | 604.30 (0.00) |
| $y_4$ | 503.26 | | 503.28 (0.02) | 503.25 (0.00) |
| $y_3$ | 404.19 | | 404.31 (0.12) | 404.21 (0.02) |
| $y_2$ | 289.16 | | 289.22 (0.06) | 289.10 (-0.06) |
| $y_1$ | 175.12 | | 175.18 (0.06) | 175.18 (0.07) |

**b**

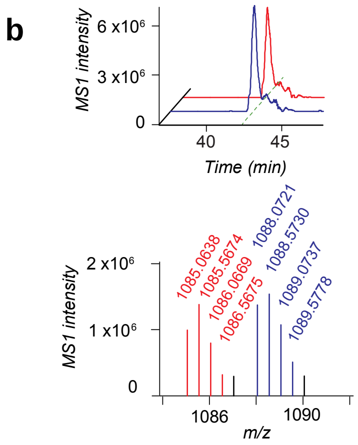

**d**

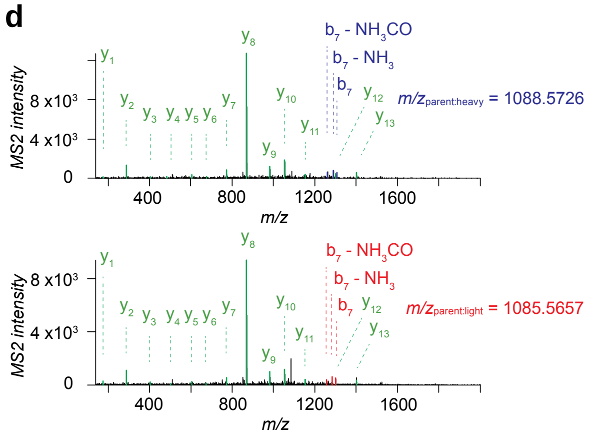

**Supplementary Fig. 11. MS characterization of the probe 3-labelled glyoxylyl group of SCRN3**.

**a**, structures and theoretical parent masses of heavy- and light-tagged SCRN3 glyoxylyl peptides labelled by probe **3** and processed by the isoTOP-ABPP method. **b**, parent EICs (*upper*) and corresponding isotopic envelopes (*lower*) for heavy- (blue) and light- (red) tagged peptides detected from SCRN3-transfected HEK293T cells. **c**, summary table of theoretical versus observed spectra assignments generated under low-resolution MS2 conditions. **d**, MS2 spectra generated from parent ions of **b**. unshifted ions are shown in green and the shifted ion series in blue (heavy) and red (light).

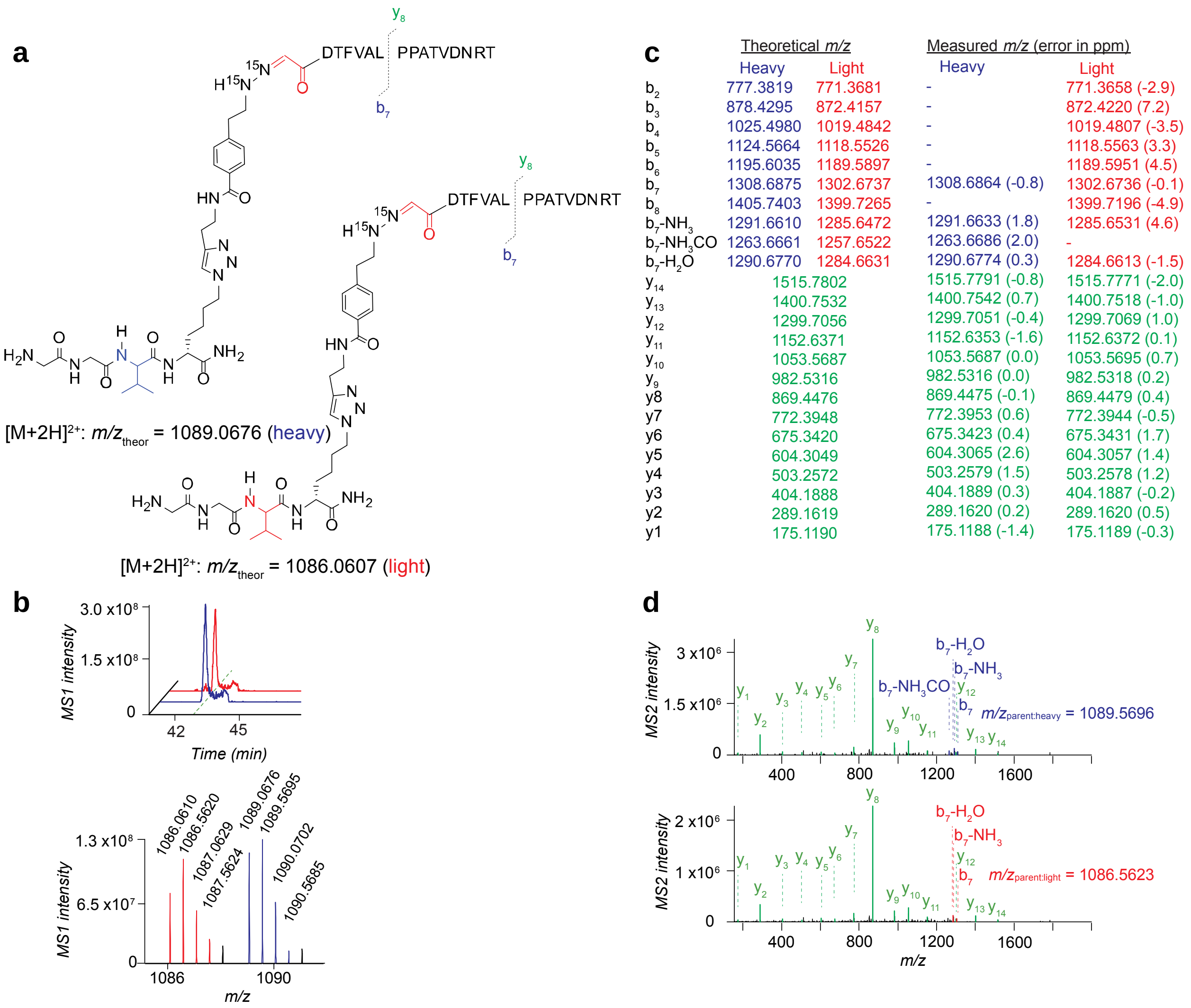

| | Theoretical *m/z* Heavy | Theoretical *m/z* Light | Measured *m/z* (error in ppm) Heavy | Measured *m/z* (error in ppm) Light |
|---|---|---|---|---|
| $b_2$ | 777.3819 | 771.3681 | - | 771.3658 (-2.9) |
| $b_3$ | 878.4295 | 872.4157 | - | 872.4220 (7.2) |
| $b_4$ | 1025.4980 | 1019.4842 | - | 1019.4807 (-3.5) |
| $b_5$ | 1124.5664 | 1118.5526 | - | 1118.5563 (3.3) |
| $b_6$ | 1195.6035 | 1189.5897 | - | 1189.5951 (4.5) |
| $b_7$ | 1308.6875 | 1302.6737 | 1308.6864 (-0.8) | 1302.6736 (-0.1) |
| $b_8$ | 1405.7403 | 1399.7265 | - | 1399.7196 (-4.9) |
| $b_7$-$NH_3$ | 1291.6610 | 1285.6472 | 1291.6633 (1.8) | 1285.6531 (4.6) |
| $b_7$-$NH_3CO$ | 1263.6661 | 1257.6522 | 1263.6686 (2.0) | - |
| $b_7$-$H_2O$ | 1290.6770 | 1284.6631 | 1290.6774 (0.3) | 1284.6613 (-1.5) |
| $y_{14}$ | 1515.7802 | | 1515.7791 (-0.8) | 1515.7771 (-2.0) |
| $y_{13}$ | 1400.7532 | | 1400.7542 (0.7) | 1400.7518 (-1.0) |
| $y_{12}$ | 1299.7056 | | 1299.7051 (-0.4) | 1299.7069 (1.0) |
| $y_{11}$ | 1152.6371 | | 1152.6353 (-1.6) | 1152.6372 (0.1) |
| $y_{10}$ | 1053.5687 | | 1053.5687 (0.0) | 1053.5695 (0.7) |
| $y_9$ | 982.5316 | | 982.5316 (0.0) | 982.5318 (0.2) |
| y8 | 869.4476 | | 869.4475 (-0.1) | 869.4479 (0.4) |
| y7 | 772.3948 | | 772.3953 (0.6) | 772.3944 (-0.5) |
| y6 | 675.3420 | | 675.3423 (0.4) | 675.3431 (1.7) |
| y5 | 604.3049 | | 604.3065 (2.6) | 604.3057 (1.4) |
| y4 | 503.2572 | | 503.2579 (1.5) | 503.2578 (1.2) |
| y3 | 404.1888 | | 404.1889 (0.3) | 404.1887 (-0.2) |
| y2 | 289.1619 | | 289.1620 (0.2) | 289.1620 (0.5) |
| y1 | 175.1190 | | 175.1188 (-1.4) | 175.1189 (-0.3) |

**Supplementary Fig. 12. MS characterization of the probe $^{15}N_2$-3 labelled glyoxylyl group of SCRN3**.

**a**, structures and theoretical parent masses of heavy- and light-tagged SCRN3 glyoxylyl peptides labelled by probe **$^{15}N_2$-3** and processed by the isoTOP-ABPP method. **b**, parent EICs (*upper*) and corresponding isotopic envelopes (*lower*) for heavy- (blue) and light- (red) tagged peptides detected from SCRN3-transfected HEK293T cells. **c**, summary table of theoretical versus observed spectra assignments generated under high-resolution MS2 conditions. **d**, MS2 spectra generated from parent ions of **b**. unshifted ions are shown in green and the shifted ion series in blue (heavy) and red (light).

**a**

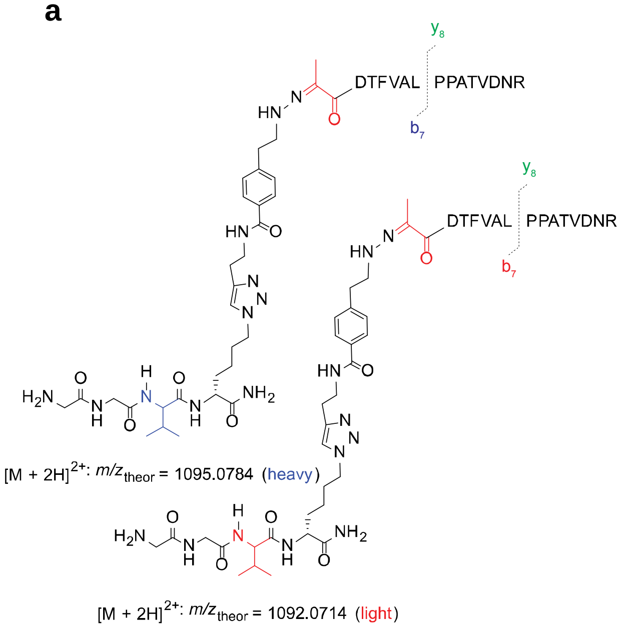

**c**

| | Theoretical *m/z* Heavy | Theoretical *m/z* Light | Measured *m/z* (error in Da) Heavy | Measured *m/z* (error in Da) Light |
|---|---|---|---|---|
| $b_2$ | 789.40 | 783.39 | 789.58 (0.17) | 783.50 (0.11) |
| $b_3$ | 890.45 | 884.44 | 890.31 (-0.14) | 884.64 (0.20) |
| $b_4$ | 1037.52 | 1031.51 | 1037.82 (0.3) | 1031.10 (-0.40) |
| $b_5$ | 1136.59 | 1130.57 | 1136.75 (0.16) | 1130.87 (0.29) |
| $b_6$ | 1207.63 | 1201.61 | 1207.90 (0.28) | 1201.79 (0.17) |
| $b_7$ | 1320.71 | 1314.70 | 1320.86 (0.15) | 1315 (0.31) |
| $b_8$ | 1417.76 | 1411.75 | 1417.94 (0.18) | 1412.18 (0.43) |
| $b_9$ | 1514.81 | 1508.80 | - | 1508.24 (-0.57) |
| $b_{10}$ | 1585.85 | 1579.84 | 1585.19 (-0.67) | 1579.85 (0.02) |
| $b_{11}$ | 1686.90 | 1680.89 | - | - |
| $b_{12}$ | 1785.97 | 1779.95 | - | 1780.41 (0.46) |
| $b_{13}$ | 1900.99 | 1894.98 | 1900.85 (-0.15) | - |
| $b_7$ - $NH_3$ | 1303.68 | 1297.67 | 1303.87 (0.19) | 1297.83 (0.16) |
| $b_7$ - $NH_3CO$ | 1275.69 | 1269.67 | 1275.79 (0.10) | 1269.83 (0.15) |
| $b_7$ - $H_2O$ | 1302.70 | 1296.68 | 1302.77 (0.07) | 1296.87 (0.18) |
| $y_{14}$ | 1515.78 | | 1515.94 (0.16) | 1515.79 (0.00) |
| $y_{13}$ | 1400.75 | | 1400.92 (0.17) | 1400.89 (0.14) |
| $y_{12}$ | 1299.71 | | 1299.89 (0.19) | 1299.89 (0.19) |
| $y_{11}$ | 1152.64 | | 1152.80 (0.16) | 1152.95 (0.31) |
| $y_{10}$ | 1053.57 | | 1053.69 (0.12) | 1053.75 (0.18) |
| $y_9$ | 982.53 | | 982.62 (0.09) | 982.65 (0.12) |
| $y_8$ | 869.45 | | 869.58 (0.13) | 869.56 (0.11) |
| $y_7$ | 772.39 | | 772.53 (0.14) | 772.46 (0.07) |
| $y_6$ | 675.34 | | 675.48 (0.14) | 675.48 (0.14) |
| $y_5$ | 604.30 | | 604.42 (0.12) | 604.71 (0.41) |
| $y_4$ | 503.26 | | 503.46 (0.20) | 503.26 (0.00) |
| $y_3$ | 404.19 | | 404.31 (0.12) | 404.21 (0.02) |
| $y_2$ | 289.16 | | 289.24 (0.08) | 289.17 (0.00) |
| $y_1$ | 175.12 | | 175.11 (-0.01) | 175.13 (0.01) |

**b**

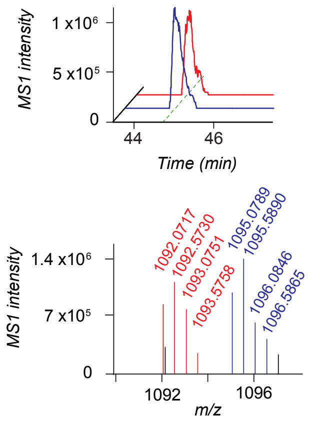

**d**

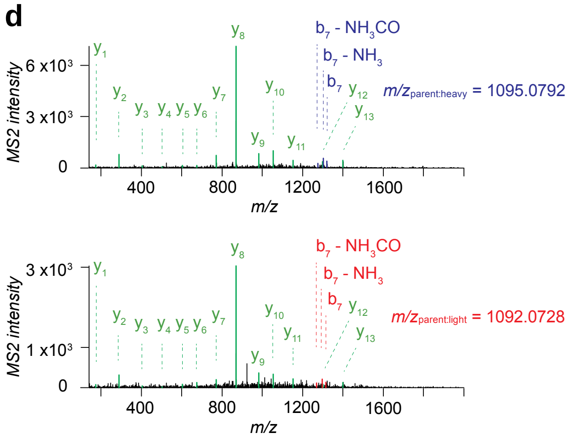

**Supplementary Fig. 13. MS characterization of the probe 3-labelled pyruvoyl group of SCRN3**.

**a**, structures and theoretical parent masses of heavy- and light-tagged SCRN3 pyruvoyl peptides labelled by probe **3** and processed by the isoTOP-ABPP method. **b**, parent EICs (*upper*) and corresponding isotopic envelopes (*lower*) for heavy- (blue) and light- (red) tagged peptides detected from SCRN3-transfected HEK293T cells. **c**, summary table of theoretical versus observed spectra assignments generated under low-resolution MS2 conditions. **d**, MS2 spectra generated from parent ions of **b**. unshifted ions are shown in green and the shifted ion series in blue (heavy) and red (light).

**a**

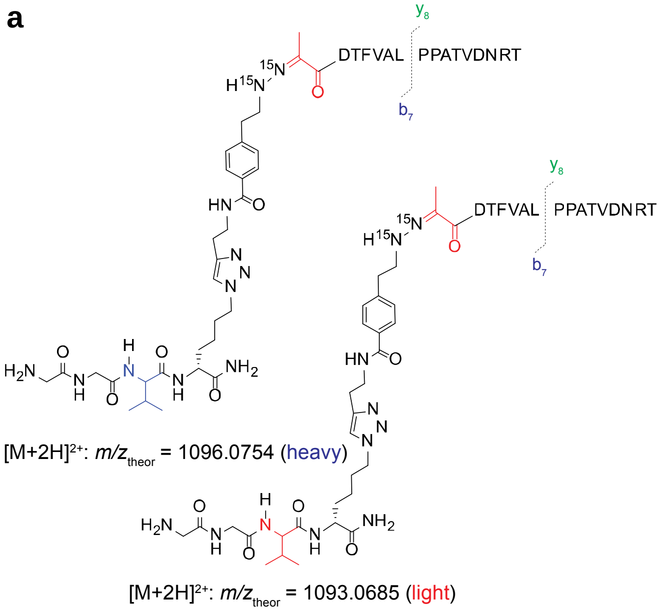

**c**

| | Theoretical *m/z* Heavy | Theoretical *m/z* Light | Measured *m/z* (error in ppm) Heavy | Measured *m/z* (error in ppm) Light |
|---|---|---|---|---|
| $b_2$ | 791.3975 | 785.3837 | 791.3964 (-1.5) | 785.3903 (8.4) |
| $b_5$ | 1138.5820 | 1132.5682 | - | 1132.5427 (-22.5) |
| $b_7$ | 1322.7032 | 1316.6894 | 1322.7012 (-1.5) | 1316.6914 (1.5) |
| $b_7$-$NH_3$ | 1305.6767 | 1299.6629 | - | 1299.6803 (13.4) |
| $b_7$-$NH_3$CO | 1277.6818 | 1271.6679 | 1277.6881 (5.0) | 1271.6743 (5.0) |
| $b_7$-$H_2O$ | 1304.6927 | 1298.6788 | 1304.6796 (-10.0) | - |
| $y_{14}$ | 1515.7802 | | - | 1515.7429 (-24.6) |
| $y_{13}$ | 1400.7532 | | 1400.7487 (-3.2) | 1400.7539 (0.5) |
| $y_{12}$ | 1299.7056 | | 1299.704 (-1.2) | 1299.6803 (-19.5) |
| $y_{11}$ | 1152.6371 | | 1152.6338 (-2.9) | 1152.625 (-10.5) |
| $y_{10}$ | 1053.5687 | | 1053.5677 (-0.9) | 1053.567 (-1.6) |
| $y_9$ | 982.5316 | | 982.5303 (-1.4) | 982.5314 (-0.2) |
| y8 | 869.4476 | | 869.4471 (-0.5) | 869.4499 (2.6) |
| y7 | 772.3948 | | 772.3945 (-0.3) | 772.3942 (-0.8) |
| y6 | 675.3420 | | 675.3384 (-5.3) | 675.3364 (-8.3) |
| y5 | 604.3049 | | 604.3046 (-0.6) | 604.3073 (4.0) |
| y4 | 503.2572 | | - | 503.2622 (9.9) |
| y3 | 404.1888 | | 404.188 (-2.0) | 404.1881 (-1.6) |
| y2 | 289.1619 | | 289.1619 (0.0) | 289.1624 (1.9) |
| y1 | 175.1190 | | 175.1188 (-1.0) | 175.1197 (3.8) |

**b**

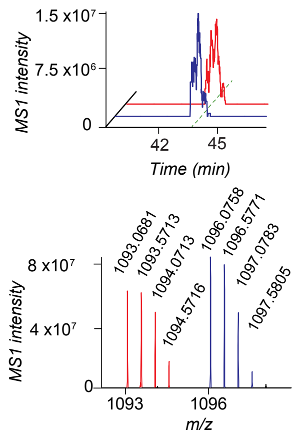

**d**

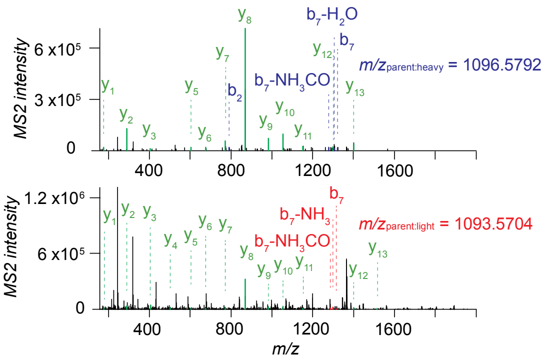

**Supplementary Fig. 14. MS characterization of the probe $^{15}N_2$-3 labelled pyruvoyl group of SCRN3.**

**a**, structures and theoretical parent masses of heavy- and light-tagged SCRN3 pyruvoyl peptides labelled by probe **$^{15}N_2$-3** and processed by the isoTOP-ABPP method. **b**, parent EICs (*upper*) and corresponding isotopic envelopes (*lower*) for heavy- (blue) and light- (red) tagged peptides detected from SCRN3-transfected HEK293T cells. **c**, summary table of theoretical versus observed spectra assignments generated under high-resolution MS2 conditions. **d**, MS2 spectra generated from parent ions of **b**. unshifted ions are shown in green and the shifted ion series in blue (heavy) and red (light).

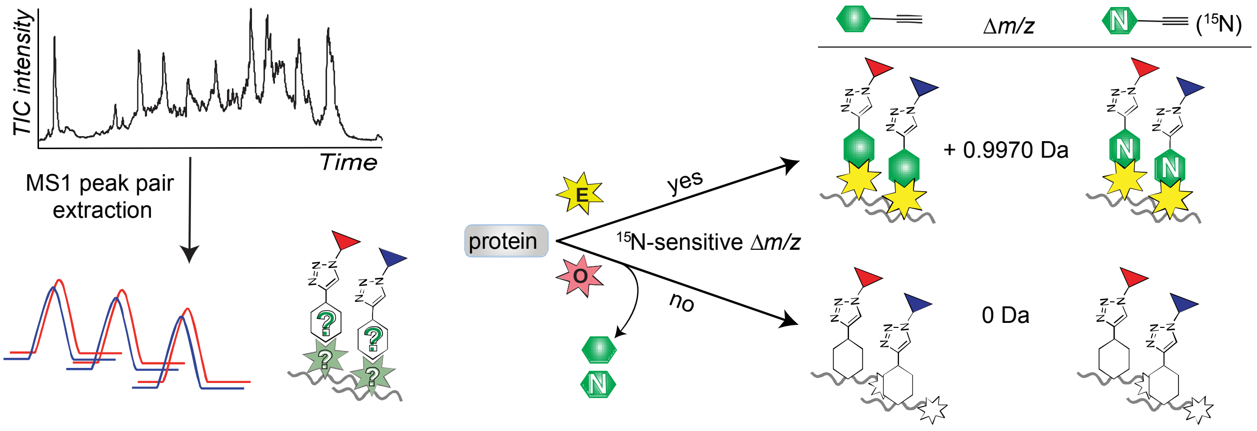

**Supplementary Fig. 15. Workflow for characterization of isotopic probes-labelled peptides.**

Experimental strategy to elucidate probe labelling mechanism using isotopic probes. MS1 peak pairs from isoTOP-ABPP can be extracted by computational script, but the covalent labelling mechanism of probe-protein interaction remains unknown (*left*). A strategy to unveil the labelling mechanism is to use isotopic probes for isoTOP-ABPP (*right*). Peptide pairs labelled by isotopic probes show a mass difference of 0.9970 when hydrazine warheads are retained or show no mass difference when hydrazine warheads are lost.

**a**

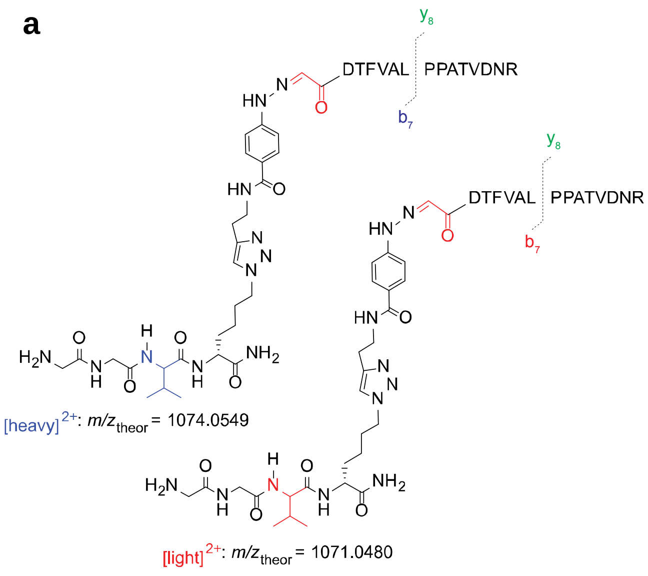

**c**

| | Theoretical *m/z* Heavy | Theoretical *m/z* Light | Measured *m/z* (error in Da) Heavy | Measured *m/z* (error in Da) Light |
|---|---|---|---|---|
| $b_2$ | 747.36 | 741.34 | 747.50 (0.14) | 741.46 (0.12) |
| $b_3$ | 848.40 | 842.39 | 848.52 (0.11) | 842.44 (0.05) |
| $b_4$ | 995.47 | 989.46 | 995.56 (0.09) | 989.55 (0.09) |
| $b_5$ | 1094.54 | 1088.53 | 1094.59 (0.05) | 1088.56 (0.04) |
| $b_6$ | 1165.58 | 1159.56 | 1165.59 (0.01) | 1159.63 (0.07) |
| $b_7$ | 1278.66 | 1272.65 | 1278.70 (0.04) | 1272.67 (0.02) |
| $b_8$ | 1375.71 | 1369.70 | 1375.79 (0.07) | 1369.74 (0.04) |
| $b_{11}$ | 1644.85 | 1638.84 | 1644.87 (0.01) | 1638.88 (0.04) |
| $b_{12}$ | 1743.92 | 1737.91 | 1743.95 (0.03) | 1737.78 (-0.13) |
| $b_{13}$ | 1858.95 | 1852.93 | 1858.85 (-0.10) | 1853.05 (0.12) |
| $b_{14}$ | 1972.99 | 1966.98 | 1972.92 (-0.07) | - |
| $b_7$ - $NH_3$ | 1261.64 | 1255.62 | 1261.69 (0.05) | 1255.67 (0.04) |
| $b_7$ - $NH_3CO$ | 1233.64 | 1227.63 | 1233.68 (0.04) | 1227.65 (0.02) |
| $b_7$ - $H_2O$ | 1260.65 | 1254.64 | 1260.82 (0.17) | 1254.62 (-0.02) |
| $y_{14}$ | 1515.78 | | 1515.82 (0.04) | 1515.82 (0.04) |
| $y_{13}$ | 1400.75 | | 1400.80 (0.04) | 1400.80 (0.04) |
| $y_{12}$ | 1299.71 | | 1299.75 (0.05) | 1299.75 (0.05) |
| $y_{11}$ | 1152.64 | | 1152.68 (0.04) | 1152.70 (0.06) |
| $y_{10}$ | 1053.57 | | 1053.62 (0.05) | 1053.62 (0.05) |
| $y_9$ | 982.53 | | 982.54 (0.01) | 982.58 (0.05) |
| $y_8$ | 869.45 | | 869.55 (0.10) | 869.51 (0.06) |
| $y_7$ | 772.39 | | 772.47 (0.07) | 772.49 (0.10) |
| $y_6$ | 675.34 | | 675.45 (0.11) | 675.48 (0.14) |
| $y_5$ | 604.30 | | 604.45 (0.14) | 604.36 (0.06) |
| $y_4$ | 503.26 | | 503.34 (0.08) | 503.33 (0.07) |
| $y_3$ | 404.19 | | 404.29 (0.10) | 404.26 (0.07) |
| $y_2$ | 289.16 | | 289.25 (0.09) | 289.23 (0.06) |
| $y_1$ | 175.12 | | 175.16 (0.04) | 175.15 (0.03) |

**b**

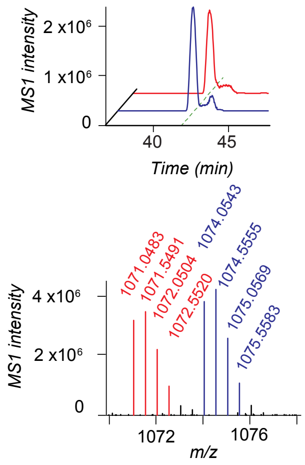

**d**

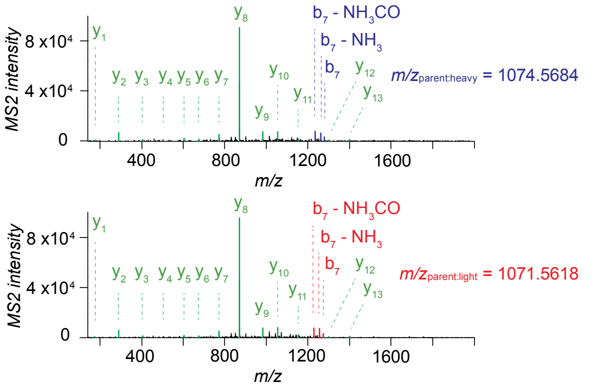

**Supplementary Fig. 16. MS characterization of the probe 2-labelled glyoxylyl group of SCRN3**.

**a**, structures and theoretical parent masses of heavy- and light-tagged SCRN3 glyoxylyl peptides labelled by probe **2** and processed by the isoTOP-ABPP method. **b**, parent EICs (*upper*) and corresponding isotopic envelopes (*lower*) for heavy- (blue) and light- (red) tagged peptides detected from SCRN3-transfected HEK293T cells. **c**, summary table of theoretical versus observed spectra assignments generated under low-resolution MS2 conditions. **d**, MS2 spectra generated from parent ions of **b**. unshifted ions are shown in green and the shifted ion series in blue (heavy) and red (light).

a

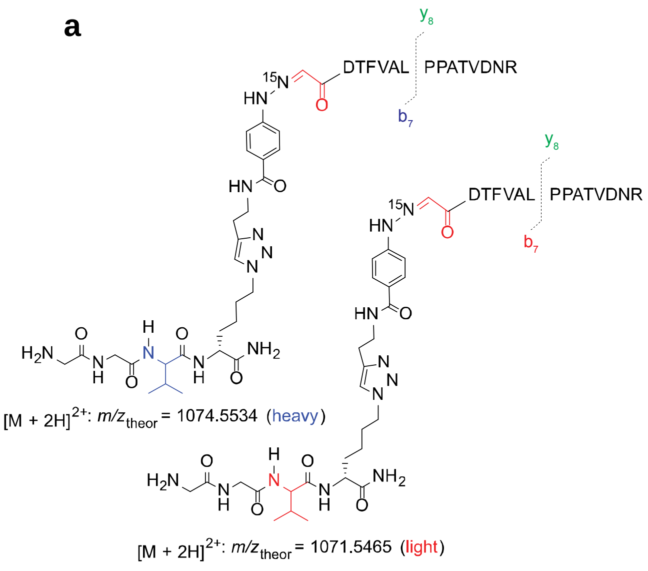

c

| | Theoretical *m/z* Heavy | Theoretical *m/z* Light | Measured *m/z* (error in Da) Heavy | Measured *m/z* (error in Da) Light |
|---|---|---|---|---|
| $b_2$ | 748.35 | 742.34 | 748.51 (0.15) | 742.26 (-0.08) |
| $b_3$ | 849.40 | 843.39 | 849.50 (0.10) | 843.26 (-0.12) |
| $b_4$ | 996.47 | 990.46 | 996.47 (0.00) | 990.47 (0.02) |
| $b_5$ | 1095.54 | 1089.52 | 1095.61 (0.07) | 1089.74 (0.21) |
| $b_6$ | 1166.58 | 1160.56 | 1166.64 (0.07) | 1160.19 (-0.37) |
| $b_7$ | 1279.66 | 1273.65 | 1279.71 (0.05) | 1273.67 (0.02) |
| $b_8$ | 1376.71 | 1370.70 | 1376.78 (0.07) | 1370.80 (0.10) |
| $b_9$ | 1473.76 | 1467.75 | 1473.57 (-0.19) | 1467.69 (-0.06) |
| $b_{10}$ | 1544.80 | 1538.79 | 1544.85 (0.05) | 1538.65 (-0.13) |
| $b_{11}$ | 1645.85 | 1639.84 | 1645.83 (-0.02) | 1639.93 (0.09) |
| $b_{12}$ | 1744.92 | 1738.90 | 1744.81 (-0.11) | - |
| $b_{13}$ | 1859.94 | 1853.93 | 1859.81 (-0.14) | - |
| $b_{14}$ | 1973.99 | 1967.97 | 1974.24 (0.25) | - |
| $b_7 - NH_3$ | 1262.63 | 1256.62 | 1262.73 (0.10) | 1256.65 (0.03) |
| $b_7 - NH_3CO$ | 1234.64 | 1228.62 | 1234.68 (0.04) | 1228.70 (0.08) |
| $b_7 - H_2O$ | 1261.65 | 1255.63 | 1261.69 (0.04) | 1255.74 (0.11) |
| $y_{14}$ | 1515.78 | | 1515.83 (0.05) | 1515.31 (-0.47) |
| $y_{13}$ | 1400.75 | | 1400.77 (0.02) | 1400.95 (0.2) |
| $y_{12}$ | 1299.71 | | 1299.72 (0.02) | 1299.88 (0.18) |
| $y_{11}$ | 1152.64 | | 1152.70 (0.06) | 1152.68 (0.04) |
| $y_{10}$ | 1053.57 | | 1053.60 (0.03) | 1053.68 (0.11) |
| $y_9$ | 982.53 | | 982.58 (0.05) | 982.61 (0.08) |
| $y_8$ | 869.45 | | 869.52 (0.08) | 869.49 (0.04) |
| $y_7$ | 772.39 | | 772.46 (0.07) | 772.46 (0.06) |
| $y_6$ | 675.34 | | 675.40 (0.06) | 675.46 (0.12) |
| $y_5$ | 604.30 | | 604.33 (0.02) | 604.39 (0.08) |
| $y_4$ | 503.26 | | 503.39 (0.14) | 503.26 (0.00) |
| $y_3$ | 404.19 | | 404.23 (0.04) | 404.21 (0.02) |
| $y_2$ | 289.16 | | 289.23 (0.07) | 289.21 (0.05) |
| $y_1$ | 175.12 | | 175.07 (-0.05) | 175.08 (-0.04) |

b

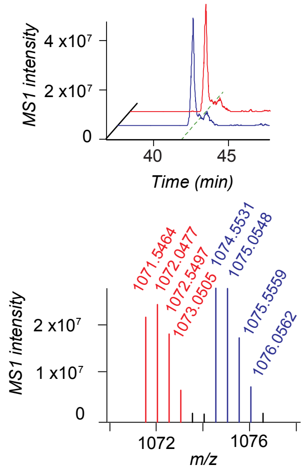

d

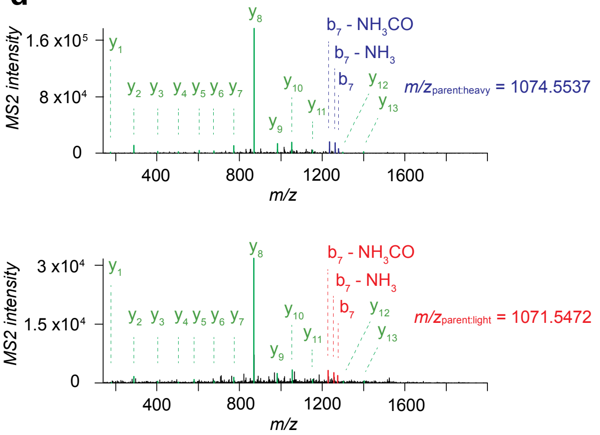

**Supplementary Fig. 17. MS characterization of the probe $^{15}$N-2 labelled glyoxylyl group of SCRN3**.

**a**, structures and theoretical parent masses of heavy- and light-tagged SCRN3 glyoxylyl peptides labelled by probe **$^{15}$N-2** and processed by the isoTOP-ABPP method. **b**, parent EICs (*upper*) and corresponding isotopic envelopes (*lower*) for heavy- (blue) and light- (red) tagged peptides detected from SCRN3-transfected HEK293T cells. **c**, summary table of theoretical versus observed spectra assignments generated under low-resolution MS2 conditions. **d**, MS2 spectra generated from parent ions of **b**. unshifted ions are shown in green and the shifted ion series in blue (heavy) and red (light).

**a**

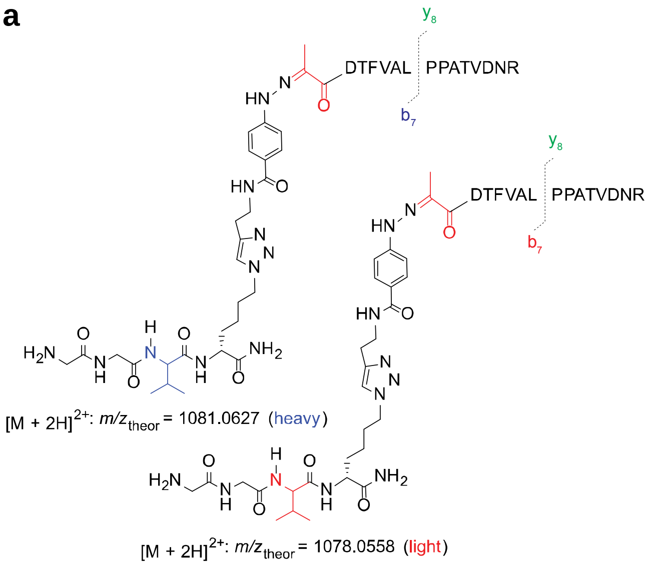

**b**

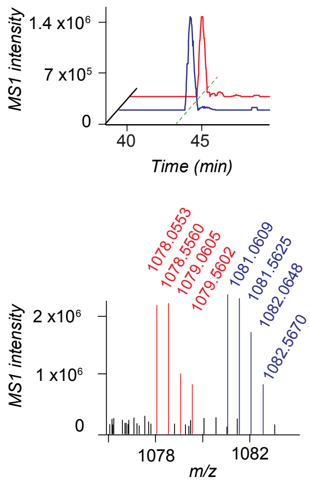

**c**

| | Theoretical *m/z* | | Measured *m/z* (error in Da) | |
|---|---|---|---|---|
| | Heavy | Light | Heavy | Light |
| $b_2$ | 761.37 | 755.36 | 761.70 (0.32) | 755.33 (-0.03) |
| $b_3$ | 862.42 | 856.41 | 862.53 (0.11) | 856.45 (0.04) |
| $b_4$ | 1009.49 | 1003.47 | 1009.16 (-0.33) | 1003.44 (-0.03) |
| $b_5$ | 1108.56 | 1102.54 | 1108.60 (0.04) | 1102.54 (-0.01) |
| $b_6$ | 1179.59 | 1173.58 | 1179.61 (0.02) | 1173.57 (-0.01) |
| $b_7$ | 1292.68 | 1286.66 | 1292.69 (0.01) | 1286.67 (0.01) |
| $b_8$ | 1389.73 | 1383.72 | 1389.96 (0.23) | 1383.72 (0.00) |
| $b_9$ | 1486.78 | 1480.77 | 1486.78 (-0.01) | 1480.90 (0.13) |
| $b_{10}$ | 1557.82 | 1551.81 | 1557.56 (-0.26) | 1551.95 (0.14) |
| $b_{11}$ | 1658.87 | 1652.85 | 1659.16 (0.29) | 1652.77 (-0.09) |
| $b_{12}$ | 1757.94 | 1751.92 | 1757.90 (-0.04) | 1751.70 (-0.22) |
| $b_{13}$ | 1872.96 | 1866.95 | - | 1867.25 (0.30) |
| $b_7 - NH_3$ | 1275.65 | 1269.64 | 1275.70 (0.05) | 1269.71 (0.08) |
| $b_7 - NH_3CO$ | 1247.66 | 1241.64 | 1247.62 (-0.04) | 1241.66 (0.02) |
| $b_7 - H_2O$ | 1274.67 | 1268.65 | 1274.75 (0.08) | 1268.75 (0.10) |
| $y_{14}$ | 1515.78 | | 1515.82 (0.04) | 1515.70 (-0.09) |
| $y_{13}$ | 1400.75 | | 1400.8 (0.05) | 1400.81 (0.06) |
| $y_{12}$ | 1299.71 | | 1299.84 (0.13) | 1299.60 (-0.11) |
| $y_{11}$ | 1152.64 | | 1152.65 (0.01) | 1152.67 (0.03) |
| $y_{10}$ | 1053.57 | | 1053.57 (0.00) | 1053.61 (0.04) |
| $y_9$ | 982.53 | | 982.57 (0.04) | 982.57 (0.04) |
| $y_8$ | 869.45 | | 869.45 (0.00) | 869.46 (0.01) |
| $y_7$ | 772.39 | | 772.42 (0.02) | 772.42 (0.02) |
| $y_6$ | 675.34 | | 675.46 (0.12) | 675.41 (0.07) |
| $y_5$ | 604.30 | | 604.37 (0.07) | 604.30 (0.00) |
| $y_4$ | 503.26 | | 503.28 (0.02) | 503.25 (0.00) |
| $y_3$ | 404.19 | | 404.31 (0.12) | 404.21 (0.02) |
| $y_2$ | 289.16 | | 289.22 (0.06) | 289.10 (-0.06) |
| $y_1$ | 175.12 | | 175.18 (0.06) | 175.18 (0.07) |

**d**

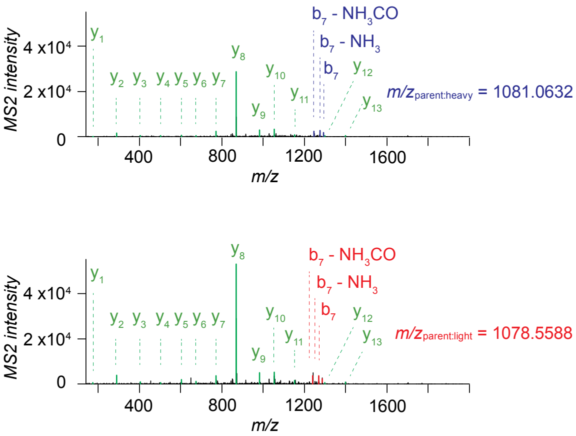

**Supplementary Fig. 18. MS characterization of the probe 2-labelled pyruvoyl group of SCRN3.**

**a**, structures and theoretical parent masses of heavy- and light-tagged SCRN3 pyruvoyl peptides labelled by probe **2** and processed by the isoTOP-ABPP method. **b**, parent EICs (*upper*) and corresponding isotopic envelopes (*lower*) for heavy- (blue) and light- (red) tagged peptides detected from SCRN3-transfected HEK293T cells. **c**, summary table of theoretical versus observed spectra assignments generated under low-resolution MS2 conditions. **d**, MS2 spectra generated from parent ions of **b**. unshifted ions are shown in green and the shifted ion series in blue (heavy) and red (light).

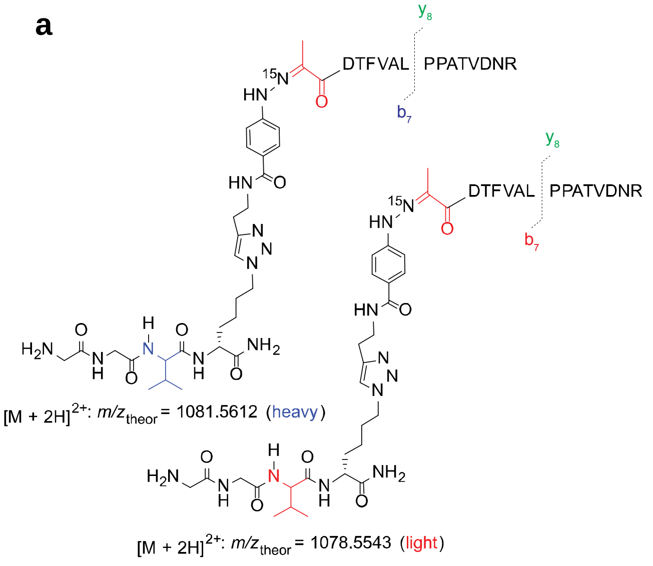

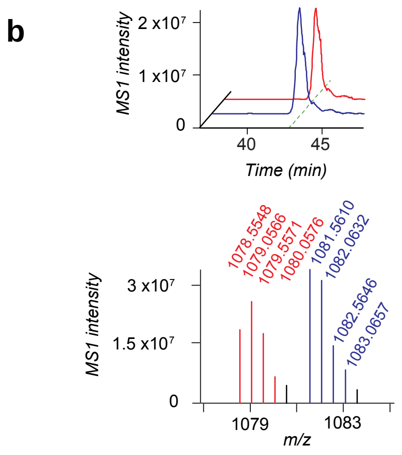

c

| | Theoretical *m/z* | | Measured *m/z* (error in Da) | |
|---|---|---|---|---|
| | Heavy | Light | Heavy | Light |
| $b_2$ | 762.37 | 756.36 | 762.41 (0.04) | 756.32 (-0.04) |
| $b_3$ | 863.42 | 857.40 | 863.79 (0.37) | 857.56 (0.16) |
| $b_4$ | 1010.49 | 1004.47 | 1010.54 (0.06) | 1004.46 (-0.01) |
| $b_5$ | 1109.55 | 1103.54 | 1109.46 (-0.09) | 1103.90 (0.36) |
| $b_6$ | 1180.59 | 1174.58 | 1180.30 (-0.29) | 1174.63 (0.05) |
| $b_7$ | 1293.67 | 1287.66 | 1293.82 (0.15) | 1287.66 (0.00) |
| $b_8$ | 1390.73 | 1384.71 | 1390.91 (0.18) | 1384.67 (-0.04) |
| $b_9$ | 1487.78 | 1481.77 | - | 1481.18 (-0.59) |
| $b_{10}$ | 1558.82 | 1552.80 | 1558.78 (-0.03) | 1552.87 (0.06) |
| $b_{12}$ | 1758.93 | 1752.92 | - | 1752.36 (-0.56) |
| $b_{14}$ | 1988.00 | 1981.99 | 1988.01 (0.00) | - |
| $b_7 - NH_3$ | 1276.65 | 1270.63 | 1276.69 (0.04) | 1270.68 (0.04) |
| $b_7 - NH_3CO$ | 1248.65 | 1242.64 | 1248.69 (0.04) | 1242.63 (-0.01) |
| $b_7 - H_2O$ | 1275.66 | 1269.65 | 1275.74 (0.08) | 1269.52 (-0.13) |
| $y_{14}$ | 1515.78 | | 1515.71 (-0.07) | 1515.93 (0.15) |
| $y_{13}$ | 1400.75 | | 1400.75 (0.00) | 1400.78 (0.03) |
| $y_{12}$ | 1299.71 | | 1299.72 (0.02) | 1299.81 (0.10) |
| $y_{11}$ | 1152.64 | | 1152.69 (0.05) | 1152.60 (-0.03) |
| $y_{10}$ | 1053.57 | | 1053.57 (0.00) | 1053.63 (0.06) |
| $y_9$ | 982.53 | | 982.54 (0.01) | 982.59 (0.06) |
| $y_8$ | 869.45 | | 869.45 (0.00) | 869.48 (0.04) |
| $y_7$ | 772.39 | | 772.39 (0.00) | 772.41 (0.02) |
| $y_6$ | 675.34 | | 675.48 (0.14) | 675.42 (0.08) |
| $y_5$ | 604.30 | | 604.27 (-0.03) | 604.36 (0.05) |
| $y_4$ | 503.26 | | 503.18 (-0.08) | 503.44 (0.18) |
| $y_3$ | 404.19 | | 404.19 (0.00) | 404.19 (0.00) |
| $y_2$ | 289.16 | | 289.17 (0.00) | 289.16 (0.00) |
| $y_1$ | 175.12 | | 175.01 (-0.11) | 175.08 (-0.04) |

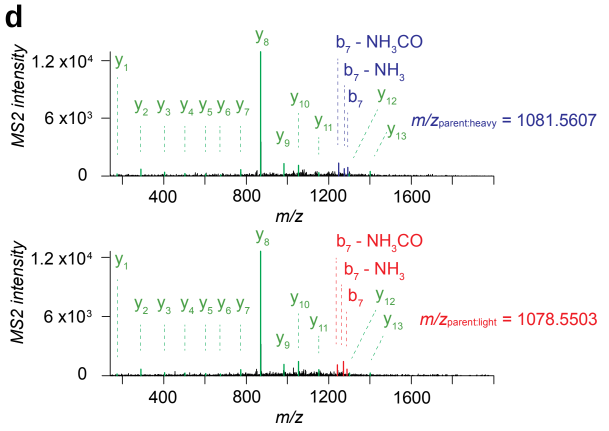

**Supplementary Fig. 19. MS characterization of the probe $^{15}$N-2 labelled pyruvoyl group of SCRN3**.

**a**, structures and theoretical parent masses of heavy- and light-tagged SCRN3 pyruvoyl peptides labelled by probe **$^{15}$N-2** and processed by the isoTOP-ABPP method. **b**, parent EICs (*upper*) and corresponding isotopic envelopes (*lower*) for heavy- (blue) and light- (red) tagged peptides detected from SCRN3-transfected HEK293T cells. **c**, summary table of theoretical versus observed spectra assignments generated under low-resolution MS2 conditions. **d**, MS2 spectra generated from parent ions of **b**. unshifted ions are shown in green and the shifted ion series in blue (heavy) and red (light).

**a**

*SCRN3-probe 2*

| *identity* | *$-NHNH_2$* | *$-NH^{15}NH_2$* | *shifted (Y/N)* |
|---|---|---|---|
| Glox6 | 2140.0811 | 2141.0808 | Y |
| Pyvl6 | 2154.0970 | 2155.0939 | Y |
| unknown | 2193.9991 | 2193.9919 | N |
| unknown | 1695.7558 | 1695.7558 | N |

**b**

*FTO-probe 2*

| *identity* | *$-NHNH_2$* | *$-NH^{15}NH_2$* | *shifted (Y/N)* |
|---|---|---|---|
| His231 | 2263.0747 | 2263.0771 | N |
| Tyr220 | 1611.7522 | 1611.7534 | N |
| unknown | 2298.0649 | 2298.0655 | N |

**c**

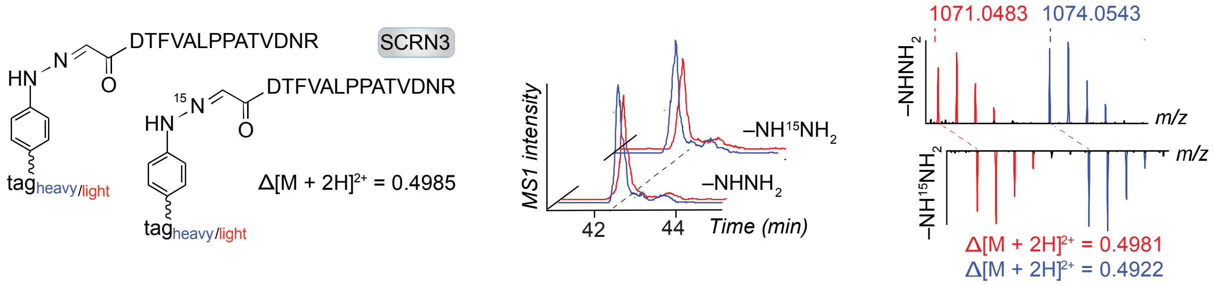

**Supplementary Fig. 20. Characterization of isotopic probes-labelled peptides in SCRN3 and FTO.**

**a**, sites of probe labelling identified from SCRN3-transfected cells treated with isotopic probes. **b**, sites of probe labelling identified from FTO-transfected cells treated with isotopic probes. **c**, peptide pairs and their isotopic envelopes from SCRN3-transfected cells treated with isotopic probes. Extracted ion chromatograms using monoisotopic *m/z* values of co-eluting heavy- and light-tagged peptides (blue and red, respectively) are shown on the *left*.

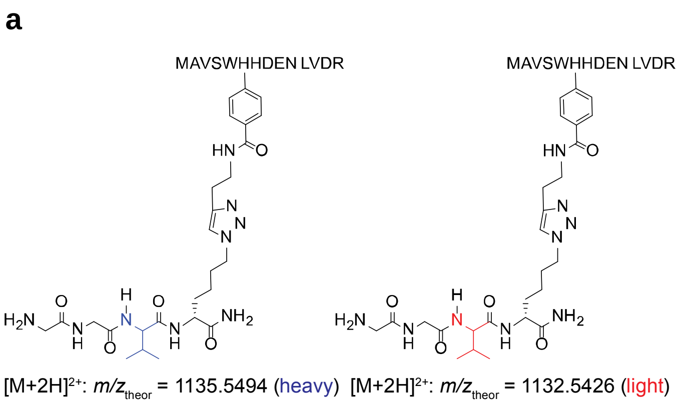

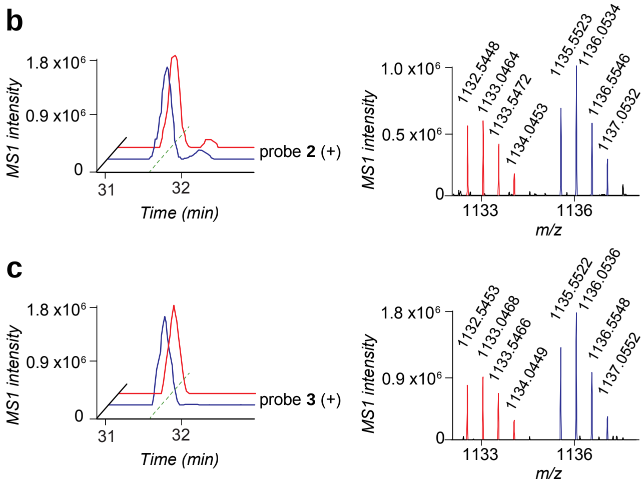

d

| | Theoretical *m/z* Heavy | Theoretical *m/z* Light | Measured *m/z* (error in ppm) Heavy | Measured *m/z* (error in ppm) Light |
|---|---|---|---|---|
| $b_6$ | 1273.6291 | 1267.6153 | 1273.6227 (-5.0) | - |
| $b_7$ | 1410.6880 | 1404.6742 | 1410.6604 (-19.6) | 1404.6783 (2.9) |
| $b_8$ | 1525.7150 | 1519.7012 | - | 1519.6971 (-2.7) |
| $b_{11}$ | 1881.8846 | 1875.8707 | 1881.8936 (4.8) | - |
| $y_9$ | 1695.8342 | 1689.8204 | 1695.8210 (-7.8) | 1689.8108 (-5.7) |
| $b_2$ | 203.0849 | | 203.0847 (-1.0) | 203.0848 (-0.5) |
| $b_4$ | 389.1853 | | 389.1866 (3.3) | - |
| y8 | 997.4697 | | 997.4695 (-0.2) | 997.4698 (0.1) |
| y7 | 860.4108 | | 860.4094 (-1.6) | 860.4117 (1.0) |
| y6 | 745.3839 | | 745.3836 (-0.4) | 745.3841 (0.3) |
| y5 | 616.3413 | | 616.3417 (0.6) | 616.3409 (-0.6) |
| y4 | 502.2984 | | 502.2997 (2.6) | 502.2976 (-1.6) |
| y3 | 389.2143 | | 389.2137 (-1.5) | 389.2131 (-3.1) |
| y2 | 290.1459 | | 290.1458 (-0.3) | 290.1458 (-0.3) |
| y1 | 175.1190 | | 175.1188 (-1.1) | 175.1187 (-1.7) |

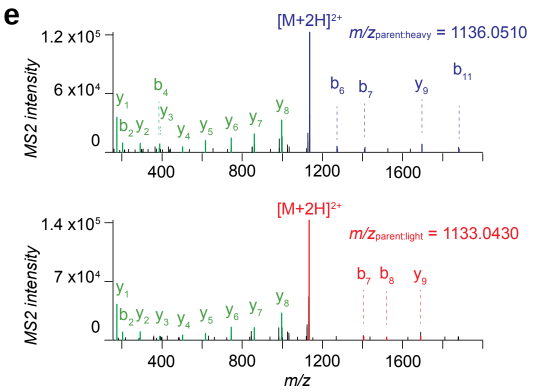

**Supplementary Fig. 21. MS characterization of the probe 2 and probe $^{15}$N-2 labelled peptides of FTO**.

**a**, structures and theoretical parent masses of heavy- and light-tagged FTO peptides labelled by probes **2** and **$^{15}$N-2** and processed by the isoTOP-ABPP method. **b**, parent EICs (*left*) and corresponding isotopic envelopes (*right*) for heavy- (blue) and light- (red) tagged peptides detected from FTO-transfected HEK293T cells treated with probe **2**. **c**, parent EICs (*left*) and corresponding isotopic envelopes (*right*) for heavy- (blue) and light- (red) tagged peptides detected from FTO-transfected HEK293T cells treated with probe **$^{15}$N-2**. **d**, summary table of theoretical versus observed spectra assignments generated under high-resolution MS2 conditions. **e**, MS2 spectra generated from parent ions of **b**. unshifted ions are shown in green and the shifted ion series in blue (heavy) and red (light).

**a**

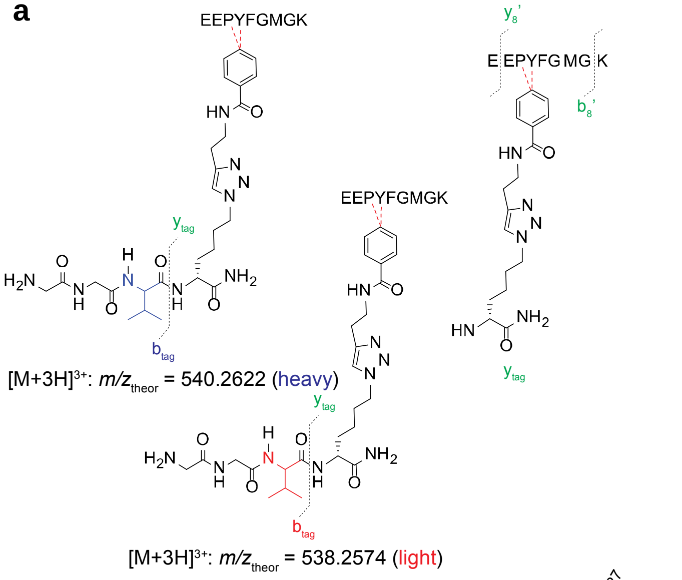

**c**

| | Theoretical $m/z$ | | Measured $m/z$ (error in ppm) | |
|---|---|---|---|---|
| | Heavy | Light | Heavy | Light |
| $b_4$ | 1080.5141 | 1074.5003 | 1080.5142 (0.1) | - |
| $b_5$ | 1227.5825 | 1221.5687 | 1227.5266 (-45.5) | - |
| $b_{tag}$ | 220.1324 | 214.1186 | 220.1325 (0.5) | 214.1186 (0.0) |
| $b_2$ | 259.0925 | | 259.0927 (0.8) | 259.0926 (0.4) |
| y5 | 539.2646 | | 539.2648 (0.4) | 539.2649 (0.6) |
| y4 | 392.1962 | | 392.1965 (0.8) | 392.1964 (0.5) |
| y3 | 335.1748 | | 335.175 (0.6) | 335.175 (0.6) |
| y2 | 204.1343 | | 204.1344 (0.5) | 204.1343 (0.0) |
| y1 | 147.1128 | | 147.1129 (0.7) | 147.1129 (0.7) |
| b8 | 1253.5408 | | 1253.5377 (-2.5) | 1253.5396 (-1.0) |
| y8'-NH3CO | 1225.5823 | | 1225.5353 (-38.3) | 1225.5369 (-37.0) |
| b8'-NH3CO | 1208.5193 | | 1208.51 (-7.7) | 1208.5082 (-9.2) |
| y7 | 1141.5612 | | 1141.5609 (-0.3) | 1141.5573 (-3.4) |
| y6 | 1044.5084 | | 1044.5032 (-5.0) | 1044.5121 (3.5) |

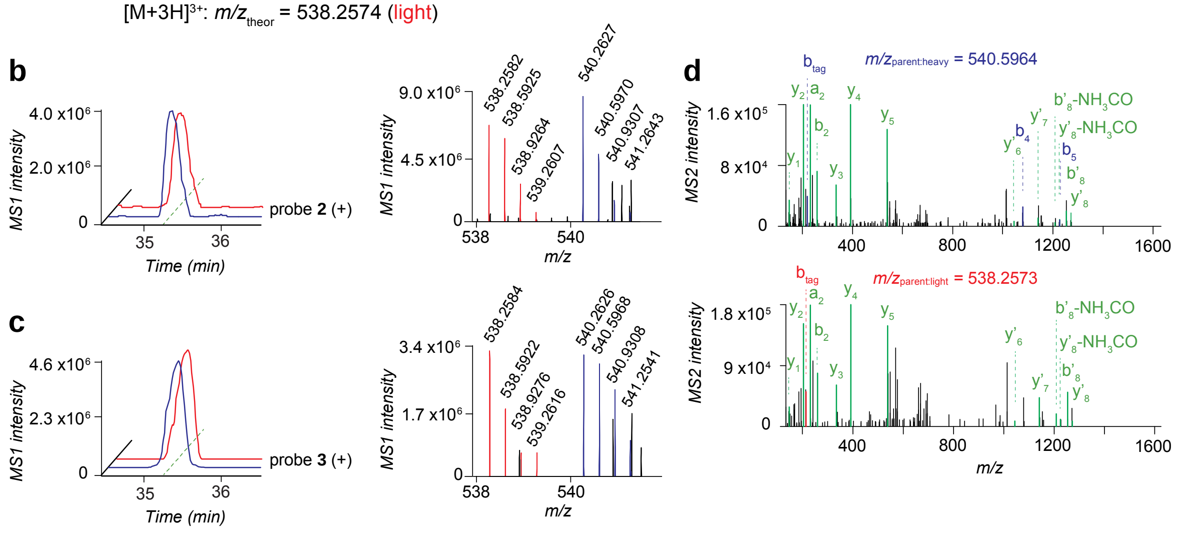

**Supplementary Fig. 22. MS characterization of the probe 2 and probe $^{15}$N-2 labelled peptides of FTO.**

**a**, structures and theoretical parent masses of heavy- and light-tagged FTO peptides labelled by probes **2** and **$^{15}$N-2** and processed by the isoTOP-ABPP method. **b**, parent EICs (*left*) and corresponding isotopic envelopes (*right*) for heavy- (blue) and light- (red) tagged peptides detected from FTO-transfected HEK293T cells treated with probe **2**. **c**, parent EICs (*left*) and corresponding isotopic envelopes (*right*) for heavy- (blue) and light- (red) tagged peptides detected from FTO-transfected HEK293T cells treated with probe **$^{15}$N-2**. **d**, summary table of theoretical versus observed spectra assignments generated under high-resolution MS2 conditions. **e**, MS2 spectra generated from parent ions of **b**. unshifted ions are shown in green and the shifted ion series in blue (heavy) and red (light).

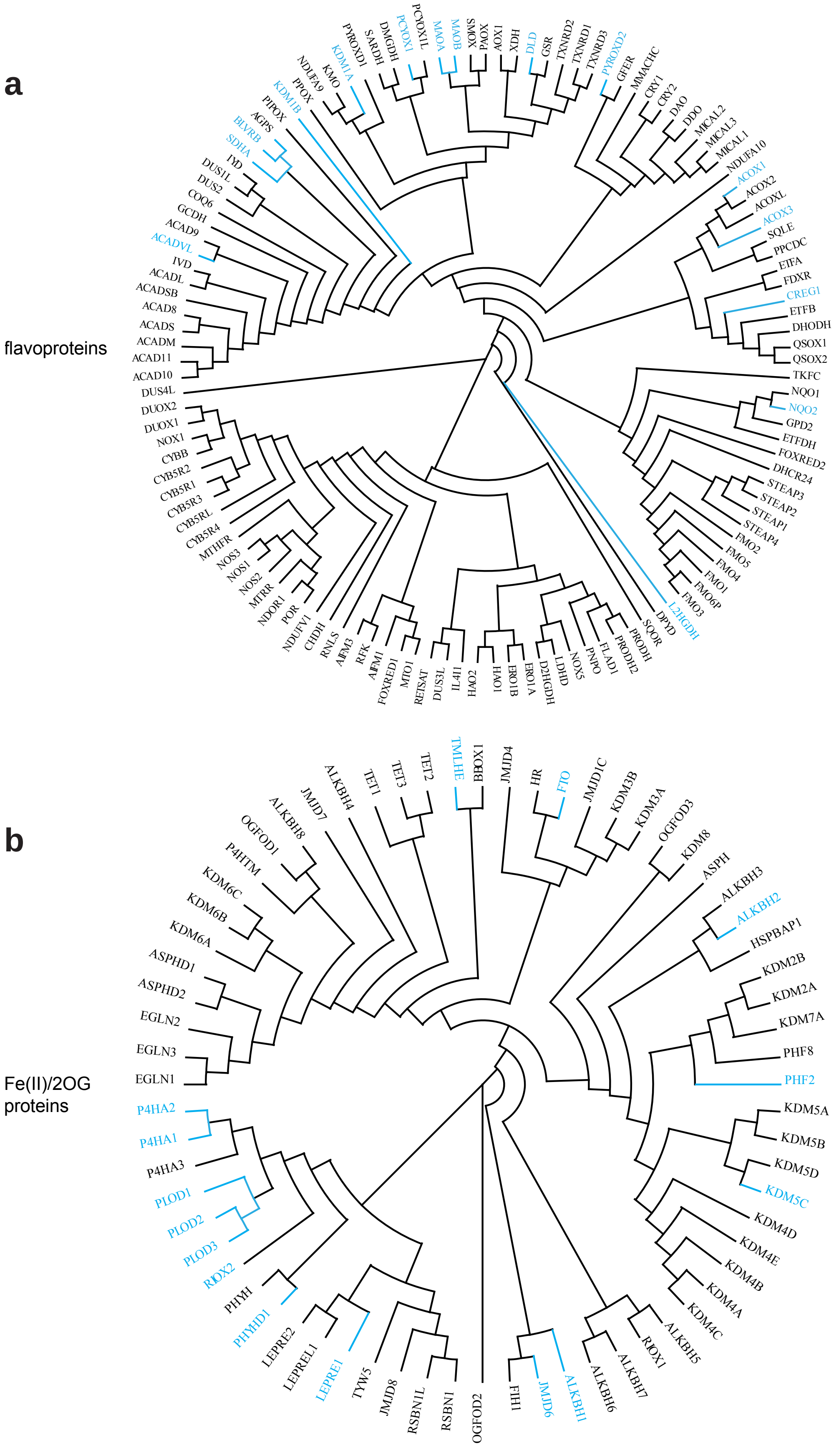
a
flavoproteins
DUS4L
ACAD10
ACAD11
ACADM
ACADS
ACAD8
ACADSB
ACADL
IVD
ACADVL
ACAD9
GCDH
COQ6
DUS2
DUS1L
IVD
SDHA
BLVRB
AGPS
PIPOX
KDM1B
PPOX
NDUFA9
KMO
KDM1A
PYROXD1
SARDH
DMGDH
PCYOX1
PCYOX1L
MAOA
MAOB
SMOX
PAOX
AOX1
XDH
DLD
GSR
TXNRD2
TXNRD1
TXNRD3
PYROXD2
GFER
MMACHC
CRY1
CRY2
DAO
DDO
MICAL2
MICAL3
MICAL1
NDUFA10
ACOX1
ACOX2
ACOXL
ACOX3
SQLE
PPCDC
ETFA
FDXR
CREG1
ETFB
DHODH
QSOX1
QSOX2
TKFC
NQO1
NQO2
GPD2
ETFDH
FOXRED2
DHCR24
STEAP3
STEAP2
STEAP1
STEAP4
FMO2
FMO5
FMO4
FMO1
FMO6P
FMO3
L2HGDH
DPYD
SQOR
PRODH
PRODH2
FLAD1
PNPO
NOX5
LDHD
D2HGDH
ERO1A
ERO1B
HAO1
HAO2
IL4I1
DUS3L
RETSAT
MTO1
FOXRED1
AIFM1
RFK
AIFM3
RNLS
CHDH
NDUFV1
POR
NDOR1
MTRR
NOS2
NOS1
NOS3
MTHFR
CYB5R4
CYB5RL
CYB5R3
CYB5R1
CYB5R2
CYBB
NOX1
DUOX1
DUOX2
b
Fe(II)/2OG proteins
EGLN1
EGLN3
EGLN2
ASPHD2
ASPHD1
KDM6A
KDM6B
KDM6C
P4HTM
OGFOD1
ALKBH8
JMJD7
ALKBH4
TET1
TET3
TET2
TMLHE
BBOX1
JMJD4
HR
FTO
JMJD1C
KDM3B
KDM3A
OGFOD3
KDM8
ASPH
ALKBH3
ALKBH2
HSPBAP1
KDM2B
KDM2A
KDM7A
PHF8
PHF2
KDM5A
KDM5B
KDM5D
KDM5C
KDM4D
KDM4E
KDM4B
KDM4A
KDM4C
ALKBH5
RIOX1
ALKBH7
ALKBH6
ALKBH1
JMJD6
FIH1
OGFOD2
RSBN1
RSBN1L
JMJD8
TYW5
LEPRE1
LEPREL1
LEPRE2
PHYHD1
PHYH
RIOX2
PLOD3
PLOD2
PLOD1
P4HA3
P4HA1
P4HA2

**Supplementary Fig. 23. Phylogenetic analysis of protein targets by hydrazine probes.**

**a**, phylogenetic tree of human flavoproteins. Proteins targeted by probes **1**, **2** or **3** with an average enrichment ratio ≥ 5 or competition ratio ≥ 3 are highlighted in phylogenetic tree (blue).
**b**, phylogenetic tree of human Fe(II)/2OG enzymes. Proteins targeted by probes **1**, **2** or **3** with an average enrichment ratio ≥ 5 or competition ratio ≥ 3 are highlighted in phylogenetic tree (blue).

### 3. Biological Methods

#### Materials

All materials were obtained from commercial suppliers and used without further purification. Phenylhydrazine probe (**2**) was synthesized following previous work[1]. Phenelzine probe (**3**), $^{15}N$-phenylhydrazine probe (**$^{15}$N-2**) and $^{15}N_2$-phenelzine probe (**$^{15}N_2$-3**) were synthesized according to the detailed experimental procedures outlined below. Phenelzine sulphate salt (**4**) was purchased from Millipore Sigma. GSK2879552 (**7**) was purchased from Selleckchem. Bizine (**8**) was purchased from APExBIO. D8 (**9**) and D31 (**10**), used as KDM1A inhibitors[2], were gifts from Professor Bin Yu in School of Pharmaceutical Sciences at Zhengzhou University. 1,4-DPCA (**5**), used as an iron chelator[3], was a gift from Professor Ellen Heber-Katz in the Laboratory of Regenerative Medicine at Lankenau Institute for Medical Research. Meclofenamate (**6**) was purchased from MedChemExpress.

#### Stock solutions of probes and competitor

Solutions (100 μM) of probe **2** and probe **$^{15}$N-2** were prepared in 10% dimethyl sulfoxide (DMSO) according to previous procedure[1]. Working stock solutions (100–400 μM) of probe **3**, probe **$^{15}N_2$-3** and phenelzine **4** were prepared in $H_2O$ from the trifluoroacetate, trifluoroacetate and sulphate salts, respectively. The stocks were neutralized to pH 6.0-7.0 by 5.0 M sodium hydroxide solution using fisherbrand plastic pH indicator strips (pH range: 0.0-14.0). Solutions were stored at -80 °C prior to use.

#### Cloning and mutagenesis

Transfection grade plasmids encoding human proteins in mammalian expression vectors were purchased from manufacturers or prepared according to a previous protocol[1]. Plasmids containing full length human genes of lysine-specific histone demethylase 1A (KDM1A) from clone ID OHu27145, ribosyldihydronicotinamide dehydrogenase [quinone] (NQO2) and its mutant Y156A from clone ID OHu23491D, secernin-1 (SCRN1) from clone ID OHu13636D and alpha-ketoglutarate-dependent dioxygenase FTO and its mutants (H231A, D233A, H307A and R316A) from clone ID OHu24899D in mammalian vector pcDNA3.1+/C-(K)-DYK were purchased from GenScript. Plasmid containing full length gene of secernin-2 (SCRN2) in bacterial pET-30b(+) vector was purchased from GenScript. Bacteria stock transformed by plasmids containing full length genes of amine oxidase [flavin-containing] A (MAOA) from clone ID 2990003 and amine oxidase [flavin-containing] B (MAOB) from clone ID 4800208 in pOTB7 and pBluescriptR vector, respectively, were purchased from Dharmacon Inc. Plasmids containing full length genes of secernin-3 (SCRN3) and its mutant C6A and FTO in mammalian expression pRK5 vector (code 3944 in Addgene database) were amplified from DNA templates in previous study[1].

Bacteria containing human MAOA (MAOB) genes were seeded on LB agar plates with chloramphenicol (or ampicillin) resistance. After overnight incubation at 37 °C, a single colony from plate was picked and grown in LB medium with resistance for 16 h at 37 °C. Plasmids were extracted from the bacteria using Zyppy Plasmid Miniprep Kit (Zymo) according to the manufacturer's instructions. The plasmids were used as template DNA for amplification (C1000 Touch Thermal Cycler, Bio-Rad) to introduce NotI and SalI restriction sites using Phusion polymerase (New England BioLabs Inc.) according to the manufacturer's protocol. Primers for MAOA plasmid are 5'-TTTTGTCTACGCCACCATGGAGAATCAAGAGAAGGCGAGTATC-3' (forward) and 5'- AAAAGCGGCCGCAGACCGTGGCAGGAGC-3' (reverse). Primers for MAOB are 5'-TTTTGTCTACGCCACCATGAGCAACAAATGCGACGTGG-3' (forward) and 5'-AAAAGGGGGCTACTTGTGAGAGTCGCGGCCGCTTTT-3' (reverse). MAOA (or MAOB) gene and the pRK5 empty vector were then digested by restriction enzymes NotI and SalI (New England BioLabs Inc.), the resulted linear gene and vector were purified by Gel Extraction Kit (QIAGEN), ligated by T4 DNA ligase (NEB). The ligated plasmids were transformed into chemically competent MAX Efficiency Escherichia coli (E. coli) strain DH5α cells (invitrogen) following heat shock procedure or ElectroMAX DH5α-E Competent Cells (invitrogen) using MicroPulser Electroporator (Bio-Rad) following manufacturer's protocol. The cells were then grown in LB medium with antibiotic resistance for plasmid extraction by Zyppy.

Plasmid containing human SCRN2 gene was used as DNA template for amplification to introduce SalI and NotI restriction sites using Phusion polymerase (NEB). Primers with the following sequences were used for amplification: 5'-TTTTGTCGACGCCACCATGGCGAGCAGCAGCCCGG-3' (forward) and 5'-AAAAGCGGCCGCCGCATACGCTTGGCTCTCACGCTT-3' (reverse). After digestion by restriction enzymes, the isolated gene was ligated into pRK5 empty vector. The resulting plasmids were transformed into chemically competent MAX Efficiency DH5α cells using heat shock. The cells were then grown in LB medium with antibiotic resistance for plasmid purification and sequencing.

Plasmids containing human NQO2 gene was used as DNA template for amplification to introduce SalI and HindIII restriction sites using Phusion polymerase (NEB). Primers with the following sequences were used for amplification: 5'-

AAAAGTCGACATGGCAGGTAAGAAAGTACTCATTG-3' (forward) and 5'-TTTTAAGCTTTTGCCCGAAGTGCCAGT-3' (reverse). After digestion by restriction enzymes, the isolated gene was ligated into empty pET-45b(+) vector with *N*-terminal HisTag, the resulted plasmids were transformed into chemically competent MAX Efficiency DH5α cells using heat shock. The cells were then grown in LB medium with antibiotic resistance for plasmid purification and sequencing.

Plasmids containing genes encoding wild type proteins (MAOA, MAOB, KDM1A, SCRN2 and SCRN1) were used as DNA templates for mutagenesis. Mutagenesis primers were designed by QuikChange Primer Design (Agilent), and purchased from Integrated DNA Technologies, Inc. (IDT). Mutants (MAOA-C406A, MAOB-C397A, KDM1A-K661A, SCRN2-C12A, and SCRN1-S7C) were generated by QuikChange site-directed mutagenesis (Agilent) using Pfu Ultra High-Fidelity DNA polymerase (Agilent) or Phusion polymerase (NEB) according to manufacturer's protocols.

The concentrations of plasmids were all determined by Nanodrop One (Thermo Scientific). All primers were purchased from Integrated DNA Technologies, Inc. (IDT). All plasmids were sent to Sanger sequencing facility (Penn Genomics Analysis Core) for confirmation prior to use. The sequencing primers for genes in pRK5 vector are: 5'-ATTTAGGTGACACTATAGAA-3' (sp6, forward) and 5'-TGTAACCATTATAAGCTG-3' (reverse, obtained from IDT). The sequencing primers for genes in pcDNA3.1+/C-(K)-DYK vector are: 5'-CGCAAATGGGCGGTAGGCGTG-3' (CMV forward) and 5'-TAGAAGGCACAGTCGAGG-3' (BGH reverse). The sequencing primers for genes in pET-45b(+) vector are: 5'-TAATACGACTCACTATAGGG-3' (T7 promoter/primer, forward) and 5'-GCTAGTTATTGCTCAGCGG-3' (T7 terminator, reverse). The proteins expressed via transient transfection in HEK293T cells contain an additional 15 amino acids appended to their *C*-terminus (A3G4DYKD4K) when pRK5 vector was applicable, or 8 amino acids appended to their *C*-terminus (DYKD4K) when pcDNA3.1+/C-(K)-DYK was applicable. The DYKD4K appendage in each protein contains the classic FLAG epitope that permits detection by western blotting or purification by affinity chromatography. Plasmids were purified by QIAGEN Plasmid Maxi Kit when large amounts were required.

**Expression and purification of NQO2 protein**

The expression and purification of NQO2 were adapted from previous studies[4,5]. Bacterial expression of NQO2 protein was confirmed by small scale growth (10 mL) after transformation into chemically competent BL21 (DE3*) cells (Invitrogen) using heat shock. Large cultures (1 L/flask) were grown at 37 °C in rich LB broth [35 g/L tryptone, 20 g/L yeast extract, 5 g/L NaCl,

0.5% glycerol, and 0.1 g/L carbenicillin, pH 7.3] to an optical density of 0.7-0.9 at 600 nm (~8 h). The flasks were cooled rapidly on ice prior to induction with isopropyl β-d-1-thiogalactopyranoside (IPTG) at a final concentration of 1 mM. Cultures were grown at 16 °C for an additional 16-20 h and cells were harvested by centrifugation (5,000 g, 30 min, 4 °C). The resulted cell paste (yield: ~8-10 g/L growth) was flash frozen in liquid nitrogen and stored at -80 °C. The paste (~30 g) was resuspended (5 mL/g) in 50 mM Na-HEPES buffer (pH 7.5) containing 300 mM NaCl, 5 mM imidazole, 0.1 mg/mL DNase I, 1 mg/mL lysozyme, 1 mM $MgCl_2$ and 1 mM $CaCl_2$. The cells were then lysed by two passages through a microfluidizer at 18,000 psi and the cell debris was pelleted by centrifugation (30,000 g, 20 min, 4 °C). The supernatant was slowly stirred (50-100 rpm) for ~1 h with Nickle-nitrilotriacetic acid (Ni-NTA) resin (QIAGEN) (ratio: 0.5 mL resin/g paste) at 4 °C for batch purification. The slurry was loaded onto the column (2.5 x 20 cm) and washed with 50 mM Na-HEPES buffer (pH 7.5) containing 300 mM NaCl and 5 mM imidazole until absorption of the eluate at 280 nm (A280) and 260 nm (A260) were both ~0 (~5 column volumes). Protein was eluted from the resin with 50 mM Na-HEPES buffer (pH 7.5) containing 100 mM NaCl and 200 mM imidazole (~8 column volumes). Fractions containing protein (yellow color) were combined and dialyzed against 50 mM Na-HEPES buffer (pH 7.5) containing 150 mM NaCl for 12 h at 4 °C, followed by 2 × 2 h dialysis at 4 °C. Following dialysis, the protein was concentrated to ~8 mg/mL prior to being flash-frozen and stored at -80 °C with the presence of 10% glycerol. The yield of NQO2 was ~3.5 mg/g paste. Protein concentration was determined on a Nanodrop One by assuming a molar absorptivity (ε280) of 44,920 $M^{-1}cm^{-1}$ (estimated using ProtParam in Expasy). The flavin content of recombinant NQO2 was confirmed by yellow color, LC-MS analysis, and the characteristic broad UV peak at 450 nm measured on Nanodrop One.

**Liquid chromatography-mass spectrometry (LC-MS) analysis of FAD adducts**

Detection of FAD adducts was adapted from previous studies[4-6]. NQO2 protein stock was diluted to 300 μM by PBS. An aliquot of 250 μL in the presence of 6 mM NRH was added PBS (control), phenelzine **4** or probe **3** with a final concentration of 3 mM (10-molar fold excess). The mixture was incubated for 30 min at 37 °C. The reaction was then quenched by incubation at 95 °C for 7 min to inactivate NQO2 and release the FAD adducts. After centrifugation (17,000 g, 5 min, 4 °C), the denatured protein was precipitated, and the supernatant was transferred to an LC vial for LC-MS analysis.

An Acquity UPLC system coupled to an Acquity SQ detector (Waters) was used for LC-MS analysis. An Acquity C18 column (2.1 × 50 mm, 1.8 μm, Waters) maintained at 30 °C was used with a mobile phase consisting of a linear gradient of A (0.1% formic acid in water) and B (0.1% formic acid in MeCN) under the following conditions: 0 → 0.5 → 2.5→ 3 min, 5% → 5% → 95% → 95% B. The flow rate was 0.3 mL/min, the injection volume was 2 μL, and the UV detector was set at 254 nm to display chromatograms. The MS scan range was 150–2,000 Da in both positive and negative modes. All data acquisition and analyses were controlled by Waters MassLynx Version 4.1 software.

**Cell culture**

Low-passage HEK293T (human embryonic kidney cells) and MDA-MB-231 (triple-negative breast cancer cells) cells (ATCC) were cultured in Dulbecco's Modified Eagle's Medium (DMEM) (with high glucose, *L*-glutamine and sodium pyruvate) supplemented with 10% (*v*/*v*) fetal bovine serum (FBS) (Corning), penicillin (100 U/mL)/streptomycin (100 μg/mL) at 37 °C in a humidified atmosphere containing 5% $CO_2$. For Stable Isotope Labelling by Amino acids in Cell culture (SILAC) experiments[7,8], each cell line was passaged a minimum of six times in DMEM for SILAC (Thermo Fisher Scientific, A33822) containing dialyzed FBS (Silantes, 281001200), penicillin (100 U/mL)/streptomycin (100 μg/mL), and isotopically enriched *L*-[$^{13}C_6{}^{15}N_2$]lysine hydrochloride and *L*-[$^{13}C_6{}^{15}N_4$]arginine hydrochloride (100 μg/mL each, 550 μM and 475 μM, respectively, Sigma) as "heavy" medium [or naturally abundant isotopologues lysine hydrochloride and arginine hydrochloride (100 μg/mL each, 550 μM and 475 μM, respectively, Sigma) as "light" medium].

**Transfection of HEK293T cells**

HEK293T cells were grown to ~40% confluence under standard growth conditions before transfection. A standard transfection ratio of vector/transfection reagent (3:1, *w*/*w*) was used. 4 μg plasmid in appropriate expression vector and 12 μg of neutralized (pH 7) polyethyleneimine (PEI, Polysciences Inc.) 'MAX' (MW 40,000, 1 mg/mL) were premixed in 200 μL FBS-free DMEM medium, the resulted mixture was incubated at room temperature for 30 min before added to a 6-cm dish. 10 μg plasmid and 30 μg PEI in 500 μL FBS-free DMEM was used for a 10-cm dish. A 6-cm (or 10-cm) dish received 4 μg (or 10 μg) of the appropriate empty vector was used as control group ('mock') of transfection. Cells were incubated for ~48 h before labelling *in situ*.

***In situ* labelling of cells with hydrazine probes**

For gel-based experiments, cells were seeded and incubated until ~100% confluence in 6-cm (or 10-cm) dish containing 3 mL (or 10 mL) culture medium. Probe (or competitor) treatment solutions at 1 mM were prepared in ice-cold serum-free DMEM (~10% culture volume) supplemented with 10 mM Na-HEPES (pH 7.5). Cells were then transferred on ice, washed with culture volume ice-cold phosphate buffered saline (PBS, pH 7.4), replenished with treatment solution and incubated for 30 min at 37 °C. For time-dependent labelling experiments, cells were incubated with probe for 0.5, 15 and 30 min, respectively. For dose-dependent labelling experiments, cells were treated with 0.25, 1, 2, and 5 mM probe, respectively. For competition labelling experiments, cells were pretreated with non-clickable competitor at desired concentration for 15 min, then added probe stock solution to a final concentration of 1 mM, and subsequently incubated for another 15 min. To generate the inhibitory curves of inhibitors, cells were pretreated with inhibitor for 30 min, then added probe stock solution, and subsequently incubated for another 30 min.

For mass spectrometry-based experiments, cell labelling was carried out in a similar manner maintaining the same concentrations using isotopically "light" and "heavy" SILAC cells that were also grown to ~100% confluence prior to treatment. Isotopically heavy cells were treated with 1 mM probe **3**, while isotopically light cells were treated with the non-clickable analog phenelzine **4** at 1 mM in "enrichment" experiments (or treated with phenelzine at 10 mM for 15 min, then added probe stock solution to a final concentration of 1 mM and incubated for another 15 min in "competition" experiments).

Following treatments, cells were placed on ice followed by aspiration of treatment solutions, washed with culture volume ice-cold PBS to remove residual probe (and/or competitor), harvested by scraping, collected by centrifugation (1,400 g, 2 min, 4 °C), washed again by resuspension in ice-cold PBS and centrifugation, and frozen as pellets at -80 °C prior to use.

**Proteome preparation of cells for gel- and MS-based experiments**

Cell pellets were resuspended on ice in ice-cold PBS (200-400 μL) and lysed by a Branson SFX250 Sonifier equipped with a 102C microtip (3 × 10 pulses, 0.3 second on and 2 second off, 15% energy). The resuspension volume and sonication parameters were adjusted according to the cell pellet yield. Soluble (cytosolic) and membrane proteomes were separated by ultracentrifugation (100,000 g, 30 min, 4 °C) in a thermo fisher S55-A2 rotor. Soluble proteome was transferred to new tubes, and membrane proteome was resuspended by adding the same volume of PBS followed by probe sonication. Protein concentrations of each fraction were determined by DC protein assay kit from Bio-Rad on a microplate reader (Biotek ELx808 plate reader). For SILAC experiments, isotopically heavy and light cell lysates were mixed 1:1 by protein weight.

**Gel-based analysis of probe-labelled proteins**

Soluble (or membrane) proteomes from treated cells were diluted to 1 and 2 mg/mL, respectively. To each sample (50 μL), 6 μL of a freshly prepared copper(I)-catalyzed azide alkyne cycloaddition (CuAAC or "click") reagent mixture containing 3 μL of 1.7 mM tris(benzyltriazolylmethyl)amine (TBTA) in DMSO:*t*-BuOH (1:4), 1 μL of 50 mM $CuSO_4$ in $H_2O$, 1 μL of 1.25 mM rhodamine-azide in DMSO, and 1 μL of freshly prepared 50 mM tris(2-carboxyethyl)phosphine (TCEP) in $H_2O$ was added to conjugate the fluorophore to probe-labelled proteins. TBTA was purchased from TCI and rhodamine-azide (TAMRA azide, 5-isomer) was purchased from Lumiprobe. Upon addition of the click mixture, each sample was immediately vortexed and then allowed to react at room temperature under rotation for 1 h before quenched by addition of 17 μL of 4X sodium dodecyl sulfate (SDS) loading buffer. Samples (15 μL/well for 15-well gel, or 40 μL/well for 10-well gel) were resolved by SDS-PAGE (10% or 4-20% acrylamide mini gel from invitrogen) and visualized by in-gel fluorescence scanning on a ChemiDoc MP Imaging System (Bio-Rad).

**Western blotting**

After scanning fluorescence, the proteins in gel were transferred to polyvinylidene difluoride (PVDF, MilliporeSigma) in Towbin buffer using a Trans-Blot® SD Semi-Dry Transfer Cell (Bio-Rad). The membrane was blocked for ~1 h at room temperature with 5% nonfat dry milk (w/v) in Tris-buffered saline with 0.05% Tween 20 (TBST) and incubated with primary antibody in the same solution overnight at 4 ºC or 1 h at room temperature. The antibodies included anti-FLAG (1:2500, F1804, Sigma) and anti-His6 (1:1000, ab18184, Abcam). The blots were washed (3 × 2 min, TBST), incubated with secondary goat anti-mouse antibody (1:10,000, ab150113, abcam) in milk for 1 h at room temperature, washed again (3 × 2 min, TBST), and visualized on a ChemiDoc MP Imaging System (Bio-Rad) using a Alexa Fluor 488 channel.

**Sample preparation of SILAC experiments**

Profiling experiments were adapted as previously reported[1]. The isotopically heavy and light (1:1) lysate (soluble or membrane) mixture (1 mg total protein) was diluted to 1 mL in PBS. Click reactions were scaled accordingly (except for biotin-azide) to maintain final concentrations of 100 μM TBTA, 1 mM $CuSO_4$, 100 μM biotin-azide (10 mM in DMSO) and 1 mM TCEP. The mixture was vortexed and placed on a rotator at room temperature for 1 h. Sequential addition of pre-chilled methanol (MeOH, 2 mL), chloroform ($CHCl_3$, 0.5 mL) and PBS (1 mL) on ice quenched the reaction. The precipitated proteome was centrifuged (5,000 g, 10 min, 4 °C) to fractionate the protein interphase from the organic and aqueous solvent layers. The protein pellet was washed with cold 1:1 MeOH:$CHCl_3$ (3 × 1 mL), mildly sonicated in cold 4:1 MeOH:$CHCl_3$ (2.5 mL) and pelleted once more by centrifugation (5,000 g, 10 min, 4 °C) to ensure click reagents were efficiently removed. The protein pellet was resuspended by mild sonication in a freshly prepared solution of proteomics-grade urea (500 µL, 6 M in PBS). Disulfides were reduced with TCEP (final concentration: 9 mM) pre-neutralized with potassium carbonate (final concentration: 27 mM) for 30 min at 37 °C. Reduced thiols were then alkylated by iodoacetamide (final concentration: 45 mM) for 30 min at room temperature protected from light. SDS [2% (w/v)] was added to ensure complete denaturation. The solution was diluted to 0.2% SDS in PBS (final volume: 5 mL) and incubated with pre-equilibrated streptavidin agarose resin (50 µL column volume, 100 µL 1:1 slurry, Pierce) for ~1.5–2 h at room temperature on a rotator. The streptavidin beads were collected by centrifugation (1,400 g, 1–2 min) and sequentially washed with 0.2% SDS in PBS (3 × 5 mL), PBS (3 × 5 mL) and $H_2O$ (3 × 5 mL) to remove unbound protein, excess detergent, and small molecules. The resin was transferred to a Protein LowBind tube (Eppendorf) and bound proteins were digested on-bead overnight at 37 °C in ~200 µL PBS containing 2 µg sequencing grade porcine trypsin (Promega), 2 M urea and 1 mM $CaCl_2$. The proteolyzed supernatant was transferred to a fresh Protein LowBind tube, acidified with formic acid (5%) to inactivate trypsin, stored at –80 °C or desalted immediately.

**IsoTOP-ABPP sample preparation to isolate probe-captured peptides**

To enrich and discover the probe-labelled peptide(s) of a protein in cells, the previously described isoTOP-ABPP protocol was adapted[1,9,10]. HEK293T cells were transfected by plasmid containing gene of interest, treated with probe, and prepared as described above to obtain soluble proteome (2 mg total protein in 1 mL PBS). Half of the proteome (0.5 mL) was conjugated to the light TEV tag and the other half to the heavy TEV tag (structures shown in **Biological Methods**). Click reactions were scaled accordingly to maintain final concentrations of 100 µM TBTA, 1 mM $CuSO_4$, 100 µM of light or heavy biotin-TEV-azide (5 mM in DMSO), and 1 mM TCEP. The mixture was vortexed and placed on a rotator at room temperature for 1 h. The light- and heavy-labelled samples were combined and centrifuged (16,000 g, 5 min, 4 ºC) to yield a protein pellet that was mildly sonicated in and washed by ice-cold methanol (2 × 0.5 mL), then solubilized with 1.2% SDS (1 mL in PBS) by mild sonication. The sample was diluted to ~0.2% SDS with PBS (~6 mL) and incubated with pre-equilibrated streptavidin agarose resin (100 µL 1:1 slurry) for ~3 h at room temperature. The resin was washed as described above for SILAC experiments, transferred to clean microcentrifuge tubes, and resuspended in urea (500 μL, 6 M in PBS). Cysteines were reduced and alkylated with TCEP and iodoacetamide, respectively as described above. The resin was washed once with 1 mL PBS to remove the reagents and bound proteins were digested with trypsin (2 μg) for overnight (8–12 h) at 37 ºC in the presence of 2 M urea (200 μL, in PBS) and $CaCl_2$ (1 mM).

Unmodified peptides, urea, and trypsin were removed by sequential washes with PBS (9 × 0.5 mL). The resin was transferred to clean microcentrifuge tubes and equilibrated with TEV buffer (50 mM Tris, pH 8). Remaining immobilized peptides were released with TEV protease (~2.2 μM in ~335 μL TEV buffer at 30 ºC for 6 h). TEV proteolytic peptides containing heavy- and light-TEV tags were transferred to new tube and recovered from the resin with $H_2O$ (2 × 50 μL). Sample was desalted immediately without addition of formic acid or stored at –80 ºC prior to desalting.

**Peptide desalting**

Peptide samples were desalted prior to analysis by using in-house packed stage-tips. Stage-tips were manufactured by sealing five disks of C18 material (Empore, 3M Company) at the bottom of a P200 tip. Stage-tips were equilibrated with 50 μL of methanol, 50 μL of 80% acetonitrile in $H_2O$ containing 0.1% trifluoroacetic acid (TFA), and 50 μL of water containing 0.1% TFA by centrifugation (1,000 g, ~1-2 min). The sample was loaded to the stage-tip and centrifuged to flow through, washed with 75 μL of water containing 0.1% TFA. The stage-tip was transferred to a new collection tube, eluted by 75 μL of 80% acetonitrile in $H_2O$ containing 0.1% TFA. The sample was dried by SpeedVac (SAVANT SVC100H Refrigerated Condensation Trap) under vacuum for 30-60 min at room temperature, stored at –80 °C. Prior to analysis, sample was dissolved in 10 μL of 2% acetonitrile in $H_2O$ containing 0.1% TFA, transferred to LC vial. Unless otherwise noted, an aliquot (5 μL) was injected into LC-MS/MS system.

**Nano-column preparation**

Nano-columns for chromatographic separation of peptides were packed in-house. A 30 cm long fused-silica capillary column (inner diameter 75 μm and outer diameter 375 μm) was cut off from tubing roll (cat. log. No.: 1068150019, molex) by ceramic scoring wafer (RESTEK). Centered at 4.5 cm from the left end, coating material with a length of 1-1.5 cm was burned off completely using lab flame (Bunsen or alcohol burner). The left end of the capillary was immersed into frit mixture (Kasil 1624:formamide = 15:5 μL, ratio 3:1) until the solution covered the bare glass area by capillary reaction, and then the same end was dipped into $H_2O$ to push the frit mixture further (~0.2-0.3 cm) inside the capillary. The capillary was heated for 1-3 seconds at the spot

~5.0 cm from the left end by a soldering tip (cat. log. No: SSA51, SEALODY) prewarmed to 200 °C until the frit solidified to a length of ~0.3 cm. It is optional to check the color of the solid frit (brown to pale yellow from center to both ends) under microscope (cat. log. No.: 1810430, AmScope). The *right* end of the capillary was then immersed into a $H_2O$-containing LC vial inside a stainless-steel chamber (manufactured by Physical Sciences Machine Shop, University of Pennsylvania) which was then sealed by screws. After the capillary was fixed by ferrule (1/8" Ferrule 0.4 mm ID PTFE, Chromatography Research Supplies) tightened with screw, the chamber was then pressured by helium (20-200 psi) to push $H_2O$ through the frit. After ~10 drops of $H_2O$ coming off from the capillary (~5-10 min), the pressure inside chamber was released, the $H_2O$-containing LC vial was replaced with a pre-sonicated LC vial containing ~2-5 mg of C18 material (3 µm, ReproSil-Pur 120 C18-AQ), 1.5 mL of $CH_2Cl_2$:MeOH (2-3:1, v/v), and a magnetic stir bar. The chamber was sealed, placed on a stir plate with a speed of 700 rpm, and pressured up to 1000 psi to pack the C18 material into the capillary for ~15 cm long from the frit (packing time: ~1-2 h). The pressure in the chamber was released, the capillary was pulled up to be above the solvent in the LC vial, and the chamber was pressured to flow helium gas through the capillary until dry. The spot at 4.5 cm from the left side of the capillary was then fixed at the center (laser point) of a prewarmed (~10 min) laser puller (Sutter P-2000). The capillary was pulled once by 5 loops of the following parameters: HEAT 280, FIL 0, VEL 30, DEL 200, PULL 0. The resulted open tip size should be ~3 µm with a taper length of ~1.3 mm. It is optional to check whether the tip is open under microscope after it is immersed in $H_2O$ for a few seconds. The tip can be gently scratched against the surface area of ceramic scoring wafer to obtain a bigger opening (with a higher flow rate on LC). The resulted nano-column was equilibrated with 10 µL of 0.1%TFA in $H_2O$ at up to 400 bar with a flow rate at least 300 nL/min, and the portion that was not packed with C18 material was removed. Trypsin-digested BSA peptides (New England BioLabs Inc.) and HeLa Protein Digest Standard (Pierce) were used for routine quality control of nano-column, LC and mass spectrometry prior to sample analysis.

**Sample analysis by liquid chromatography-tandem mass spectrometry (LC-MS/MS)**

Samples were analyzed using methods adapted from previous studies[1,10]. In brief, an LC-MS/MS system consisted of an Easy-nLC 1200 coupled to a Fusion Orbitrap (Thermo Scientific) was used for peptide analysis. Peptide samples were maintained at 4 °C on sample tray in LC. Separation of SILAC samples was carried out on an in-house packed nano-column at room temperature with a mobile phase consisting of a linear gradient of A (0.1%TFA in $H_2O$) and B (80% acetonitrile in $H_2O$ containing 0.1% TFA) under the following conditions: 0 → 5 → 55 →

65 → 85 min, 2% → 8% → 35% → 100% → 100% B. The gradient for IsoTOP-ABPP sample was: 0 → 5 → 60 → 70 → 100 min, 0% → 0% → 45% → 100% → 100% B. The gradient for peptides from a pure protein was: 0 → 20 → 25 → 34 min, 0% → 40% → 100% → 100% B. The flow rate was 300 nL/min with the exception that 500 nL/min was used for the analysis of SCRN2 peptides from IsoTOP-ABPP sample preparation. The injection volume of all samples is 5 μL.

The voltage applied to the nano-LC electrospray ionization source was 2.4 kV. The temperature of ion transfer tube (ITC) was set at 275 °C. Spectra were collected in a data-dependent acquisition mode such that each scan cycle (3 seconds) involved a single high-resolution (120,000) full MS spectrum of parent ions (MS1 scan from *m/z* 400–1800) collected in the orbitrap. Parent ions assigned as peptide in charge states +2-6 with intensity higher than 5,000 were included for fragmentation. HCD-induced fragmentation (MS2) scans were recorded in ion trap. Dynamic exclusion was set as repeat count of 1 within exclusion time of 20 s. All other parameters were left as default values. For some peptide samples from IsoTOP-ABPP preparation and pure proteins, MS2 spectra were detected by orbitrap at a resolution of 30,000, dynamic exclusion was set as repeat count of 2 within exclusion time of 20 s. High quality of MS2 spectra of some IsoTOP-ABPP peptides were obtained by fragmenting paired parent ions with a mass difference of 6.0138 (difference between introduced heavy and light TEV tags) only.

**Peptide identification**

Each data file (in ".raw" format) was generated by the instrument (Xcalibur software). A derived file (in ".ms2" format) containing MS2 spectra for all fragmented parent ions were extracted using RawConverter (version 1.1.0.23) with monoisotopic selection (2015 released, publicly available at http://fields.scripps.edu/rawconv). Each ".ms2" file was searched using the ProLuCID algorithm against a reverse-concatenated, nonredundant database of the human proteome (UniProt release –03/26/2018) and filtered using DTASelect 2.0 within the Integrated Proteomics Pipeline (IP2) software. Cysteine residues were searched with a static modification for *S*-carbamidomethylation (+57.02146 Da). Methionine residues were searched with up to one differential modification for oxidation (+15.9949 Da). Peptides were required to have at least one tryptic terminus but an unlimited number of missed cleavages were allowed in the database search. For SILAC samples, each dataset was searched for both light and heavy isotopologues of the same peptide by specifying the mass shift of heavy residues as static modifications on lysine (+8.0142 Da) and arginine (+10.0082 Da) in a coupled 'heavy' search. The parent ion mass tolerance for a minimum envelope of three isotopic peaks was set to 50 ppm, the minimum peptide length was six residues, the false-positive rate was set at 1% or lower, and a minimum of two peptides of a protein must be detected in order to be advanced to the next step of analysis.

**Peptide and protein quantification**

Heavy and light parent ion chromatograms associated with successfully identified peptides were extracted and compared using in-house software (CIMAGE) as previously described[10]. At least one ion of a co-eluting heavy-light pair must be accurately identified from a fragmentation event that occurred within the retention time window (± 0.5 min) of parent ion elution. To ensure that the correct pair of peaks is quantified, chromatograms are extracted within a 10 ppm error tolerance of the theoretical *m/z* values, the signal-to-noise ratio must be > 2.5 and the 'co-elution correlation score' and 'envelope correlation score' must have $R^2$ values ≥ 0.8. In addition, peptides detected as 'singletons' where only the heavy ion of a peptide pair was identified, but for which passed all other filtering parameters, are given a default assigned ratio of '20', which is defined as any measured ratio that is ≥ 20.

**Determination of high-reactivity protein targets**

Custom R script, postCIMAGE, was used to process all .txt data files generated from CIMAGE. Briefly, protein ratios were then determined by the median peptide ratio derived from three or more unique quantified peptides, to further eliminate false positives and any stochastic variability in the data. Protein ratios that comply with these criteria from a single experiment were then averaged with ratios acquired from other replicates to provide final values. These ratios of probe **3** targets were aligned with that of probe **1** and probe **2** ("alkyl probe **1**" and "aryl probe **2**" previously) targets from previous work[1]. The aligned data are reported in **Supplementary Dataset 2** including each type of experiment in both cell lines with both probes. Single replicates for each type of dataset (only new data from probe **3**) are also included to show peptides identified per protein, their measured *m/z* values and median peptide ratios. Proteins that exhibited final ratios ≥ 5 in enrichment experiments and ≥ 3 in competition experiments were considered as the most robust targets of the probes. Note that infrequently detected proteins that failed to be quantified in both enrichment and competition experiments from a cell line were excluded as potential high-reactivity targets. Probe targets in cells (HEK293T and MDA-MB-231) represent data combined from ≥ 6 independent biological replicates, respectively, with each experiment performed with several independent batches of synthesized probes.

#### Functional annotation of protein targets by hydrazine probes

Protein targets (group 1: average enrichment ratio ≥ 5 and competition ratio ≥ 3, and group 2: average enrichment ratio ≥ 5 or competition ratio ≥ 3, respectively) were queried against the Drugbank database (Version 5.1.6) and fractionated into DrugBank and non-DrugBank proteins. Proteins were queried against PANTHER Classification Systems (Version 15.0) to assign functional keywords at the protein level and further classified into protein functional classes. Enzymes were classified into enzyme classes by PANTHER Protein Class. Protein targets with average enrichment ratio ≥ 5 or competition ratio ≥ 3 of probes **1**, **2** and **3** were compared with each other to generate Venn Diagram.

#### Phylogenetic analysis of protein targets

We performed phylogenetic analysis in order to visualize the distribution and overlapping of probed proteins with human targets. Phylogenetic trees were built for both Fe(II)/2OG enzymes and flavoproteins respectively. To collect all known human flavoproteins, we exhausted the UniProt protein sequence database by searching for the keywords “flavoprotein” and “flavin protein”. This identified 166 queries that are annotated as “reviewed human flavoproteins” according to UniProt description. After careful manual curation, 123 of them were considered as the most authentic ones and were kept for downstream analysis. A similar strategy was used for preparing all known human Fe(II)/2OG enzymes by leveraging InterPro database. After text mining for human Fe(II)/2OG enzymes in 56 highly related from InterPro, 131 queries were gathered and fed to manual curation, from which a final list of 68 most authentic ones were selected for downstream analysis. To build the trees, we first prepared multiple sequence alignments (MSAs) with the collected sequences using ClustalW[11]. Then, maximum likelihood trees were built with MAGAX[12]. Probed proteins were highlighted with dark blue and the visualization was achieved with Dendroscope[13].

#### Characterization of glyoxylyl modification in SCRN2 by isoTOP-ABPP

Previous study discovered glyoxylyl and pyruvoyl cofactors on Cys6 of SCRN3.[1] Protein sequence alignment of SCRN1/2/3 proteins identified similar ‘SCD’ motif between SCRN2 and SCRN3 (**Fig. 2b**). The labelling of recombinant SCRN2 and loss of labelling of its C12A mutant by both probe **2** and probe **3** show that Cys12 in SCRN2 behaves similarly with that of Cys6 in SCRN3, suggesting that SCRN2 may harbor similar functionalities with glyoxylyl and/or pyruvoyl in SCRN3. We initially tried to label the targeted peptide in SCRN2 by isoTOP-ABPP using both probe **2** and probe **3**, since SCRN2 is a target of both probes. We did not observe any MS1 and

MS2 spectra from probe **2**-treatment samples, however, trace MS2 spectra with parent ion matching theoretical probe **3**-labelled SCRN2 peptide were observed occasionally. Although these trace observations were not with high confidence, they gave us hope that SCRN2 may have functional PTMs. Because the targeted peptide (DCFVSVPPASAIPAVIFAK) of SCRN2 is 19 amino acid residues, 5 residues longer than targeted SCRN3 peptide (DTFVALPPATVDNR), and the 'C' was expected to be carbamidomethylated by iodoacetamide (IAA) during the isoTOP-ABPP protocol, we hypothesized that the resulted SCRN2 peptide is more hydrophobic than SCRN3 peptide. Chromatography of the SCRN2 peptide might be a reason why we did not get good MS1 and MS2 spectra. Therefore, we put in effort to optimize the LC method. Using a high flow method (500 nL/min) instead of a routine flow method (300 nL/min), we were able to get both MS1 and MS2 spectra for SCRN2 peptide harboring a glyoxylyl (**Supplementary Fig. 9-10**). However, we never observed any data supporting the presence of pyruvoyl on the same peptide. The SCRN2 peptide captured by probe **3** was prone to show its charge 3 state, although the doubly charged ion in MS1, $[M + 2H]^{2+}$, was obtained with a much lower intensity than that of charge 3 state (data not shown). This long peptide gave rich MS2 information in high-resolution for identification, notably, $y_{13}$ and its doubly charged ion showed up as the two highest ions due to the proline effect[14] which was seen in SCRN3 peptides[1]. Many b ions containing the isotopic tags indicated the site of probe modification and were used to verify the peptide modification as a glyoxylyl. Besides the classic b and y ion series from a peptide, many secondary b' and b'' ions derived from the fragmentations of $y_7$ and $y_{13}$ ions were observed in MS2 spectra with low mass errors in ppm, and thus served as additional confirmation of the peptide sequence.

#### Identification of probe 3-labelled peptides in NQO2 and KDM1A by isoTOP-ABPP

The mechanism-based inactivation of flavoproteins provided insight into the reaction and corresponding product for the transformation for probe **3**[15,16]. Our observation of the FAD-probe adducts in NQO2 (**Supplementary Fig. 6**) suggested the same chemical transformation (e.g. loss of hydrazine N-atoms and alkylation of the cofactor). This chemistry translates to a predicted mass shift for a peptide, however the site remains unknown. To determine where radical coupling occurs in the active site, we applied two strategies that were used to cross-validate one another: i) an IP2 search, ii) and an MS2 script match. The first strategy is to search the differential modifications (619.3587 for heavy TEV tag clicked with transformed probe, and 613.3449 for light TEV tag clicked with transformed probe) on each amino acid using IP2 software. In this method, we first searched against the FASTA containing NQO2 (or

KDM1A) only to identify the modified amino acid residue(s) using a narrowed search space. The second strategy is to match the peptide containing the modification with the experimental MS2 scans by an in-house developed script, the detailed description of which can be found in the following section. Upon identification of probe labelling sites, subsequent static modification searches using these sites were conducted against the database containing the human proteome to eliminate false positive peptides belonging to other proteins than the targeted protein.

**Identification of probe 2-labelled peptides in FTO by isoTOP-ABPP**

FTO was efficiently labelled by probe **2** but not probe **3**, however, it has no reported reactivity with hydrazine. Mutagenesis study shows that the labelling is active-site-directed since each iron ligand and 2OG binding residue important for catalysis is necessary for covalent capture (**Fig. 2c**). There are two unknowns to consider: i) the active site residue(s) that are labelled and ii) the structural changes of probe, both of which make the discovery and assignment of such peptides challenging. Because of these unknowns, existing peptide identification software, e.g. IP2, to computationally search and assign the labelled peptides are incompatible.

Therefore, we first investigated whether the hydrazine warhead forms the covalent bond with FTO by using an isotopically labelled probe (**Fig. 1a**). The experimental strategy is shown in **Supplementary Fig. 15**. Cells transfected with FTO were treated with probe **2** and probe **3**, respectively. SCRN3, with known labelling mechanisms with probe **2,** was used as positive control and was predicted to show two pairs (glyoxylyl and pyruvoyl peptides) of probe labelled peptides. A mass shift of 0.9970 between probe **2** and probe **3** treatments (**Supplementary Fig. 16-19**) was expected because the *N*-atoms of hydrazine are retained in probe capture, consistent with a polar capture mechanism of an electrophilic cofactor.

To find the pairs belonging to FTO, we extracted all the MS1 pairs in each sample using an in-house developed Java script which requires .MS1 file generated from .RAW file as input. Each monoisotopic precursor (charge states 2-3) in MS1 was searched for a possible isotopic pair partner with a mass difference of 6.0138 (difference between heavy and light TEV tags). The error tolerance of the two monoisotopic peaks was set at 5 ppm, and the relative intensity of the higher monoisotopic peak is required to be < 2 times of the lower peak. The tolerance of the retention time difference between the first appearances of the monoisotopic peaks in a pair was confined within 0.5 min. The two isotopic envelopes were required to be >75% similar with each other before being identified as a pair. All the MS1 pairs extracted from four samples (SCRN3-transfected cells treated with probe **2**, SCRN3-transfected cells treated with probe **3**, FTO-transfected cells treated with probe **2**, and FTO-transfected cells treated with probe **3**) are shown in **Supplementary Dataset 3**.

We next filtered the pairs to narrow down the pairs specific to SCRN3 and FTO. Pairs meeting the following criteria were kept for subsequent processing: 1) retention time within 20-45 min, 2) calculated parent mass > 1500 Da, and 3) the intensity of the pair > 1e6. Afterwards, background pairs present in both SCRN3 and FTO samples were removed and specific pairs to SCRN3 or FTO were kept to check the mass shift (0.9970 indicating the retention of $^{15}N$ in the labelled peptide, or 0.0000 Da indicating the loss of $^{15}N$ in the labelled peptide) between probe **2** and probe **3** samples. Two positive control pairs showed up in SCRN3 as shifted pairs. Interestingly, FTO gave three unshifted pairs and zero shifted pairs indicating the loss of $^{15}N$ (or hydrazine) in the FTO-probe **2** reaction.

To reveal the identities of the top three promising peptides labelled by probe **2**, we developed another Java script to search the plausible MS2 spectra of a given peptide from .MS2 file. This script permits only one peptide per search and virtually fragments the input peptide into theoretical b and y ions which are then matched with every MS2 scan in the experimental data. MS2 ions with mass errors of 0.2 Da for low-resolution data or 20 ppm for high-resolution data are considered as matched ions, the minimum number of which is set at 3-5 to eliminate false positives. The intensities of the matched MS2 ions are required to be > 5% of that of base peak in the same spectrum. For the three FTO pairs, we had to assume that each pair was a resultant of an FTO peptide, and thus had to do trial-and-error cycles for all possible FTO peptides after virtual trypsin digestion (http://prospector.ucsf.edu/prospector/mshome.htm). We have further optimized the script to be capable of automatically matching FTO peptides with experimental MS2 scans and ranking the peptides by matching scores. This workflow identified two labelled peptides of FTO. Detailed elucidation of the heavy and light MS2 spectra gave the sites of probe labelling and the mass differences between unmodified and labelled peptides for structural explanation (**Supplementary Fig. 21-22**).

**Click chemistry and Imaging of probe 3 labelled proteome in HEK293T cells**

HEK293T cells (ATCC CRL-11268) were cultured in DMEM high glucose with GlutaMAX (Thermo Scientific, cat# 10569044), supplemented with 10% heat-inactivated characterized FBS (HyClone, cat# SH30071.01) and 100 U/mL of penicillin/streptomycin (Corning, cat# 30-002-CI). Cells were plated onto poly-D-Lysine-coated (Sigma, cat# P7405) 22 x 22 mm coverslips (one per well) at 400,000 cells/well in 6-well plates. The next day, cells were washed 3x with 2 mL serum-free starve media (culture media without FBS), and then treated in a volume of 500 uL/well in serum-free starve media supplemented with 10 mM HEPES sodium salt (Sigma, cat# H7006) pH 7.5 as follows: (1) negative control (probe without CuAAC reaction), (2) probe with CuAAC reaction, and (3) competitor + probe with CuAAC reaction. Competitor was added to wells in group 3 in a volume of 12.5 uL as a pretreatment for 15 min (final concentration: 10 mM) and then probe was added to wells in groups 2 and 3 in a volume of 5 uL for 15 min (final concentration: 1 mM). Following treatment, media was aspirated and cells were washed 2x with 1X Dulbecco's PBS plus calcium and magnesium (Corning, cat# 20030CV), fixed with 4% PFA in PBS pH 7.4 (Electron Microscopy Sciences, catalog# 157-4) for 20 min, and then washed 2x with PBS for 10 min on a shaker at room temperature.

For immunofluorescent detection and CuAAC functionalization in fixed cells, each incubation was performed at room temperature on a shaker, with 2x PBS washes for 10 min between each processing step. Cells were first incubated with WGA-488 (Thermo, catalog# W112961) at 5 ug/mL in PBS for 10 min to label plasma membranes, and then permeabilized with 0.1% Triton X-100 (Sigma, cat# T8787) in PBS for 2 min. CuAAC reaction was performed as per previous protocol[17] on the fixed and permeabilized cells. Briefly, reagents were prepared as a premixed solution in the order listed at final concentrations as follows: (1) negative control wells received 50 uM Azide-Fluor 545 (Thermo Scientific cat#760757) in PBS minus click reagents; (2) probe alone or (3) competitor + probe plus click reagents received 50 uM Azide-Fluor 545, a premixed solution of 50 mM THPTA (Sigma, cat# 762342) and 20 mM copper (II) sulfate (Sigma, cat# 451657) which is then diluted to final concentrations of 0.5 mM THPTA and 0.1 mM copper (II) sulfate in the wells, 5 mM aminoguanidine (Sigma, cat# 396494) and 5 mM L-ascorbic acid (Sigma, cat# A5960) in PBS. The CuAAC reaction was carried out for 1 h at room temperature. To visualize nuclei, cells were incubated with 300 nM DAPI (Sigma, cat# D9542) for 5 min, and then mounted to Superfrost Plus slides (Fisher, cat# 12-550-15) with 50 uL of Fluoromount-G (Southern Biotech, cat# 0100-01). Images were acquired on a confocal Nikon C2 microscope at 20X magnification for quantification and 63X oil for representative images.

#### Tissue imaging by probe 3 for proteome labelling in mouse

Mouse lung tissues were fixed in 10% neutral buffered formalin (Fisher Scientific 23-245-685) for 16 hours and dehydrated in a series of ethanol washes up to 100%. Samples were paraffin embedded and sectioned at 4-μm thickness. Tissue sections were prepared for probe labelling by removing paraffin with xylene and rehydrating the samples in 100% ethanol for 3 minutes followed by 95% and 80% ethanol for 1 minute each. Samples were then rinsed in distilled water. Antigen retrieval was performed in a pressure cooker using citrate buffer (Electron

Microscopy Sciences 62706-10). The samples were incubated with the 333 μM probe **3** or probe **3** plus 3.3 mM drug **4** resuspended in phosphate buffered saline (PBS) for one hour at room temperature. The slide was then washed in PBS. The click reaction was performed by adding $CuSO_4$ (2 mM), TCEP (1 mM) and rhodamine-azide (100 μM) and incubated at room temperature for 1 h. Slides were counterstained using antibodies against acetylated tubulin (1:200, sc23950 from Santa Cruz) and with DAPI using Fluoro-Gel II with DAPI (EMS Catalog #: 17985-50). Detection was by Alexa 488 nm conjugated anti-mouse secondary antibodies (Molecular Probes). Slides were analyzed on a Leica DMI6000B inverted microscope.

#### Synthesis of isotopic protease-cleavable biotin-azide peptide tags

Peptide tags (see structures below) for site of labelling experiments were synthesized as previously described[10] with the following exceptions. Heavy and light peptide products were purified by RP-HPLC gradient: 0–70% 90:10 *v/v* MeCN/$H_2O$ in $H_2O$, 0.05 *v*-% TFA in 35 min; flow rate: 5 mL/min using a Phenomenex Jupiter 5 μm C18 300 Å (150 × 10.0 mm) column.

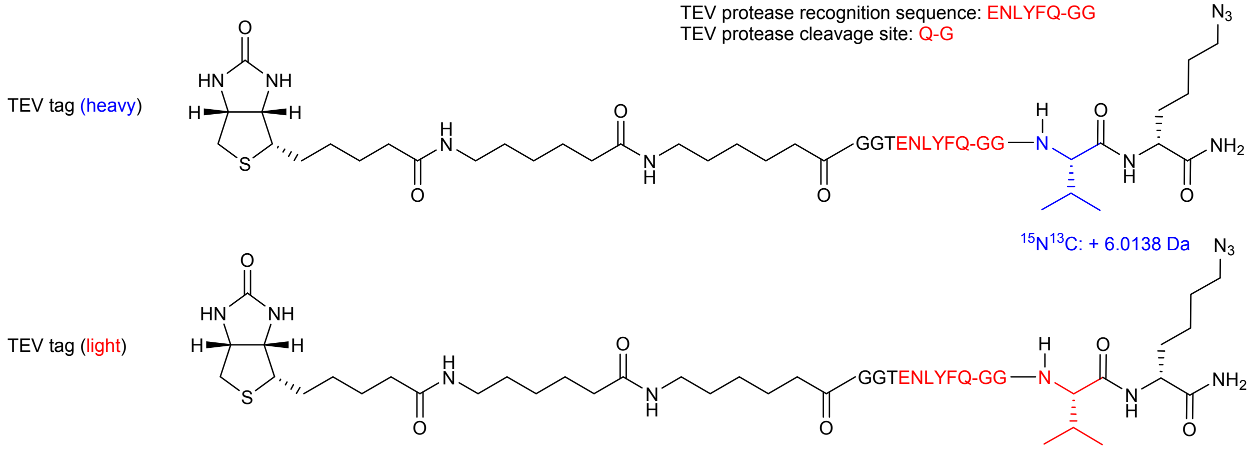

### 4. Synthetic Methods

#### Materials

All chemicals were obtained from commercial suppliers and were used without further purification. All chemical transformations requiring inert atmospheric conditions were carried out under high purity (100%) argon gas. Flash chromatography was performed on a Teledyne ISCO CombiFlash Rf automated purification system (Lincoln, NE, USA). Separation was carried out on SiliaSepFlash silica cartridges (40 - 63 μm, 60 Å) or SiliaSepFlash C18 monomeric cartridges (25 μm, 90 Å) while the products were monitored by a UV detector at 254 nm and/or evaporative light scattering detector (ELSD). NMR spectra were recorded at room temperature on a Bruker UNI-500 (500 MHz), DMX-360 (360 MHz) or UNI-400 (400 MHz) spectrometer (Bruker BioSpin, Billerica, MA, USA). Peak multiplicities are reported as follows: *s* = singlet, *bs* = broad singlet, *d* = doublet, *t* = triplet, *q* = quartet, *m* = multiplet, *bm* = broad multiplet. Chemical shifts of $^1$H NMR spectra were referenced to residual solvent peaks of $CDCl_3$ ($\delta$ 7.26 ppm), DMSO-$d_6$ ($\delta$ 2.50 ppm), $CD_3OD$ ($\delta$ 3.31 ppm) or $D_2O$ ($\delta$ 4.79 ppm). Chemical shifts of $^{13}$C NMR spectra were referenced to residual solvent peaks of $CDCl_3$ ($\delta$ 77.16 ppm), DMSO-$d_6$ ($\delta$ 39.52 ppm) or $CD_3OD$ ($\delta$ 49.00 ppm). For polar compounds like hydrazine salts, reactions were monitored by a Waters Acquity UPLC equipped with a single quadruple MS (Milford, MA, USA). LC chromatograms were obtained from an Acquity UPLC system equipped with a C18 column (1.7 μm, 2.1 x 50 mm) and a UV detector (wavelength fixed at 254 nm unless otherwise specified). The mobile phase was composed of 100% $H_2O$ containing 0.1% formic acid (A) and 100% MeCN containing 0.1% formic acid (B) at a flow rate of 0.3 mL/min with the following gradient: 0-4 min, 5%-95% B; 4-5 min, 95% B. MS data in both positive and negative modes were acquired by the aforementioned MS detector equipped with an electrospray ionization (ESI) source. TLC analysis was performed on silica gel plates (60 Å porosity, 250 μm thickness) which were then developed by hexane/EtOAc or DCM/MeOH as mobile phase and visualized by UV light (see details for each compound below). High-resolution mass spectra (HRMS) were obtained from either a Waters LCT Premier XE LC-ESI-TOF-MS (Milford, MA, USA) or a Waters GCT Premier, GC-EI-TOF-MS (Milford, MA, USA). Purities of final probes were determined by UPLC as >95% prior to use for experiments.

**Synthesis of phenelzine probe (3)**

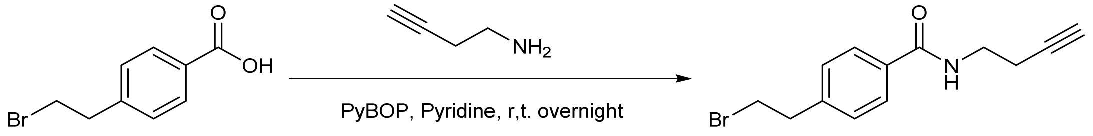

11 12

**4-(2-bromoethyl)-*N*-(but-3-yn-1-yl)benzamide (12).** But-3-yn-1-amine (0.152 g, 2.20 mmol, 1.10 equiv.) was added to a solution of 4-(2-bromoethyl)benzoic acid **11** (0.430 g, 2.00 mmol, 1.00 equiv.) and benzotriazol-1-yl-oxytripyrrolidinophosphonium hexafluorophosphate (PyBOP; 1.14 g, 2.20 mmol, 1.10 equiv.) in pyridine (10.0 mL, 0.200 M) under argon. The reaction was stirred at room temperature overnight. After disappearance of starting material **11** by UPLC/MS, the reaction was acidified with 3.0 M hydrochloric acid at 0 °C until a white precipitate was observed. The mixture was extracted with ethyl acetate (3 x 75.0 mL) and the combined organic layers were dried over anhydrous $Na_2SO_4$ and concentrated under reduced pressure. The remaining residue was purified on a silica column by flash chromatography using hexane (A) and ethyl acetate (B) as mobile phases with an 15 min gradient from 5%-70% B to afford the desired bromoethyl compound **12** as an off-white powder (0.409 g, 1.46 mmol, 73%): $^1$H NMR (360 MHz, $CDCl_3$) δ 7.74 (d, *J* = 8.0 Hz, 2H), 7.29 (d, *J* = 8.0 Hz, 2H), 6.48 (s, 1H), 3.81 – 3.56 (m, 4H), 3.40 – 3.03 (m, 2H), 2.54 – 2.50 (m, 2H), 2.05 (s, 1H). $^{13}$C NMR (90.5 MHz, $CDCl_3$) δ 167.2, 142.6, 133.1, 128.9, 127.3, 81.6, 70.2, 38.9, 38.4, 32.3, 19.5. MS (ESI$^+$) *m/z* calc'd for $C_{13}H_{14}NOBr$ $[M+H]^+$: 280.0, found 280.0.

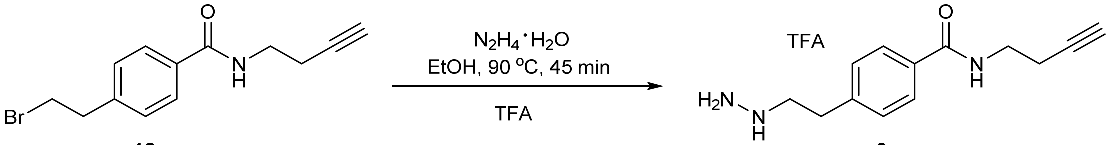

**12** **3**

***N*-(but-3-yn-1-yl)-4-(2-hydrazineylethyl)benzamide trifluoroacetate (3).** Hydrazine monohydrate (98%, 0.495 mL, 0.511 g, 10.0 mmol, 10.0 equiv.) was added dropwise to a stirring solution of bromoethyl benzamide **12** (0.280 g, 1.00 mmol, 1.00 equiv.) in ethanol (5.00 mL, 0.500 M) under argon, and the resulting mixture was stirred for 45 min at 90 °C. After disappearance of starting material **12** by UPLC/MS, the reaction mixture was cooled down to room temperature and acidified with 3 M HCl dropwise until pH < 2. The resulting solution was concentrated under reduced pressure to give the crude hydrazine hydrochloride product as a white solid. The crude product was then purified on a C18 column by flash chromatography using $H_2O$ containing 0.1% trifluoroacetic acid (A) and MeOH containing 0.1% trifluoroacetic acid (B) as mobile phases with a 10 min gradient from 0%-45% B. Fractions containing the product were combined, concentrated to remove organic solvents, and lyophilized to afford the desired product as a white solid (0.204 mg, 0.621 mmol, 62%): $^1$H NMR (500 MHz, $D_2O$) δ 7.77 (d, *J* = 8.5 Hz, 2H), 7.45 (d, *J* = 8.5 Hz, 2H), 3.57 (t, *J* = 6.5 Hz, 2H), 3.46 (t, *J* = 6.5 Hz, 2H), 3.11 (t, *J* = 7.5 Hz, 2H), 2.56 (td, *J* = 6.5, 2.5 Hz, 2H), 2.40 (t, *J* = 2.5 Hz, 1H). $^{13}$C NMR (125 MHz, $D_2O$) δ 170.7, 140.9, 132.5, 129.1, 127.7, 82.5, 70.4, 51.5, 38.4, 30.6, 18.5. Purity: 99.2% determined by UPLC. HRMS (ESI$^+$) *m/z* calc'd for $C_{13}H_{18}N_3O$ $[M+H]^+$: 232.1372, found 232.1369 (error: -1.3 ppm).

### Synthesis of $^{15}N_2$-phenelzine probe ($^{15}N_2$-3)

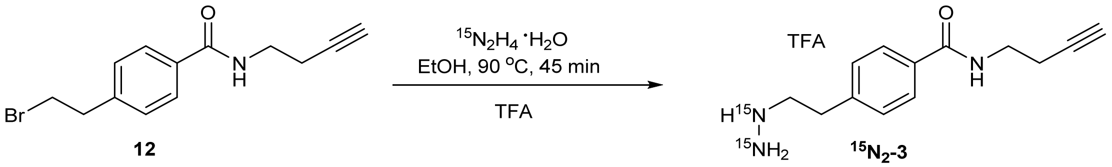

$^{15}N_2$-Hydrazine monohydrate (98%+ ATOM, 26.0 mg, 0.500 mmol, 2.50 equiv, Sigma-Aldrich, Catalog Number: 492787) was added dropwise to a stirring solution of bromoethyl benzamide **12** (56.0 g, 0.200 mmol, 1.00 equiv.) in ethanol (0.400 mL, 0.500 M) under Argon, and the resulting mixture was stirred for 30 min at 90 °C. After disappearance of starting material **12** by UPLC/MS, the reaction mixture was cooled down to room temperature and acidified with 3M HCl dropwise until pH < 2. The resulting solution was concentrated under reduced pressure to give the crude hydrazine hydrochloride product as a white solid. The crude product was then purified on a C18 column by flash chromatography using $H_2O$ containing 0.1% trifluoroacetic acid (A) and MeOH containing 0.1% trifluoroacetic acid (B) as mobile phases with a 10 min gradient from 0%-45% B. Fractions containing the product were combined, concentrated to remove organic solvents, and lyophilized to afford the desired product as a white solid (27.5 mg, 0.118 mmol, 59%): $^1$H NMR (400 MHz, $D_2O$) δ 7.65 (d, $J$ = 8.0 Hz, 2H), 7.35 (d, $J$ = 8.4 Hz, 2H), 3.45 (t, $J$ = 6.4 Hz, 2H), 3.35 (td, $J$ = 7.6, 2.0 Hz, 2H), 3.00 (td, $J$ = 7.6, 2.8 Hz, 2H), 2.45 (td, $J$ = 6.8, 2.8 Hz, 2H), 2.28 (t, $J$ = 2.8 Hz, 1H). $^{13}$C NMR (101 MHz, $D_2O$) δ 170.7, 140.8, 132.4, 129.1, 127.6, 82.4, 70.4, 51.5, 51.5, 38.4, 30.5, 18.4. $^{15}$N NMR (61 MHz, $D_2O$) δ 67.77, 67.68, 63.11, 63.03. Purity: 99.0% determined by UPLC.HRMS (ESI$^+$) *m/z* calc'd for $C_{13}H_{18}N_3O$ $[M+H]^+$: 234.1385, found 234.1380 (error: -2.1 ppm).

#### Synthesis of $^{15}$*N*-phenylhydrazine probe ($^{15}$N-2)

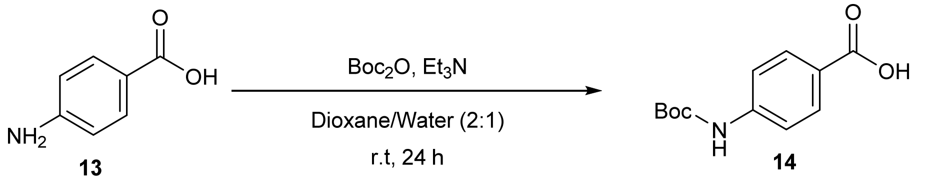

**4-((*tert*-butoxycarbonyl)amino)benzoic acid (14).** Triethylamine ($Et_3N$; 2.05 g, 20.0 mmol, 2.00 equiv.) was added to a stirring solution of 4-aminobenzoic acid **13** (1.37 g, 10.0 mmol, 1.00 equiv.) in dioxane/water (2:1, 30.0 mL, 0.333 M) and the mixture was stirred for 5 min at room temperature. Di-*tert*-butyl dicarbonate ($Boc_2O$; 4.43 g, 20.0 mol, 2.00 equiv.) was then added and the reaction was stirred for 24 h at room temperature. The reaction progress was monitored by UPLC/MS. Upon completion, the reaction mixture was concentrated under reduced pressure and then acidified with 3 M hydrochloric acid. The resulting slurry was filtered and washed with water (2 x 20.0 mL) and recrystallized from hot methanol to yield 4-((*tert*-butoxycarbonyl)amino)benzoic acid **14** as white solid (2.16 g, 9.10 mmol, 91% yield) without further purification.

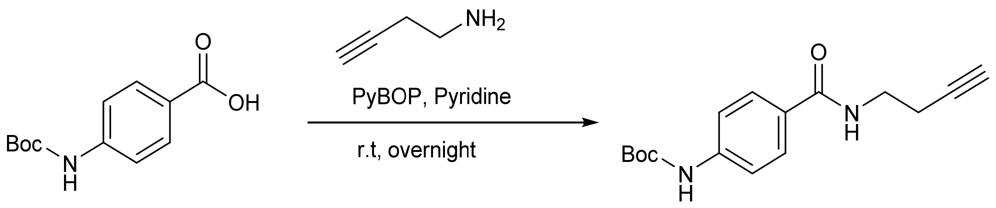

**14** **15**

***tert*-Butyl (4-(but-3-yn-1-ylcarbamoyl)phenyl)carbamate (15).** But-3-yn-1-amine (76.0 mg, 1.10 mmol, 1.10 equiv.) was added to a solution of 4-((*tert*-butoxycarbonyl)amino)benzoic acid **14** (0.260 g, 1.00 mmol, 1.00 equiv.) and benzotriazol-1-yl-oxytripyrrolidinophosphonium hexafluorophosphate (PyBOP; 0.572 g, 1.10 mmol, 1.10 equiv.) in anhydrous pyridine (5.00 mL, 0.200 M) under argon. The reaction was stirred at room temperature overnight. After disappearance of starting material **14** by UPLC/MS, the reaction was acidified with 3 M hydrochloric acid at 0 °C until a white precipitate was observed. The mixture was extracted with ethyl acetate (3 x 15.0 mL) and the combined organic layers were dried over anhydrous $Na_2SO_4$ and concentrated under reduced pressure. The remaining residue was purified on a silica column by flash chromatography using hexane (A) and ethyl acetate (B) as mobile phases with a 25 min gradient from 0%-50% B to afford the desired compound **15** as an off-white powder (0.242 g, 0.839 mmol, 84%): $^1$H NMR (500 MHz, $CDCl_3$) δ 7.71 (d, *J* = 8.50 Hz, 2H), 7.43 (d, *J* = 8.50 Hz, 2H), 6.91 (s, 1H), 6.54 (s,1H), 3.59 (q, *J* = 6.50 Hz, 2H), 2.50 (td, *J* = 6.50, 3.00 Hz, 2H), 2.03 (s, 1H), 1.51 (s, 9H). $^{13}$C NMR (126 MHz, $CDCl_3$) δ: 167.09, 152.46, 141.64, 128.41, 128.07, 117.81, 81.61, 81.09, 70.21, 38.41, 28.30, 19.52. MS (ESI$^+$) *m/z* calc'd for $C_{16}H_{21}N_2O_3$ $[M+H]^+$: 289.15, found 289.15.

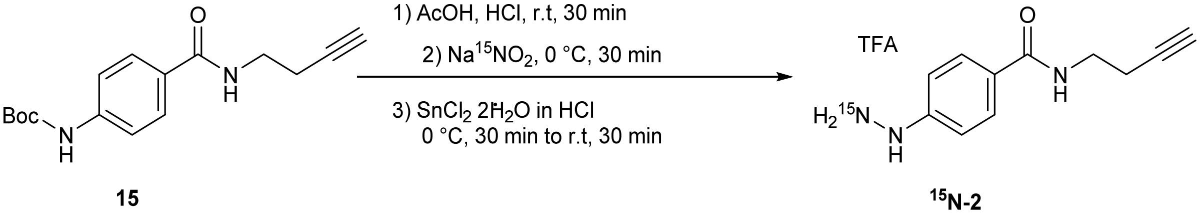

**15** **$^{15}$N-2**

***N*-(but-3-yn-1-yl)-4-(hydrazineyl-$^{15}$*N*)benzamide trifluoroacetic acid ($^{15}$N-2).** Concentrated hydrochloric acid (36% HCl; 0.500 mL, 1.00 M) was added dropwise to a stirring solution of *tert*-

butyl(4-(but-3-yn-1-ylcarbamoyl)phenyl)carbamate **15** (0.143 g, 0.500 mmol, 1.00 equiv.) in acetic acid (AcOH; 0.250 mL, 2.00 M) and the resulting mixture was stirred for 2 hours at room temperature. This solution was then cooled to 0 °C before a solution of $Na^{15}NO_2$ (41.9 mg, 0.600 mmol, 1.20 equiv.) in $H_2O$ (0.600 mL, 1.00 M) was dropwise added. The solution was stirred at 0 °C for 30 minutes before a solution of $SnCl_2{\cdot}2H_2O$ (0.248 g, 1.10 mmol, 2.20 equiv.) in concentrated HCl (0.550 mL, 2.00 M) was dropwise added. The resulting inhomogeneous solution was stirred for 30 minutes at 0 °C then allowed to warm to room temperature with stirring for 30 minutes. After disappearance of starting material **15** by UPLC/MS, the reaction mixture was concentrated under reduced pressure to give the crude hydrazine hydrochloride product as a white solid. The crude product was then purified on a C18 column by flash chromatography using $H_2O$ containing 0.1% trifluoroacetic acid (A) and MeOH containing 0.1% trifluoroacetic acid (B) as mobile phases with a 10 min gradient from 0%-40% B. Fractions containing the product were combined, concentrated to remove organic solvents, and lyophilized to afford the desired product as a pale-yellow solid (0.101 g, 0.335 mmol, 67%): $^1$H NMR (500 MHz, $D_2O$) δ 7.82 (d, *J* = 8.2 Hz, 2H), 7.10 (d, *J* = 8.1 Hz, 2H), 3.58 (t, *J* = 6.4 Hz, 2H), 2.61 – 2.53 (m, 2H), 2.40 (s, 1H). $^{13}$C NMR (126 MHz, $D_2O$) δ 170.2, 147.1, 128.9, 127.9, 114.1, 82.5, 70.4, 38.4, 18.5. $^{15}$N NMR (41 MHz, $D_2O$) δ 59.62. Purity: 98.3% determined by UPLC. HRMS (ESI$^+$) *m/z* calc'd for $C_{11}H_{14}{}^{15}N_1N_2O_1$ $[M+H]^+$: 205.1102, found 205.1112 (error: 4.9 ppm).

**Synthesis of NRH**

Under argon atmosphere, nicotinamide riboside (NR) (50 mg, 0.12 mmol) was added to 3 mL of 50 mM potassium phosphate (pH=8.5) in a round-bottom flask at 0 °C. After stirring for 5 min, sodium dithionite ($Na_2S_2O_4$) powder (209 mg, 1.2 mmol, 10 equiv.) was added, the mixture was stirred for 30 min at 0 °C. Conversion was confirmed by LC-MS. The crude mixture was subjected to flash chromatography (water) on a C18 column (SiliCycle Inc.) to give reduced NRH with a strong absorbance at 340 nm. Fractions were combined and dried under vacuum to yield a pale-yellow solid which was stored in -80 °C prior to use[18].

### NMR Spectra of hydrazine probes

$^{1}H$ NMR (500 MHz, $D_2O$)

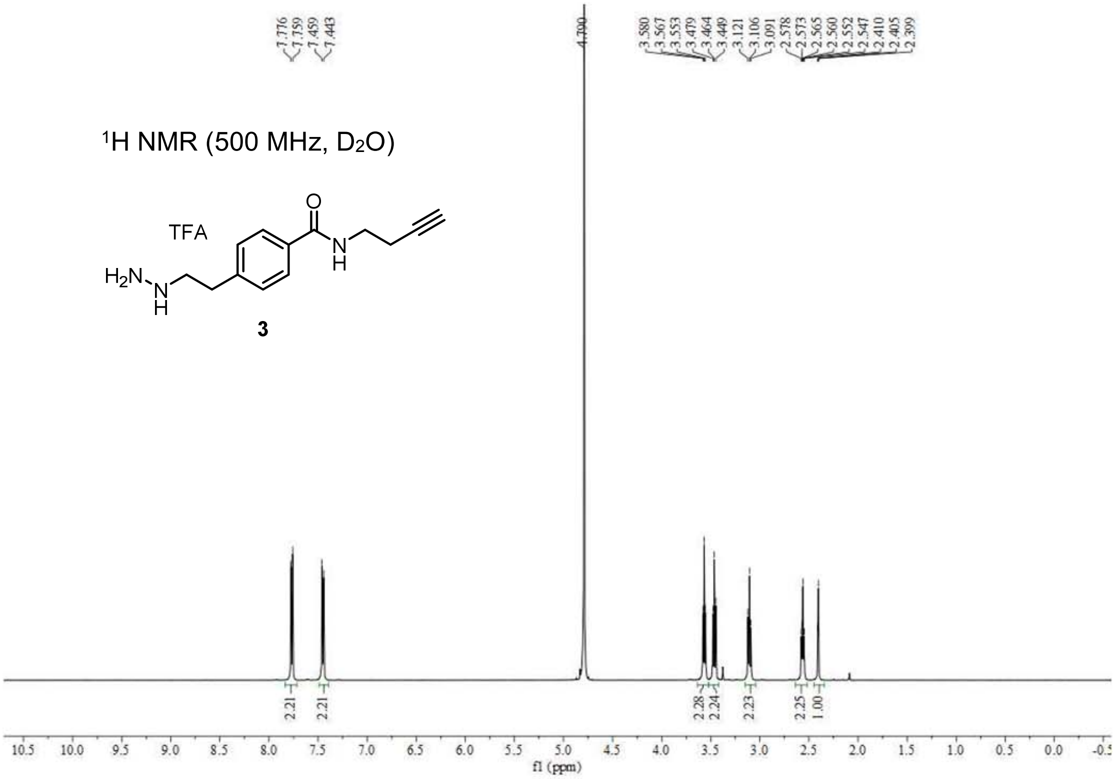

3

$^{13}C$ NMR (125 MHz, $D_2O$)

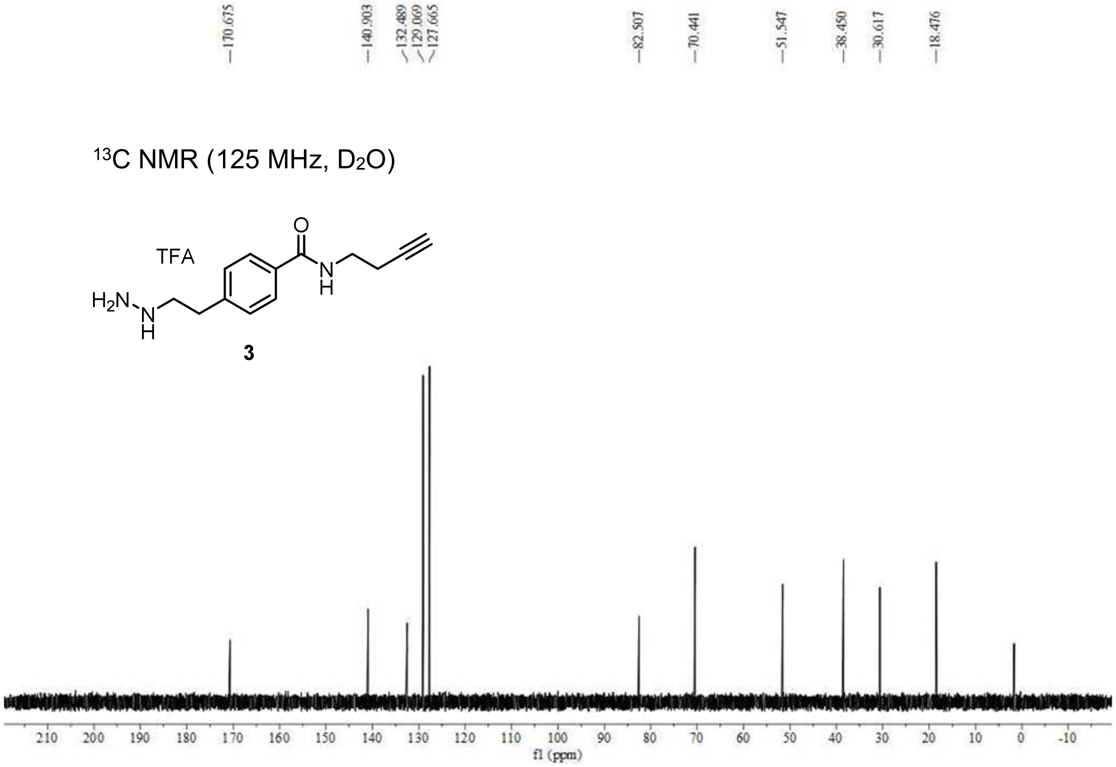

3

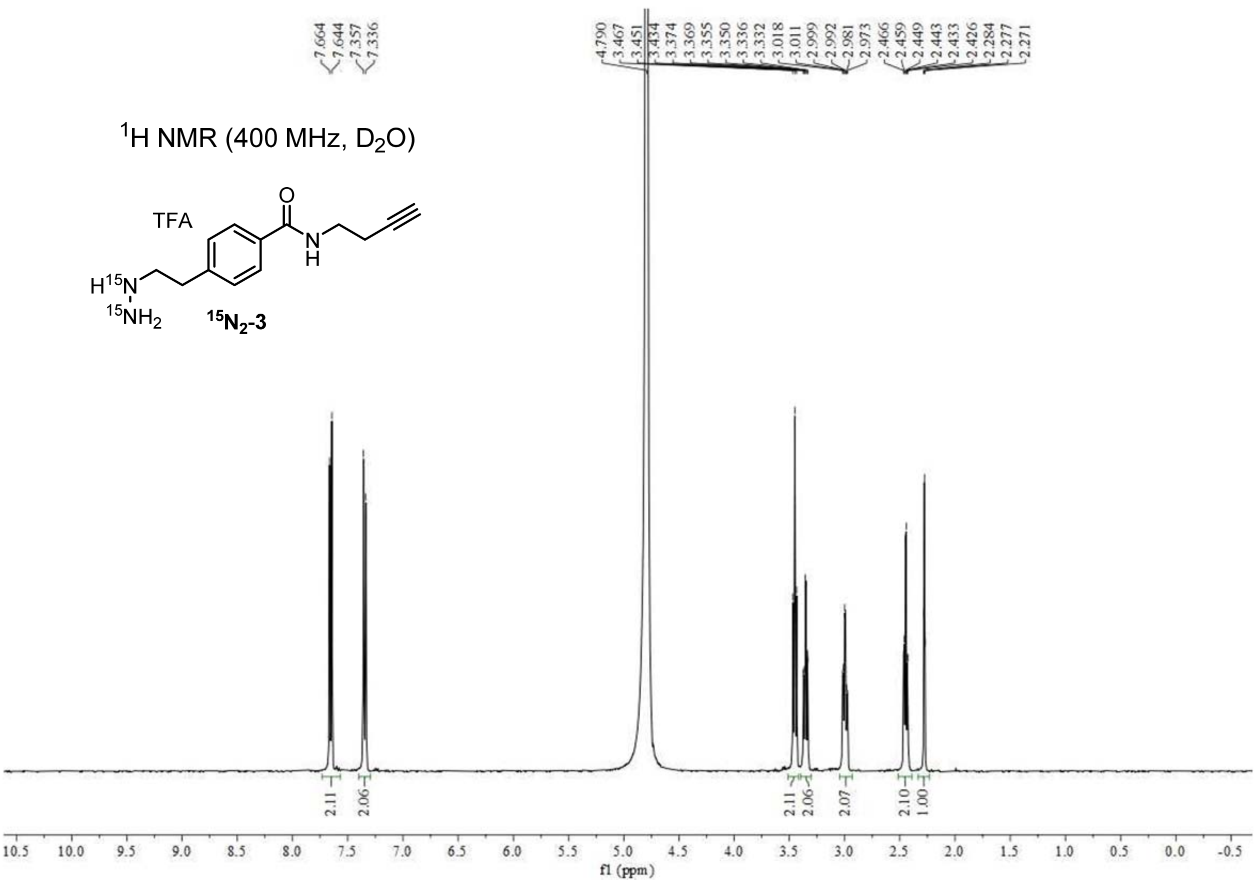

$^1$H NMR (400 MHz, $D_2O$)
TFA
$^{15}N_2$-3
f1 (ppm)

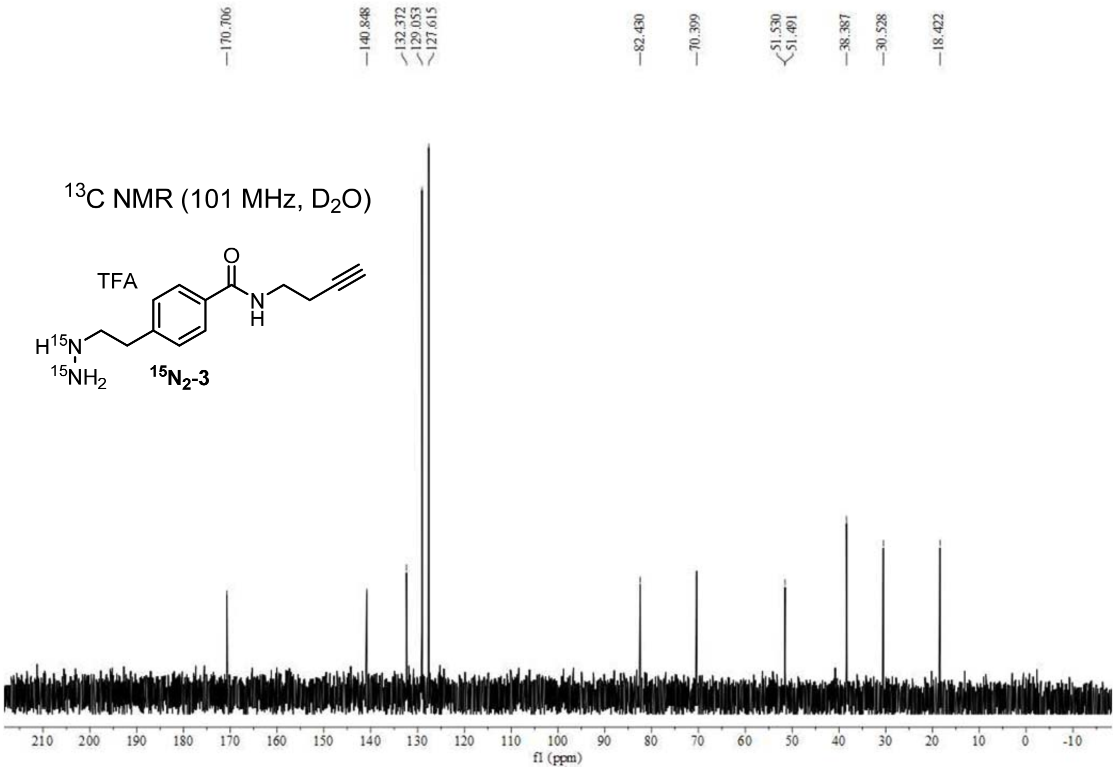
170.706
140.848
132.372
129.053
127.615
82.430
70.399
51.530
51.491
38.387
30.528
18.422
$^{13}$C NMR (101 MHz, $D_2O$)
TFA
O
N
H
H$^{15}$N
$^{15}$NH$_2$
$^{15}$N$_2$-3
210 200 190 180 170 160 150 140 130 120 110 100 90 80 70 60 50 40 30 20 10 0 -10
f1 (ppm)

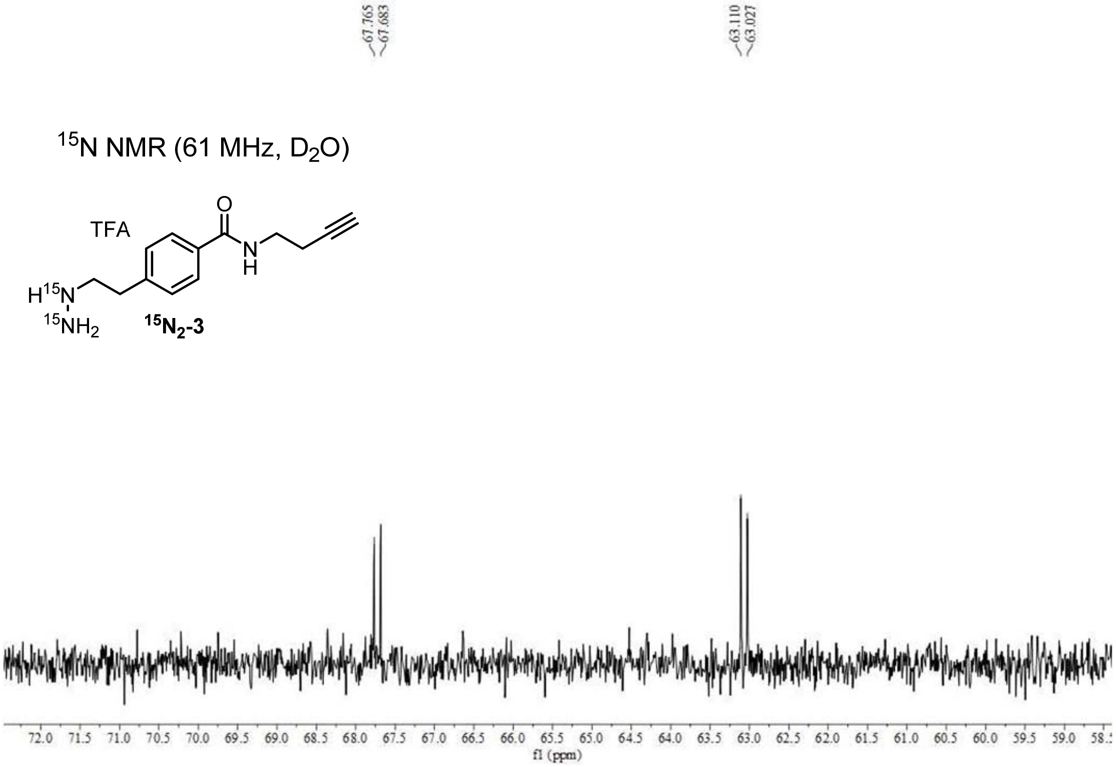
67.765
67.683
63.110
63.027
$^{15}$N NMR (61 MHz, $D_2O$)
TFA
O
N
H
H$^{15}$N
$^{15}$NH$_2$
$^{15}$N$_2$-3
72.0 71.5 71.0 70.5 70.0 69.5 69.0 68.5 68.0 67.5 67.0 66.5 66.0 65.5 65.0 64.5 64.0 63.5 63.0 62.5 62.0 61.5 61.0 60.5 60.0 59.5 59.0
f1 (ppm)

$^{1}H$ NMR (400 MHz, $D_2O$)

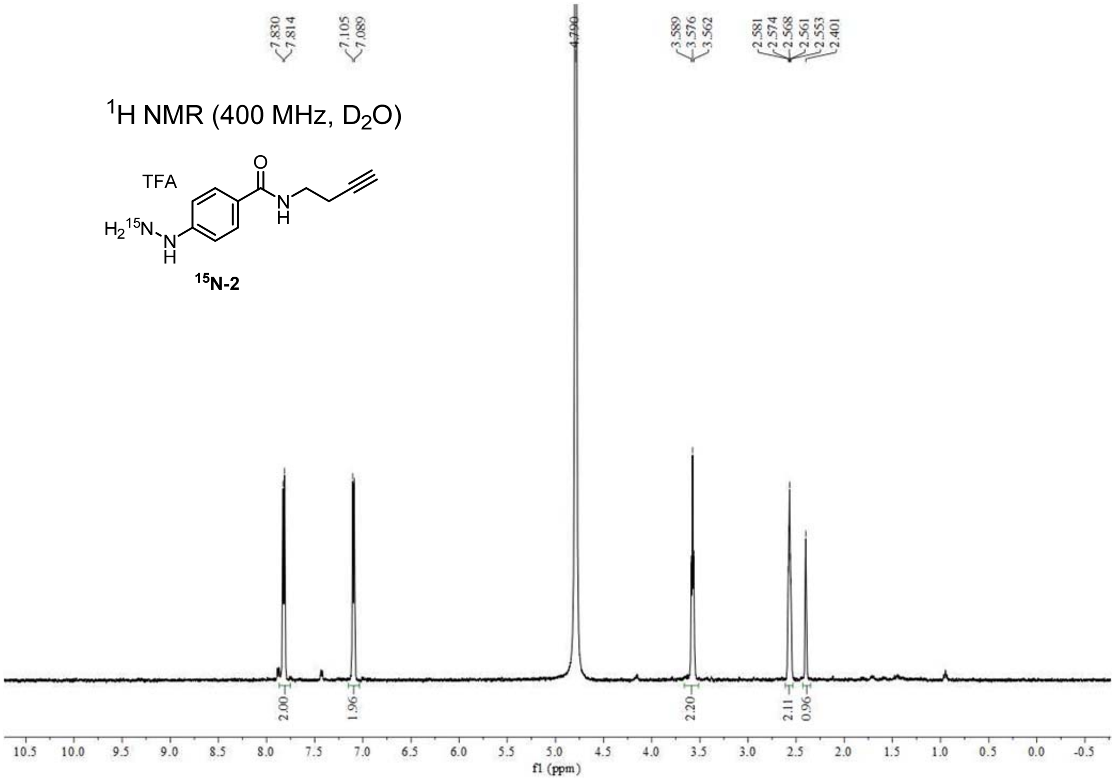

$^{13}C$ NMR (101 MHz, $D_2O$)

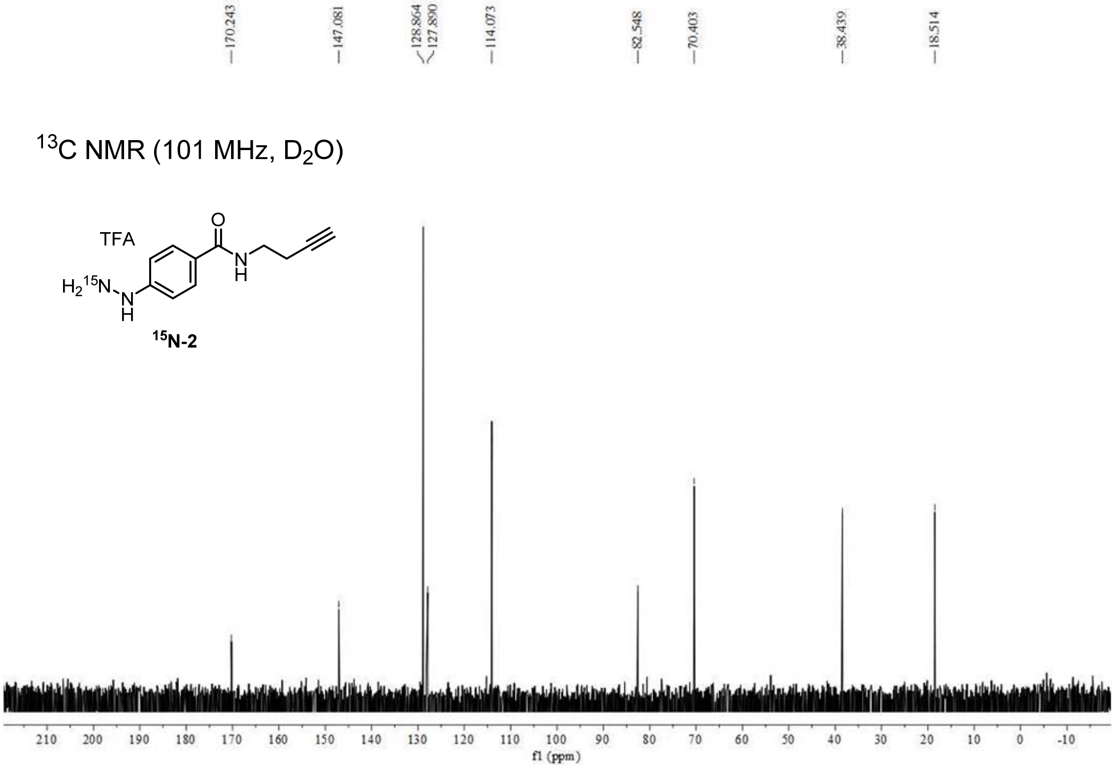

59.617

$^{15}N$ NMR (61 MHz, $D_2O$)

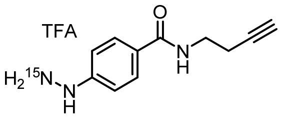

**$^{15}$N-2**

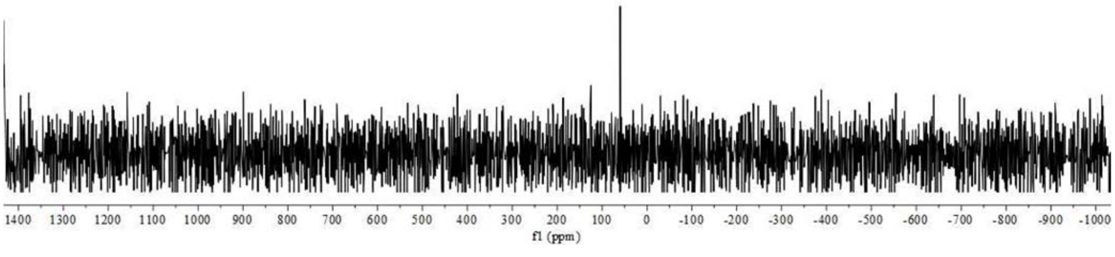
